# Combinatorial prediction of therapeutic perturbations using causally-inspired neural networks

**DOI:** 10.1101/2024.01.03.573985

**Authors:** Guadalupe Gonzalez, Xiang Lin, Isuru Herath, Kirill Veselkov, Michael Bronstein, Marinka Zitnik

## Abstract

Phenotype-driven approaches identify disease-counteracting compounds by analyzing the phenotypic signatures that distinguish diseased from healthy states. Here, we introduce PDGrapher, a causally inspired graph neural network (GNN) model that predicts combinatorial perturbagens (sets of therapeutic targets) capable of reversing disease phenotypes. Unlike methods that learn how perturbations alter phenotypes, PDGrapher solves the inverse problem of directly predicting the perturbagens needed to achieve a desired response by embedding disease cell states into networks, learning a latent representation of these states, and identifying optimal combinatorial perturbations. In experiments in nine cell lines with chemical perturbations, PDGrapher identifies effective perturbagens in more testing samples than competing methods. It also demonstrates competitive performance on ten genetic perturbation datasets. An advantage of PDGrapher is its direct prediction, in contrast to the indirect and computationally intensive approach traditionally used in phenotype-driven models. This approach accelerates training by up to 25 times compared to existing methods, providing a fast approach for identifying therapeutic perturbations and advancing phenotype-driven drug discovery.

## Main

Target-driven drug discovery, which has been the dominant approach since the 1990s, focuses on designing highly specific compounds to act against targets, such as proteins or enzymes, that are implicated in disease, often through genetic evidence [1–3]. An example of target-driven drug discovery is the development of small-molecule kinase inhibitors like imatinib. Imatinib stops the progression of chronic myeloid leukemia (CML) by inhibiting BCR-ABL tyrosine kinase, a mutated protein involved in uncontrolled proliferation of leukocytes in patients with CML [4]. Other notable examples include monoclonal antibodies such as trastuzumab, which specifically targets the HER2 receptor, a protein overexpressed in certain types of breast cancer. Trastuzumab inhibits cell proliferation while engaging the body’s immune system to initiate anti-cancer response [5]. These examples illustrate the success of target-driven drug discovery, yet the past decade has seen a revival of phenotype-driven approaches. This shift has been fueled by the observation that many first-in-class drugs approved by the US Food and Drug Administration (FDA) between 1999 and 2008 were discovered without a drug target hypothesis [6]. Instead of the “one drug, one gene, one disease” model of target-driven approaches, phenotype-driven drug discovery focuses on identifying compounds or, more broadly, perturbagens—combinations of therapeutic targets—that reverse disease phenotypes as measured by assays without predefined targets [1, 7]. Ivacaftor illustrates how these approaches can intersect. Although ivacaftor was developed through a target-driven strategy to modulate the cystic fibrosis transmembrane conductance regulator protein in individuals with specific mutations, its development relied on phenotypic assays to confirm functional improvements, such as increased chloride transport [8–10].

Phenotype-driven drug discovery has been bolstered by the advent of chemical and genetic libraries such as the Connectivity Map (CMap [11]) and the Library of Integrated Network-based Cellular Signatures (LINCS [12]). CMap and LINCS contain gene expression profiles of dozens of cell lines treated with thousands of genetic and chemical perturbagens. CMap introduced connectivity scores to quantify similarities between compound responses and disease gene expression signatures. Identifying compounds with gene expression signatures either similar to those of known disease-treating drugs or that counter disease signatures can help in selecting therapeutic leads [13–16]. These strategies have successfully identified drugs with high in vitro efficacy [16] across a range of diseases [17–19].

Deep learning methods have been used for lead discovery by predicting gene expression responses to perturbagens, including perturbagens that were not yet experimentally tested [20–23]. However, these approaches rely on chemical and genetic libraries, meaning they select perturbagens from predefined libraries and cannot identify perturbagens as new combinations of drug targets. Further, they are perturbation response methods that predict changes in phenotypes upon perturbations. Thus, they can identify perturbagens by exhaustively predicting responses to all perturbations in the library and then searching for perturbagens with the desired response. Unlike existing methods that learn responses to perturbations, phenotype-based approaches need to solve the inverse problem, which is to infer perturbagens necessary to achieve a specific response – i.e., directly predicting perturbagens by learning which perturbations elicit a desired response.

In causal discovery, the problem of identifying which elements of a system should be perturbed to achieve a desired state is referred to as optimal intervention design [24–26]. Using insights from causal discovery and geometric deep learning, here we introduce PDGrapher, an approach for the combinatorial prediction of therapeutic targets that can shift gene expression from an initial diseased state to a desired treated state. PDGrapher is formulated using a causal model, where genes represent the nodes in a causal graph, and structural causal equations define their causal relationships. Given a genetic or chemical intervention dataset, PDGrapher pinpoints a set of genes that a perturbagen should target to facilitate the transition of node states from diseased to treated. PDGrapher utilizes protein-protein interaction networks (PPI) or gene regulatory networks (GRN) as approximations of the causal graph, operating under the assumption of no unobserved confounders. PDGrapher tackles the optimal intervention design using representation learning, using a graph neural network (GNN) to represent structural equations.

PDGrapher is trained on a dataset of disease-treated sample pairs to predict therapeutic gene targets that can shift the gene expression phenotype from a diseased to a healthy or treated state. Once trained, PDGrapher processes a new diseased sample and outputs a perturbagen—a set of therapeutic targets—predicted to counteract the disease effects in that specific sample. We evaluate PDGrapher across 19 datasets, comprising genetic and chemical interventions across 11 cancer types and two proxy causal graphs, and consider different evaluation setups, including settings where held out folds contain new samples in the same cell line with the training samples, and settings where held out folds contain new samples from a cancer type that PDGrapher has never encountered during training. In held out folds that contain new samples, PDGrapher detects up to 13.37% and 1.09% ground-truth therapeutic targets in chemical and genetic intervention datasets, respectively, than existing methods. We also find that in chemical intervention datasets, candidate therapeutic targets predicted by PDGrapher are on average up to 11.58% closer to ground-truth therapeutic targets in the gene-gene interaction network than what would be expected by chance. Even in held-out folds containing new samples from a previously unseen disease, PDGrapher maintains robust performance. Unlike methods that indirectly identify perturbagens by predicting cell responses, PDGrapher directly predicts perturbagens that can shift gene expression from diseased to treated states. This feature of PDGrapher enables model training up to 25 times faster than indirect prediction methods, such as scGen [22] and CellOT [27]. Since these approaches build a separate model for each perturbation, they become increasingly ineffective when applied to datasets with a large number of perturbagens. For example, with the default setting, cellOT needs 10 hours to train for a single perturbagen in a cell line from the LINCS dataset.

PDGrapher can aid in elucidating the mechanism of action of chemical perturbagens (Figure S3), which we show in the case of vorinonstat, a histone deacetylase inhibitor used to treat cutaneous T-cell lymphoma, and sorafeniv, a multi-kinase inhibitor used in the treatment of several types of cancers (Supplementary Note 1). PDGrapher can also suggest potential anti-cancer therapeutic targets: it highlighted *KDR* as a top-predicted target for non-small cell lung cancer (see heading *PDGrapher predicts therapeutic targets across cancer types* in the *Methods* section). It identified associated drugs—vandetanib, sorafenib, catequentinib, and rivoceranib—which inhibit the kinase activity of the protein encoded by *KDR*. These drugs block VEGF signaling, suppressing endothelial cell proliferation, migration, and blood vessel formation that tumors rely on for growth and metastasis [28, 29]. By predicting combinatorial therapeutic targets based on phenotypic transitions, PDGrapher provides a scalable approach to phenotype-driven perturbation modeling.

## Results

### Overview of intervention datasets and causal graphs

We evaluate our method across a total of 38 preprocessed datasets that span two types of interventions (genetic and chemical), eleven cancer types (lung, breast, prostate, colon, skin, cervical, head and neck, pancreatic, stomach, brain, and ovarian), and two types of proxy causal graphs: protein–protein interaction networks (PPI) and gene regulatory networks (GRN). Each dataset is uniquely defined by a combination of intervention type, causal graph type, cancer type, and cell line, and is denoted in the format: Treatment type-Graph type-Cancer Type-Cell Line. The chemical-PPI datasets include cell lines A549 (lung), MCF7, MDAMB231, BT20 (breast), PC3, VCAP (prostate), HT29 (colon), A375 (skin), and HELA (cervix). The genetic-PPI datasets include A549 (lung), MCF7 (breast), PC3 (prostate), HT29 (colon), A375 (skin), ES2 (ovary), BICR6 (head and neck), YAPC (pancreas), AGS (stomach), and U251MG (brain). Similarly, the chemical-GRN and genetic-GRN datasets span the same combinations of cancer types and cell lines as their PPI counterparts. This comprehensive collection of datasets enables systematic benchmarking of our model across diverse perturbation modalities, biological contexts, and graph structures. Genetic interventions are single-gene knockout experiments by CRISPR/Cas9-mediated gene knockouts, while chemical interventions are multiple-gene treatments induced using chemical compounds. We utilize a PPI network from BIOGRID that has 10,716 nodes and 151,839 undirected edges. We additionally construct gene regulatory networks for each disease-treatment type pair using GENIE3 [30] (Supplementary Note 3) with GRNs on average having 10,000 nodes and 500,000 directed edges. The training data for PDGrapher consists of two components: disease intervention data and treatment intervention data. Disease intervention data includes paired healthy and diseased gene expression profiles, along with associated disease genes; however, these data are only available for cell lines corresponding to lung, breast, and prostate cancers. In contrast, the treatment intervention data comprise paired diseased and treated gene expression samples along with genetic or chemical perturbagens, and are available across all datasets. Tables S11 and S12 summarize the number of samples for each cell line and intervention datasets.

### Overview of PDGrapher model

Given a diseased cell state (gene expression profile), the goal of PDGrapher is to predict the genes that, if targeted by a perturbagen, would shift the cell to a treated state (Figure 1A). Unlike methods for learning the response of cells to a given perturbation [22, 27, 31, 32], PDGrapher focuses on the inverse problem by learning which perturbation elicits a desired response. PDGrapher predicts perturbagens that shift cellular states under the assumption that an optimal perturbagen is one that alters the gene expression profile of a cell to closely match a desired target state. Our approach comprises two modules (Figure 1B). First, a perturbagen discovery module *f*_*p*_ takes the initial and desired cell states and outputs a candidate perturbagen as a set of therapeutic targets 𝒰′. Then, a response prediction module *f*_*r*_ takes the initial state and the predicted perturbagen 𝒰′ and predicts the cell response upon perturbing genes in 𝒰′. Our response prediction and perturbagen discovery modules are graph neural network (GNN) models that operate on a proxy causal graph, where edge mutilations represent the effects of interventions on the graph (Figure 1C).

**Figure 1.**
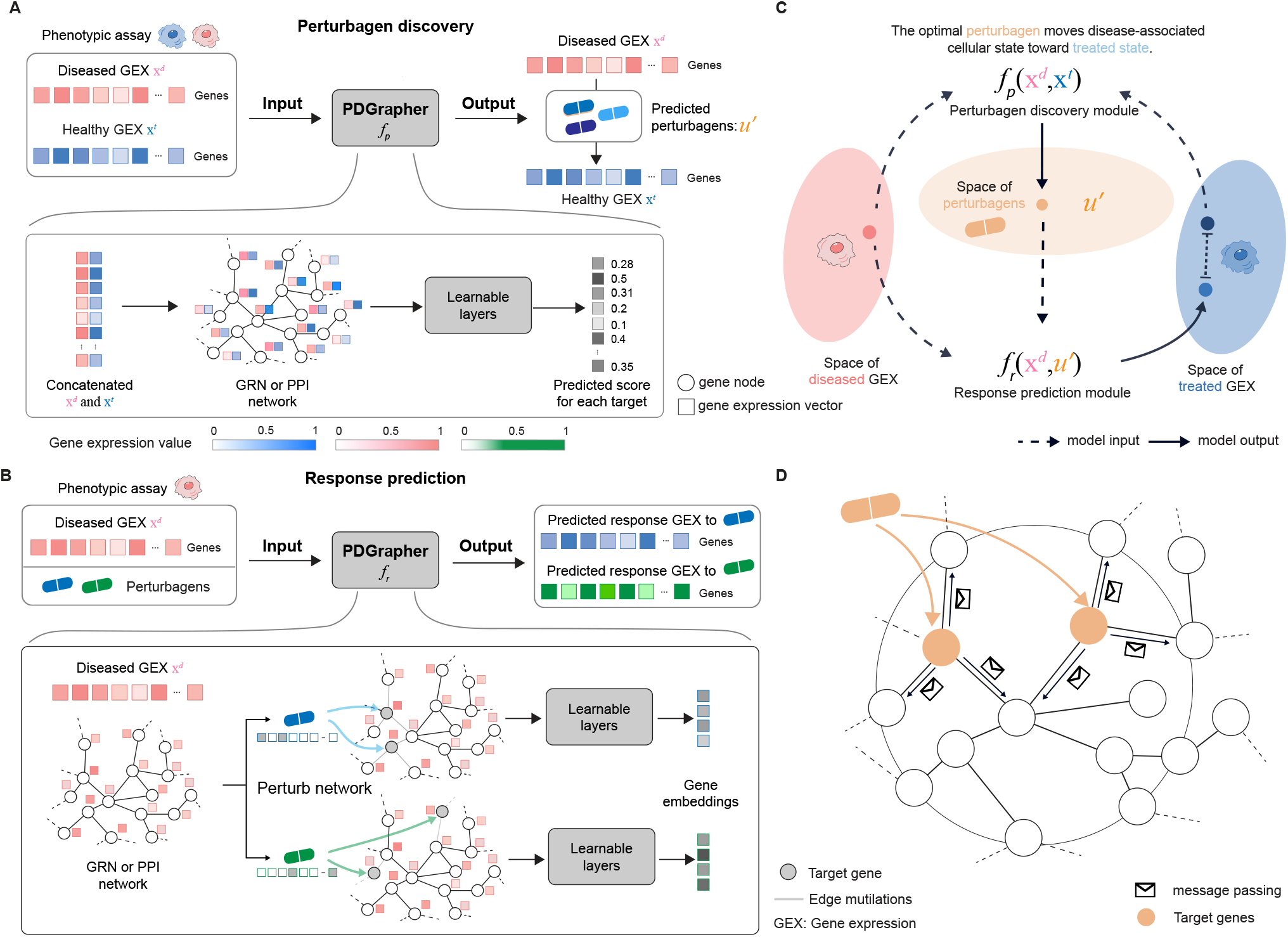
Overview of PDGrapher. **(A)** Given a paired diseased and treated gene expression samples and a proxy causal graph, PDGrapher’s perturbagen discovery module, *f*_*p*_, predicts a candidate set of therapeutic targets to shift cell gene expression from diseased to treated state. **(B)** Given a disease sample’s gene expression, a proxy causal graph, and a set of perturbagens, PDGrapher’s response prediction module, *f*_*r*_, predicts the gene expression response of the sample to each perturbagen. *f*_*r*_ represents perturbagens effects in the graph as edge mutilations. **(C)** *f*_*p*_ is optimized using 2 losses: a cross-entropy cycl(e loss to predicted a per)turbagen 𝒰′ which aims to shift the diseased cell state closely approximating the treated state: *CE* **x**^*t*^, *f*_*r*_(**x**^*d*^, *f*_*p*_(**x**^*d*^, **x**^*t*^))) (*with f*_*r*_ *frozen*), and a cross-entropy supervision loss that directly supervises the prediction of 𝒰′: *CE* (𝒰′, *f*_*p*_(**x**^*d*^, **x**^*t*^). See *Method section* for more details. **(D)** Both *f*_*r*_ and *f*_*p*_ follow the standard message-passing framework where node representations are updated by aggregating the information from neighbors in the graph.

PDGrapher is trained using an objective function with two components, one for each module, *f*_*r*_ and *f*_*p*_. The response prediction module *f*_*r*_ is trained using disease and treatment intervention data on cell state transitions, so that the predicted cell states are close to the known perturbed cell states upon interventions. The perturbagen discovery module *f*_*p*_ is trained only using the treatment intervention data; given a diseased cell state, *f*_*p*_ predicts the set of therapeutic targets 𝒰′ that caused the corresponding treated cell state. The objective function for the perturbagen discovery module consists of two elements: (1) a cycle loss that optimizes the parameters of *f*_*p*_ such that the response upon intervening on the predicted genes in 𝒰′, as measured by *f*_*r*_, closely approximates the actual treated cellular state; and (2) a supervision loss on the therapeutic targets set 𝒰′ that directly pushes PDGrapher to predict the correct perturbagen. Both models are trained simultaneously using early stopping independently so that each model finishes training upon convergence.

When trained, PDGrapher predicts perturbagens—as a set of candidate target genes—to shift cells from diseased to treated. Given a pair of diseased and treated samples, PDGrapher directly predicts perturbagens by learning which perturbations elicit target responses. In contrast, existing approaches are perturbation response methods that predict changes in phenotype that occur upon perturbation, thus, they can only indirectly predict perturbagens (Figure 2A). Given a disease-treated sample pair, a response prediction module (such as scGen [22], ChemCPA [33], Biolord [34], GEARS [35], or CellOT [27]) is used to predict the response of the diseased sample to a library of perturbagens. The predicted perturbagen is the one that produces a response that is the most similar to the treated sample. We evaluate PDGrapher’s performance in two separate settings (Figure 2BC): (1) a random splitting setting, where the samples are split randomly between training and test sets within a cell line (denoted as random for convenience), and (2) a leave-cell-out setting, where PDGrapher is trained in one cell line, and its performance is evaluated in a cell line the model never encountered during training to test how well the model generalizes to a new disease. Tables S1 and S2 show the numbers of unseen perturbagens in chemical perturbation datasets in the random and leave-cell-out splits, respectively; Tables S3 and S4 show the number of unseen perturbagens in genetic perturbation datasets in the random and leave-cell-out splits, respectively.

**Figure 2.**
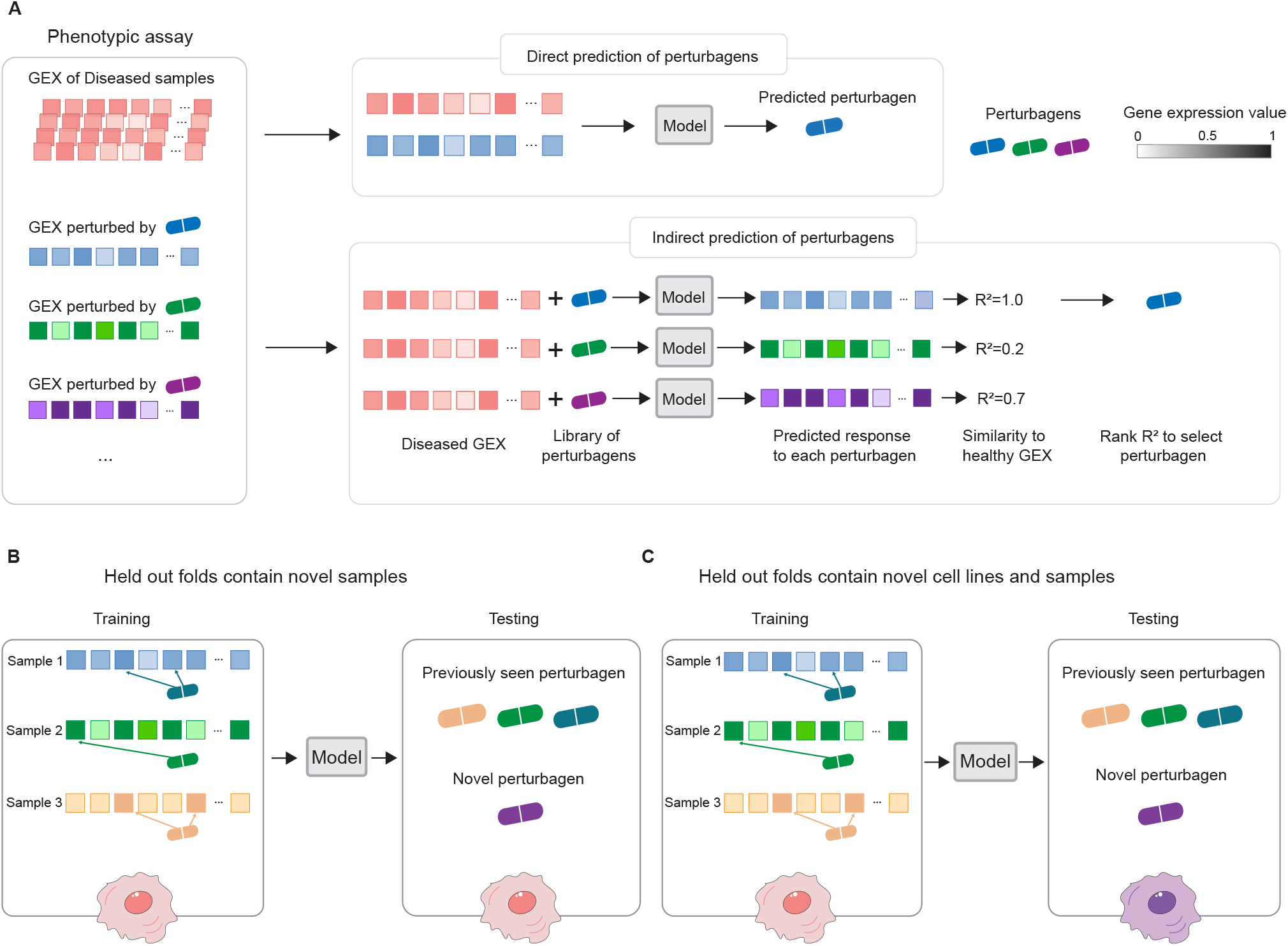
Overview of evaluation settings and data splits. **(A)** Given a dataset with paired diseased and treated samples and a set of perturbagens, PDGrapher makes a direct prediction of candidate perturbagens that shift gene expression from a diseased to a treated state, for each disease-treated sample pair. The direct prediction means that PDGrapher directly infers the perturbation necessary to achieve a specific response. In contrast to direct prediction of perturbagens, existing methods predict perturbagens only indirectly through a two-stage approach: For a given diseased sample, the model learns the response to each candidate perturbagen from an existing library and identifies the perturbagen whose induced response most closely approximates the desired treated state. Existing methods learn the response of cells to a given perturbation [22, 27, 31, 32], whereas PDGrapher focuses on the inverse problem by learning which perturbagen elicits a given response, even in the most challenging cases when the combinatorial composition of perturbagen was never seen before. **(B-C)** We evaluate PDGrapher’s performance across two settings: given a cell line, randomly splitting samples between training and testing set (B), and by splitting samples based on cell lines, where the model is trained on one cell line and evaluated on a different, previously unseen cell line to assess PDGrapher’s generalization performance. (C).

### PDGrapher predicts perturbagens to reverse disease states

In the random splitting setting, we assess the ability of PDGrapher for combinatorial prediction of therapeutic targets across chemical PPI datasets (Chemical-PPI-Lung-A549, Chemical-PPI-Breast-MCF7, Chemical-PPI-Breast-MDAMB231, Chemical-PPI-Breast-BT20, Chemical-PPI-Prostate-PC3, Chemical-PPI-Prostate-VCAP, Chemical-PPI-Colon-HT29, Chemical-PPI-Skin-A375, and Chemical-PPI-Cervix-HELA). Specifically, we measure whether, given paired diseased-treated gene expression samples, PDGrapher can predict the set of therapeutic genes targeted by the chemical compound in the diseased sample to generate the treated sample. Given paired diseased-treated gene expression samples, PDGrapher ranks genes in the dataset according to their likelihood of being therapeutic targets.

We quantify the ranking quality using normalized discounted cumulative gain (nDCG), where the gain reflects the ranking accuracy of the model. An nDCG value close to 1 indicates highly accurate predictions, with the top-ranked gene targets closely matching the ground-truth targets, while lower nDCG values indicate poorer ranking performance. This metric provides a normalized and scalable measure of ranking quality, enabling consistent comparison across different datasets and models. PDGrapher outperforms competing methods in all cell lines, achieving 0.02 (Chemical-PPI-Lung-A549), 0.13 (Chemical-PPI-Breast-MDAMB231), 0.03 (Chemical-PPI-Breast-BT20), 0.004 (Chemical-PPI-Breast-MCF7), 0.07 (Chemical-PPI-Prostate-VCAP), 0.005 (Chemical-PPI-Prostate-PC3), 0.03 (Chemical-PPI-Skin-A375), 0.06 (Chemical-PPI-Cervix-HELA), and 0.001 (Chemical-PPI-Colon-HT29), higher nDCG values compared to the second-best competing method (Figure 3B). In addition to evaluating the entire predicted target rank, it is even more practically crucial to assess the accuracy of the top-ranked predicted targets. Because perturbagens target multiple genes, we measure the fraction of samples in the test set for which we obtain a partially accurate prediction, where at least one of the top N predicted gene targets corresponds to an actual gene target (denoted as the percentage of accurately predicted samples). Here, N represents the number of known target genes of a perturbagen. PDGrapher consistently provides accurate predictions for more samples in the test set than competing methods. Specifically, it outperforms the second-best competing method by predicting ground-truth targets in an additional 7.73% (Chemical-PPI-Breast-MCF7), 9.32% (Chemical-PPI-Lung-A549), 13.37% (Chemical-PPI-Breast-MDAMB231), 4.50% (Chemical-PPI-Breast-BT20), 7.88% (Chemical-PPI-Prostate-PC3), 11.53% (Chemical-PPI-Prostate-VCAP), 7.56% (Chemical-PPI-Colon-HT29), 9.55% (Chemical-PPI-Skin-A375), and 8.41% (Chemical-PPI-Cervix-HELA) of samples (Figure 3A). We also evaluate the performance of PDGrapher using recall@1, recall@10, and recall@100, which calculate the ratio of true target genes included in the top 1, top 10, and top 100 predicted gene targets, respectively. Although the absolute recall values are modest due to the inherent difficulty of the task, PDGrapher consistently outperforms all competing methods, demonstrating its relative strength and robustness. Specifically, PDGrapher outperforms the second-best method in all the recall metrics with the averaged margin being 3.31% (Chemical-PPI-Lung-MCF7), 3.28% (Chemical-PPI-Lung-A549), 11.65% (Chemical-PPI-Breast-MDAMB231), 7.27% (Chemical-PPI-Breast-BT20), 2.50% (Chemical-PPI-Prostate-PC3), 9.53% (Chemical-PPI-Prostate-VCAP), 3.08% (Chemical-PPI-Colon-HT29), 2.87% (Chemical-PPI-Skin-A375), and 5.13% (Chemical-PPI-Cervix-HELA) (Figure 3C). We then consolidated the results using the rankings from experiments across different cell lines and metrics for each method. PDGrapher achieved the best overall rankings, with a median significantly higher than all competing methods. (Figure 3D). P-values of the chemical perturbagen discovery tests are provided in the source data of figures.

**Figure 3.**
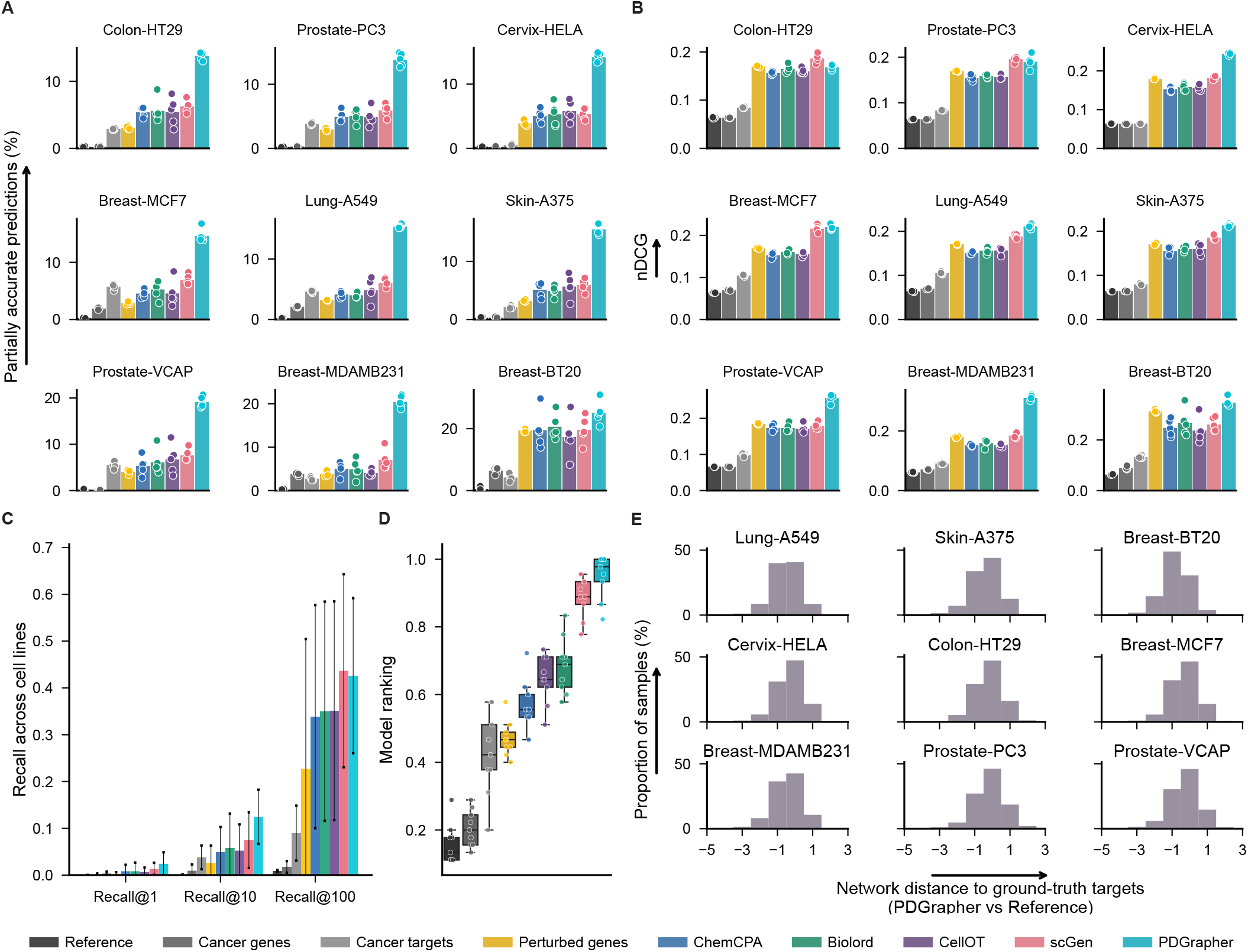
PDGrapher efficiently predicts chemical perturbagens to shift cells from diseased to treated states in held out folds containing novel samples. PDGrapher demonstrates improved performance across nine chemical perturbation datasets with various diseases, yielding up to 13.37% more accurately predicted samples in the testing sets compared to the second-best baseline (e.g., Chemical-PPI-Breast-MDAMB231: 20.43% vs 7.05%, **A**) and up to 0.13 (Chemical-PPI-Breast-MDAMB231: 0.31 vs 0.18, **B**) higher nDCG than the second-best baseline. In (A) and (B), bars show the average performance across 5 cross-validation test splits for each of the nine chemical datasets. Overlaid points represent performance values from individual data splits (n=5 per cell line). Each data split contains 20% of samples in the dataset, with each sample corresponding to a perturbation-response instance, and where replicates exist for a given drug, they are treated as independent inputs during training and evaluation. **(C)** PDGrapher recovers ground-truth therapeutic targets at higher rates (evaluated by recall 1 - 100) compared to competing methods for Chemical-PPI datasets. **(D)** Boxplots show the distribution of average model rankings across 9 cell lines (n=9); each dot corresponds to the aggregated ranking value across cross-validation splits and across all metrics for a distinct cell line. Higher value indicates better performance. The central line inside the box represents the median, while the top and bottom edges correspond to the first (Q1) and third (Q3) quartiles. The whiskers extend to the smallest and largest values within 1.5 times the interquartile range (IQR) from the quartiles. Each dot represents a data point for a specific cell line. P-values from the statistical tests are provided in the Source Data. **(E)** Shown is the difference of shortest-path distances between ground-truth therapeutic genes and predicted genes by PDGrapher and a Random baseline across nine cell lines. Predominantly negative values indicate that PDGrapher predicts sets of therapeutic genes that are closer in the network to ground-truth therapeutic genes compared to what would be expected by chance [average shortest-path distances across cell lines for PDGrapher vs Random = 2.77 vs 3.11].

PDGrapher not only provides accurate predictions for a larger proportion of samples and consistently predicts ground-truth therapeutic targets close to the top of the ranked list, but it also predicts gene targets that are closer in the network (measured by the shortest path distance) to ground-truth targets compared to what would be expected by chance (Figure 3E). In all cell lines, the ground-truth therapeutic targets predicted by PDGrapher are significantly closer to the ground-truth targets compared to what would be expected by chance (Table S6). For example, for Chemical-PPI-Lung-A549, the median distance between the predicted and ground-truth therapeutic targets is 3.0 for both PDGrapher and the Random reference. However, the distributions exhibit a statistically significant difference, with a one-sided Mann-Whitney U test that yields a p-value *<* 0.001, an effect size (rank-biserial correlation) of 0.3531 (95% CI [0.3515, 0.3549]), and a U statistic of 1.29e11. Similarly, for Chemical-PPI-Breast-MCF7, the median distance is 3.0 for both groups, yet the distributions are significantly different (p-value *<* 0.001, effect size = 0.2160 (95% CI [0.2145, 0.2173]), U statistic = 3.91e11) (Table S6). This finding suggests that PDGrapher predicts targets in a manner that reflects protein-protein interaction structure [32]. According to the local network hypothesis, which posits that genes in closer network proximity tend to be more functionally similar, PDGrapher’s predictions are more functionally related to ground-truth targets than would be expected by chance [36–38].

PDGrapher also exhibits superior performance across genetic datasets, specifically Genetic-PPI-Lung-A549, Genetic-PPI-Breast-MCF7, Genetic-PPI-Prostate-PC3, Genetic-PPI-Skin-A375, Genetic-PPI-Colon-HT29, Genetic-PPI-Ovary-ES2, Genetic-PPI-Head-BICR6, Genetic-PPI-Pancreas-YAPC, Genetic-PPI-Stomach-AGS, and Genetic-PPI-Brain-U251MG (Extended Data Figure 2). Briefly, PDGrapher successfully detected ground-truth targets in 0.87% (Genetic-PPI-Lung-A549), 0.50% (Genetic-PPI-Breast-MCF7), 0.24% (Genetic-PPI-Prostate-PC3), 0.38% (Genetic-PPI-Skin-A375), 0.36% (Genetic-PPI-Colon-HT29), 1.09% (Genetic-PPI-Ovary-ES2), 0.54% (Genetic-PPI-Head-BICR6), 0.11% (Genetic-PPI-Pancreas-YAPC), and 0.92% (Genetic-PPI-Brain-U251MG) more samples compared to the second-best competing method (Extended Data Figure 2A). Its ability to effectively predict targets at the top of the ranks is further supported by the metrics recall@1 and recall@10 (Extended Data Figure 2C). P-values of the genetic perturbagen discovery tests are provided in the source data of figures. PDGrapher and competing methods perform worse on genetic data than on chemical data. This may be due to knockout interventions generating weaker phenotypic signals than small molecule interventions. While gene knockouts are essential for understanding gene function, single-gene knockout studies can offer limited insights into complex cellular processes due to compensatory mechanisms [39–41]. Despite the modest performance in genetic intervention datasets, PDGrapher outperforms competing methods in the combinatorial prediction of therapeutic targets.

PDGrapher achieves the best performance in response prediction for both chemical (Extended Data Figure 1) and genetic perturbation (Extended Data Figure 2E-G). P-values of the response prediction tests are provided in the source data of the figures. When using GRNs as proxy causal graphs, PDGrapher has comparable performance with GRN and PPI across both genetic and chemical intervention datasets (Figures S4 and S5). One difference is that gene regulatory networks were constructed individually for each cell line, which makes leave-cell-out splitting setting prediction particularly challenging. Therefore, we only conducted random splitting setting experiments for GRN datasets. We also used PDGrapher to clarify the mode of action of the chemical perturbagens vorinostat and sorafenib in Chemical-PPI-Lung-A549 (Supplementary Note 1).

### PDGrapher generalizes to cell lines unseen during training

PDGrapher shows consistently strong performance on chemical intervention datasets in the leave-cell-out setting (Figure 4). In this setting, we use the trained models in the random-splitting setting for each cell line to predict therapeutic targets in the remaining cell lines. PDGrapher successfully predicts perturbagens that describe the cellular dynamics and shift gene expression phenotypes from a diseased to a treated state in 7.16% (Chemical-PPI-Breast-MCF7), 6.50% (Chemical-PPI-Lung-A549), 5.00% (Chemical-PPI-Breast-MDAMB231), 8.67% (Chemical-PPI-Prostate-PC3), 7.72% (Chemical-PPI-Prostate-VCAP), 7.31% (Chemical-PPI-Skin-A375), 7.08% (Chemical-PPI-Colon-HT29), and 7.13% (Chemical-PPI-Cervix-HELA) additional testing samples compared to the second-best competing method (Figure 4A). PDGrapher also outperforms the competing methods in eight of out nine cell lines by predicting nDCG values 0.02 (Chemical-PPI-Breast-MCF7), 0.01 (Chemical-PPI-Lung-A549), 0.01 (Chemical-PPI-Breast-MDAMB231), 0.03 (Chemical-PPI-Prostate-PC3), 0.02 (Chemical-PPI-Prostate-VCAP), 0.01 (Chemical-PPI-Skin-A375), 0.03 (Chemical-PPI-Colon-HT29), and 0.02 (Chemical-PPI-Cervix-HELA) higher than the second-best competing method (Figure 4B). Considering the overall performance across different cell lines and metrics, PDGrapher achieves the highest rank, with a median surpassing all competing methods (Figure 4D). Combinations of therapeutic targets predicted by PDGrapher in chemical datasets are closer to ground-truth targets than expected by chance (Figure 4E, Table S7). For example, for Chemical-PPI-Lung-A549, the median distance between predicted and ground-truth therapeutic targets is 3.0 for both PDGrapher and the Random reference. However, the distributions exhibit a statistically significant difference, with a one-sided Mann-Whitney U test yielding a p-value *<* 0.001, an effect size (rank-biserial correlation) of 0.2191 (95% CI [0.2182, 0.2200]), and a U statistic of 2.46e12. Similarly, for Chemical-PPI-Breast-MCF7, the median distance is 3.0 for both groups, yet the distributions are significantly different (p-value *<* 0.001, effect size = 0.2457 (95% CI [0.2450, 0.2464]), U statistic = 6.07e12) (Table S7). PDGrapher also outperforms existing methods in genetic perturbagen prediction across cell lines, as measured by the top targets on the predicted gene ranks (Figure S2A-D). PDGrapher also demonstrates superior performance in response prediction for both chemical (Figure S1) and genetic datasets (Figure S2E-G). P-values of the leave-cell-out perturbagen discovery tests and response prediction tests are provided in the source data of figures, respectively

**Figure 4.**
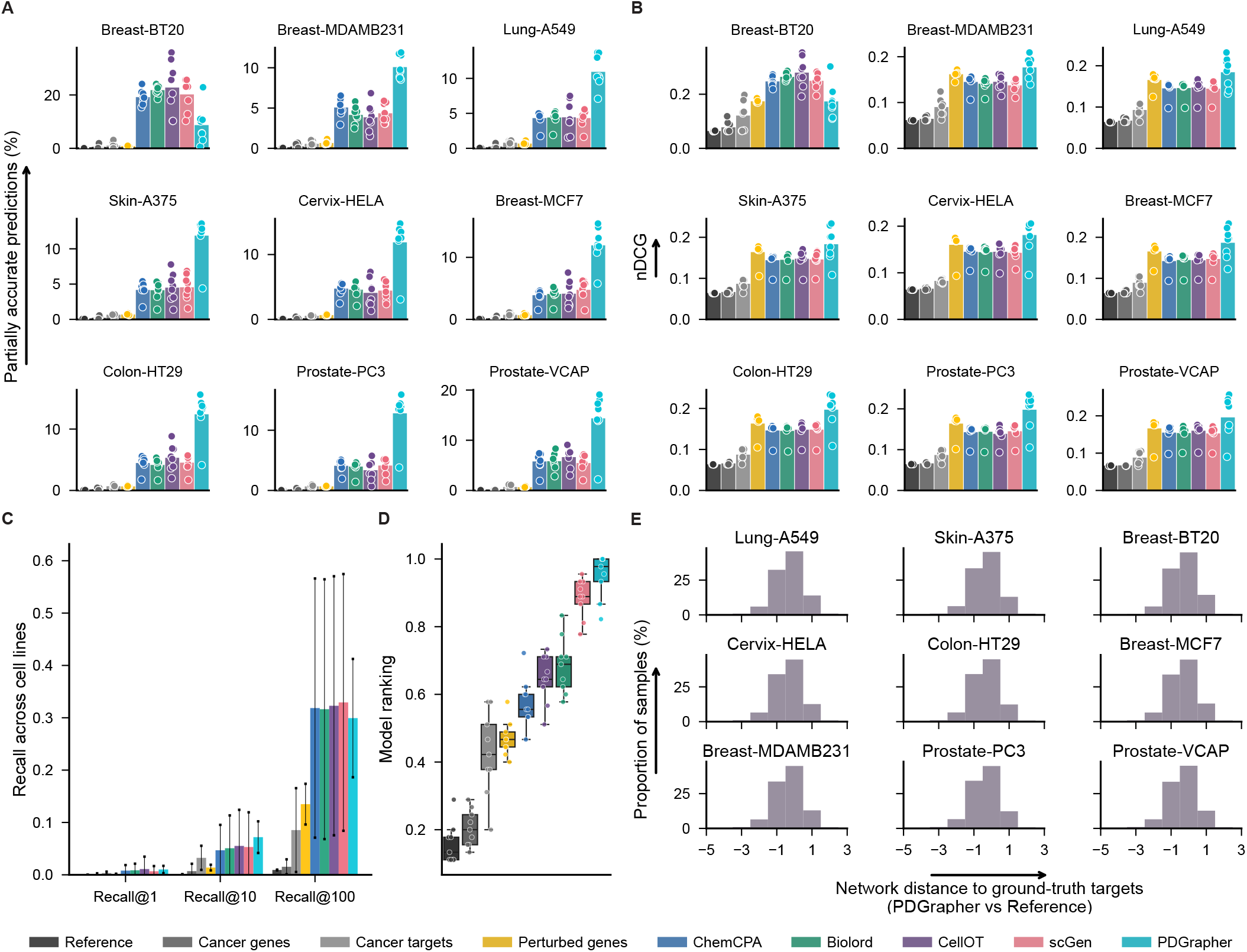
PDGrapher generalizes to new (previously unseen) cell lines and learns optimal chemical perturbagens in held out folds that contain both novel cell lines and novel samples. PDGrapher demonstrates improved performance when trained on nine chemical perturbation datasets spanning various diseases and evaluated on the remaining eight cell lines. It achieves up to 8.67% more accurately predicted samples in the testing sets compared to the second-best baseline (e.g., trained on Chemical-PPI-Prostate-PC3: 12.81% vs 4.13%, **A**) and up to 0.03 higher nDCG than the second-best baseline (e.g., trained on Chemical-PPI-Colon-HT29: 0.19 vs 0.16, **B**). In (A) and (B), bars show the average performance across 5 cross-validation test splits for each of the nine chemical datasets. Overlaid points represent performance values from individual data splits (n=5 per cell line). Each data split contains 20% samples in the dataset, with each sample corresponding to a perturbation-response instance, and where replicates exist for a given drug, they are treated as independent inputs during training and evaluation. **(C)** PDGrapher recovers ground-truth therapeutic targets at higher rates (evaluated by recall 1 - 100) compared to competing methods for Chemical-PPI datasets. **(D)** Boxplots show the distribution of average model rankings across 9 cell lines (n=9); each dot corresponds to the aggregated ranking value across cross-validation splits, train cell lines, and across all metrics for a distinct cell line. Higher value indicates better performance. The central line inside the box represents the median, while the top and bottom edges correspond to the first (Q1) and third (Q3) quartiles. The whiskers extend to the smallest and largest values within 1.5 times the interquartile range (IQR) from the quartiles. Each dot represents a data point for a specific cell line and metrics. P-values from the statistical tests are provided in the Source Data. **(E)** Shown is the difference of shortest-path distances between ground-truth therapeutic genes and predicted genes by PDGrapher and a Random baseline across nine cell lines. Predominantly negative values indicate that PDGrapher predicts sets of therapeutic genes that are closer in the network to ground-truth therapeutic genes compared to what would be expected by chance [average shortest-path distances across cell lines for PDGrapher vs Random = 2.75 vs 3.11].

Approaches that train individual models for each perturbagen (such as scGEN and CellOT) generally achieve better perturbagen prediction performance than those that use a single model for all perturbagens (Biolord, GEARS, and ChemCPA). However, training individual models becomes infeasible for large-scale datasets with many perturbagens. For example, without parallelization, scGEN would require about eight years to complete the leave-cell-out experiments on the chemical and genetic perturbation data used in this study. PDGrapher addresses this scalability challenge. Its training is up to 25 times faster than scGEN and more than 100 times faster than CellOT when using the default setting of 100,000 epochs, substantially reducing computational costs. This efficiency highlights a key advantage of PDGrapher. This improved efficiency is due to PDGrapher’s approach. Existing methods predict phenotypic responses to perturbations and identify perturbagens indirectly by searching through predicted responses for all candidates. In contrast, PDGrapher directly infers the perturbagen needed to achieve a specific response, learning which perturbations elicit a desired effect.

## PDGrapher identifies therapeutic targets validated through clinical and biological evidence

We examined PDGrapher’s ability to predict targets of anti-cancer drugs that were not encountered by the model during training time using chemical cell lines with matched healthy data: Chemical-PPI-Lung-A549, Chemical-PPI-Breast-MCF7, Chemical-PPI-Breast-MDAMB231, Chemical-PPI-Breast-BT20, Chemical-PPI-Prostate-PC3, Chemical-PPI-Prostate-VCAP. PDGrapher was used to predict gene targets to shift these diseased cell lines into their healthy states. Figure 5A shows the recovery of targets of FDA-approved drugs for varying values of K (K represents the number of predicted target genes considered in the predicted ranked list), indicating that PDGrapher can identify targets of approved anti-cancer drugs not seen during training among the top predictions.

**Figure 5.**
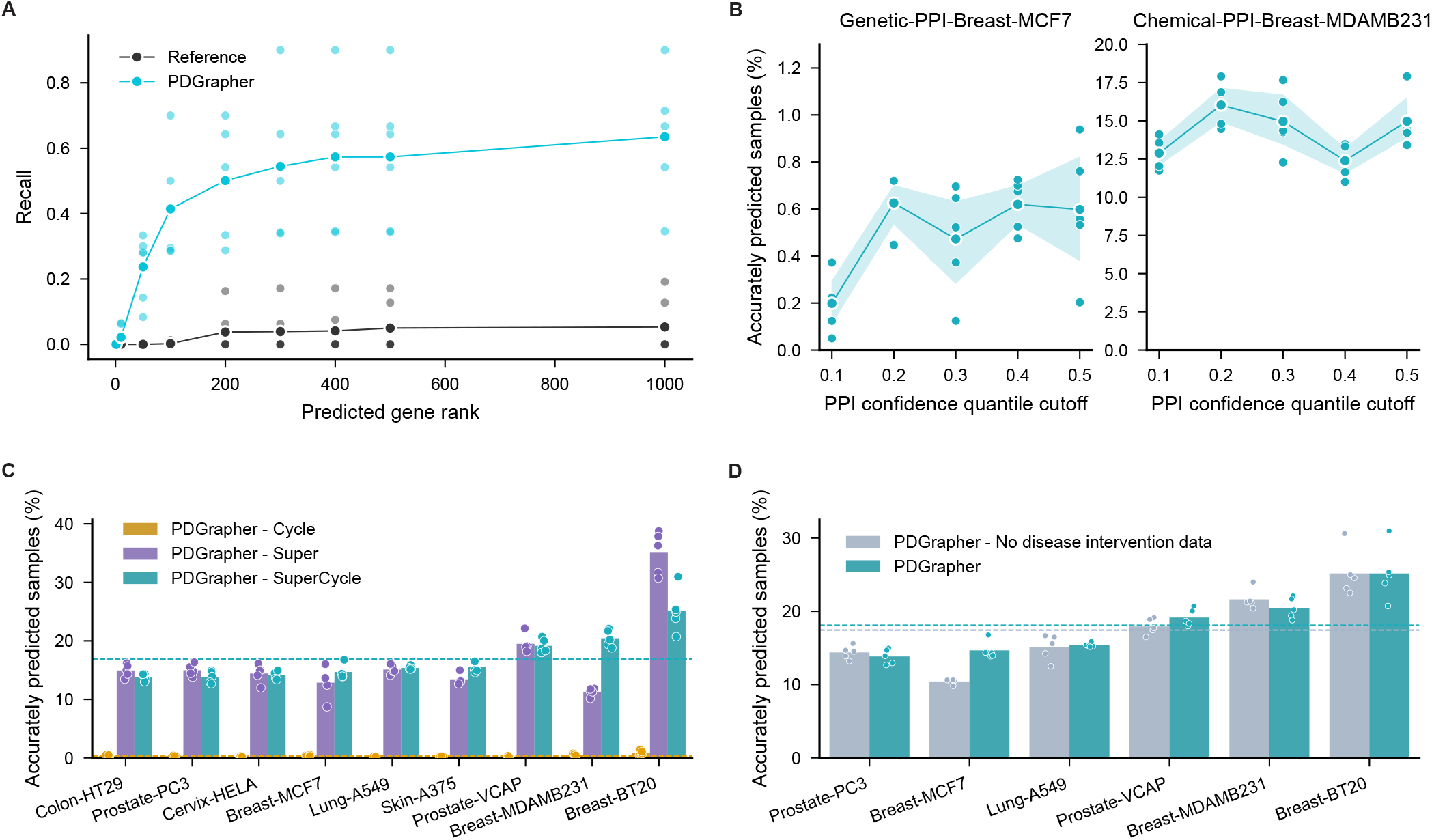
PDGrapher’s exhibits robust performance across training strategies, PPI networks, and data availability settings. **(A)** Performance of PDGrapher in the prediction of unseen approved drug targets to reverse disease effects across all cell lines with healthy counterparts in chemical perturbation datasets. Individual datapoints represent individual cell lines (n=6). **(B)** Performance of sensitivity analyses evaluated by percentage of accurately predicted samples for cell lines MDAMB231 and MCF7 under chemical and genetic perturbations, respectively. The PPI used here is from STRING (string-db.org) with a confidence score for each edge. The edges are filtered by the 0.1, 0.2, 0.3, 0.4, and 0.5 quantiles of the confidence scores as cutoffs, resulting in five PPI networks with 625,818, 582,305, 516,683, 443,051, and 296,451 edges, respectively. Data are presented as mean values across 5 cross-validation data splits per PPI confidence quantile. Shaded bands represent *±*1 standard deviation (SD) from the mean (n=5 computational replicates per quantile). Each point corresponds to performance on a specific data split. **(C)** Performance metrics of ablation study on PDGrapher’s objective function components: PDGrapher-Cycle trained using only the cycle loss, PDGrapher-SuperCycle trained using the supervision and cycle loss, and PDGrapher-Super trained using only the supervision loss, evaluated by percentage of accurately predicted samples. PDGrapher-Cycle demonstrates inferior performance, resulting in limited visibility in the bar plot. **(D)** Performance metrics of the second ablation study on PDGrapher’s input data: PDGrapher *- No disease intervention* data using only treatment intervention data and PDGrapher using both disease and treatment intervention data. The disease and treatment intervention data are organized as (healthy, mutation, disease) and (diseased, drug, treated), respectively. In (C) and (D), bars show the average performance across 5 cross-validation test splits for each of the nine chemical datasets. Overlaid points represent performance values from individual data splits (n=5 per cell line). Each data split contains 20% samples in the dataset, with each sample corresponding to a perturbation-response instance, and where replicates exist for a given drug, they are treated as independent inputs during training and evaluation. P-values from the statistical tests are provided in the Source Data.

We analyzed lung cancer by comparing the targets predicted by PDGrapher for lung cancer cell lines with the targets of candidate drugs in clinical development, curated from the Open Targets platform [42]. This evaluation tested PDGrapher’s ability to predict combinatorial chemical perturbagens. We compared the top 10 targets predicted by PDGrapher for the A549 lung cancer cell line to ten randomly selected genes. The predicted targets had significantly higher Open Targets scores and more supporting resources than the random genes (Extended Data Figure 7). Using an Open Targets evidence cutoff score of 0.5, 8 out of 10 predicted targets had evidence supporting their association with lung cancer, compared to only 2 out of 10 in the random gene set. Four drugs, tacedinaline (DrugBank ID: DB12291 and clinical trial ID: NCT00005093), selpercatinib (DrugBank ID: DB15685), pralsetinib (DrugBank ID: DB15822), and dexmedetomidine (Drug-Bank ID: DB00633 and [43]), targeting these predicted genes were not included in the training set but have been identified as potential treatments for non-small cell lung cancer(NSCLC).

We then evaluated PDGrapher’s predictions by examining FDA-approved drugs that were not present in the training set of PDGrapher. Specifically, we assessed PDGrapher’s performance using the Chemical-PPI-Lung-A549 dataset, focusing initially on pralsetinib, a targeted cancer therapy primarily used to treat NSCLC [44]. Pralsetinib is a selective RET kinase inhibitor designed to block the activity of RET proteins that have become aberrantly active due to mutations or fusions. Pralsetinib is known to target 11 key proteins: RET, DDR1, NTRK3, FLT3, JAK1, JAK2, NTRK1, KDR, PDGFRB, FGFR1, and FGFR2 [45]. Ret proto-oncogene (*RET*), the gene encoding for pralsetinib’s primary target, was ranked 11th out of 10,716 genes in the predicted list. Half of the genes encoding pralsetinib’s targets (5 out of 11) were ranked within the top 100 predicted targets by PDGrapher, including kinase insert domain receptor (*KDR*; ranked at 3), fms related receptor tyrosine kinase 3 (*FLT3*; ranked at 10), *RET* (ranked at 11), platelet derived growth factor receptor beta (*PDGFRB*; ranked at 14), and fibroblast growth factor receptor 2 (*FGFR2*; ranked at 81). This substantial overlap highlights the potential of the candidate targets identified by PDGrapher for pralsetinib-based lung cancer treatment, given that pralsetinib was not included in the training set of PDGrapher.

Next, we examined *KDR* as a therapeutic target for lung cancer. *KDR*, also known as *VEGFR-2*, has been identified as a significant therapeutic target in A549 lung adenocarcinoma cells. These cells express *KDR* at both mRNA and protein levels, facilitating autocrine signaling that promotes tumor cell survival and proliferation [28]. Activation of *KDR* enhances tumor angiogenesis and growth by upregulating oncogenic factors such as Enhancer of Zeste Homolog 2 (*EZH2*), which is associated with increased cell proliferation and migration. Inhibiting KDR has demonstrated promising therapeutic effects, including reduced cell proliferation and induced apoptosis. For instance, KDR inhibitors have been shown to decrease the malignant potential of lung adenocarcinoma cells by downregulating *EZH2* expression and increasing sensitivity to chemotherapy [29]. These findings underscore the importance of KDR as a therapeutic target in A549 lung adenocarcinoma cells, highlighting its role in tumor progression and the potential benefits of its inhibition in cancer treatment strategies. Importantly, PDGrapher has successfully identified *KDR* among the top 20 predicted targets in chemical-PPI-lung-A549, validating its precision in detecting key therapeutic targets for lung cancer.

Given that Open Targets offers more comprehensive evidence for targets currently under development, we conducted a second series of case studies using Open Targets data to evaluate PDGrapher’s capability to identify candidate therapeutic targets and drugs. This analysis aims to identify targets for lung cancer. Figure 6A presents a bubble graph that illustrates the union of the top 10 predicted targets to transition cell states from diseased to healthy in the six cell lines of three types of cancer that have available healthy controls. In the plot, the color intensity and size of the bubbles represent the number of evidence sources and the association scores for each type of evidence, respectively. Most predicted targets are supported by drugs, pathology and systemic biology, and somatic mutation databases, which were considered strong evidence sources. Two unique targets, DNA topoisomerase II alpha (*TOP2A*) and cyclin dependent kinase 2 (*CDK2*), are predicted exclusively for the lung cancer cell line (Extended Data Figure 8AB). *TOP2A* is ranked as the top predicted target by PDGrapher. This gene encodes a crucial decatenating enzyme that alters DNA topology by binding to two double-stranded DNA molecules, introducing a double-strand break, passing the intact strand through the break, and disciplining the broken strand. This mechanism is vital for DNA replication and repair processes. *TOP2A* could be a potential therapeutic target for anti-metastatic therapy of NSCLC, since it promotes metastasis of NSCLC by stimulating the canonical Wnt signaling pathway and inducing epithelial–mesenchymal transition [46]. Using the predicted target of *TOP2A*, PDGrapher then identified three drugs, aldoxorubicin, vosaroxin, and doxorubicin hydrochloride, as candidate drugs. These drugs were not part of the training dataset of PDGrapher and are in the early stages of clinical development: aldoxorubicin and vosaroxin are in Phase II trials (ClinicalTrials.gov); doxorubicin hydrochloride is in Phase I but has been shown to improve survival in patients with metastatic or surgically unresectable uterine or soft tissue leiomyosarcoma [47].

**Figure 6.**
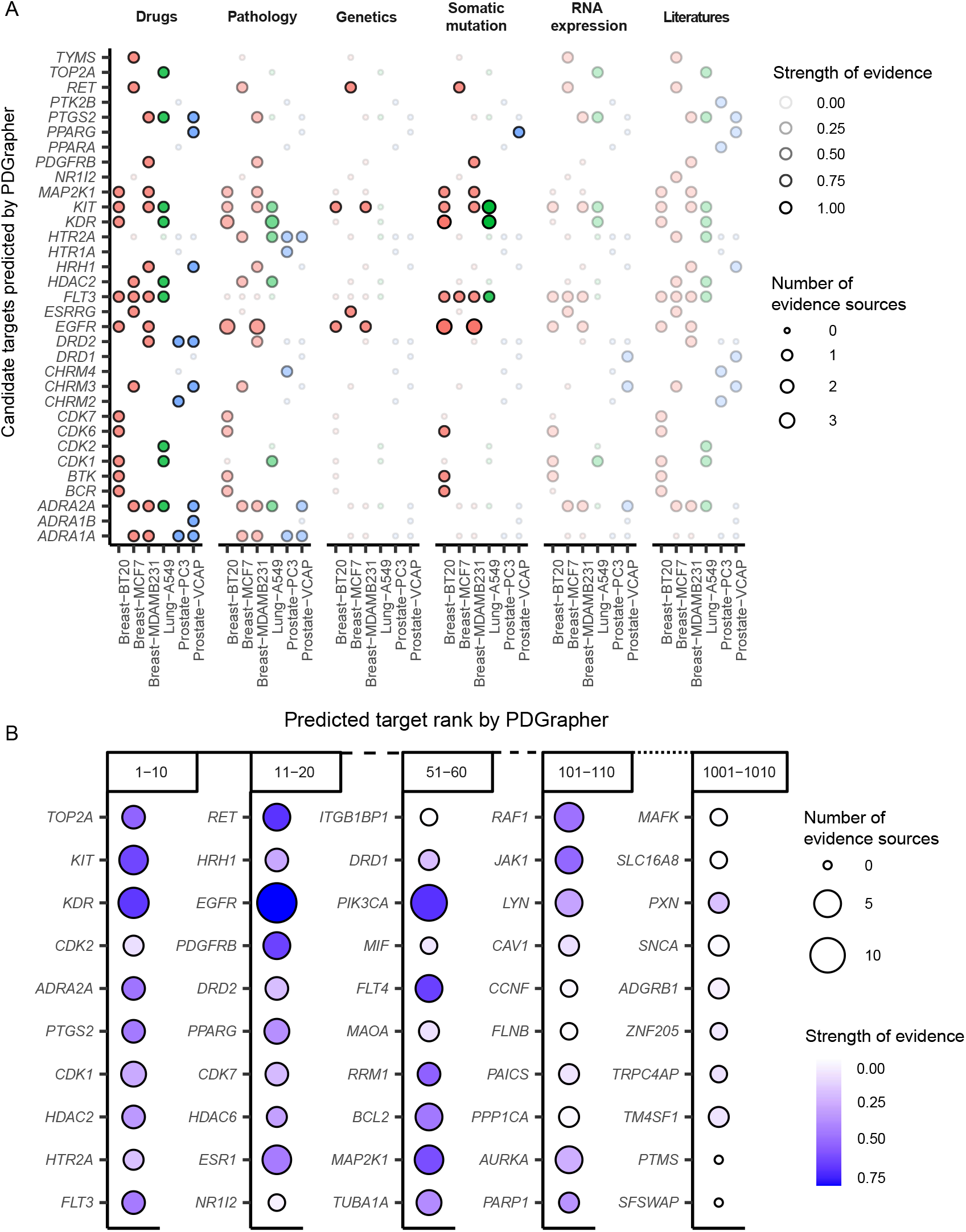
PDGrapher’s prioritization of lung cancer targets is supported by Open Targets evidence. (**A**) Union of the top 10 targets predicted by PDGrapher in lung, breast, and prostate cancer. The color intensity and size of the bubbles represent the number of evidence sources and the association scores for each type of evidence, respectively. Red, blue, and green dots stand for breast, lung, and prostate cancer, respectively. See details of the scoring system in Supplementary Note 2. (**B**) Predicted target rank from PDGrapher in five ranges, 1-10, 11-20, 51-60, 101-110, 501-510, and 1001-1010, for lung cancer (Chemical-PPI-Lung-A549). The color intensity and size of the bubbles represent the number of evidence sources and the global scores of targets from OpenTarget, respectively.

Given that PDGrapher can rank all genes based on PPI or GRN data, we assessed two questions: whether top-ranked genes have stronger evidence from Open Targets compared to lower-ranked genes, and what rank threshold should be used to identify reliably predicted genes. Figure 6B shows the number of sources of evidence and the global scores for the predicted target genes within the rank ranges of 1-10, 11-20, 51-60, 101-10, 501-510 and 1001-1010 for lung cancer (Chemical-PPI-Lung-A549). The analysis revealed a clear trend: both the number of supporting evidence sources and global scores decrease with increasing rank, validating the predictive accuracy of PDGrapher. Most targets ranked within the top 100 have strong evidence from Open Targets, indicating that a rank threshold of 100 could serve as a cut-off for selecting candidate targets.

### Training PDGrapher models

We conduct an ablation study to evaluate the components of PDGrapher’s objective function using chemical datasets. We train PDGrapher under three configurations: with only the cycle loss (PDGrapher-Cycle), only the supervision loss (PDGrapher-Super), and both losses combined (PDGrapher-SuperCycle). The experiments are performed in the random splitting setting across all nine PPI chemical datasets. We assess performance using several metrics, including the percentage of accurately predicted samples (Figure 5C), nDCG (Figure S8A), recall values (Figure S8B), and strength of evidence (Extended Data Figure 9). Results show that PDGrapher-Super achieves the highest performance in predicting correct perturbagens but performs the worst in reconstructing treated samples. In contrast, PDGrapher-Cycle performs poorly in identifying correct perturbagens but shows improved performance in predicting (reconstructing) held-out treated samples. PDGrapher-SuperCycle (the configuration used throughout this study) strikes a balance between these two objectives, achieving competitive performance in predicting therapeutic genes while demonstrating the best performance in reconstructing treated samples from diseased samples after intervening on the predicted genes. This makes PDGrapher-SuperCycle the most effective choice for balancing accuracy in perturbagen prediction with reconstruction fidelity. The findings show that supervision loss is essential for PDGrapher’s overall performance. The PDGrapher-Cycle model consistently underperforms in all cell lines and metrics. Although PDGrapher-Super often excels in ranking performance, including cycle loss (in PDGrapher-SuperCycle) proves its value by moderately improving top prediction metrics such as recall@1 and recall@10. Besides, when healthy cell line data is available, the top predictions of PDGrapher-SuperCycle demonstrates stronger strength of evidence compared to those of PDGrapher-Super in more than half (4/6) of the cell lines (Extended Data Figure 9). We chose PDGrapher-SuperCycle for this work because it provides accurate target gene predictions from the top-ranked genes in the predicted list and bases its predictions on the changes they would induce in diseased samples.

Recognizing the role of biological pathways in disease phenotypes, PDGrapher-SuperCycle can identify alternative gene targets with close network proximity that may produce similar phenotypic outcomes. The organization of genes with similar functions, where each gene contributes to specific biochemical processes or signaling cascades, allows perturbations in different genes to yield analogous effects [48]. This function-based interconnectivity implies that targeting different genes with similar functions can achieve therapeutic outcomes, as these genes collectively influence cellular phenotypic states [49]. Although PDGrapher-SuperCycle shows slightly lower performance than PDGrapher-Super in pinpointing targets (Figure 5C), it excels in identifying sets of gene targets capable of transitioning cell states from diseased to treated conditions (Figure S8B and Extended Data Figure 9).

We conducted four analyses to test the sensitivity of PDGrapher to the causal graph. The first analysis uses five PPI networks constructed with varying edge confidence cutoffs. The PPI network was obtained from STRING (https://string-db.org/) [50], which assigns a confidence score to each edge. To create networks with different levels of confidence, we filter edges based on the quantiles 0.1, 0.2, 0.3, 0.4, and 0.5 of the confidence scores, resulting in five networks with decreasing numbers of edges. For this analysis, we selected two cell lines: Chemical-PPI-Breast-MDAMB231 and Genetic-PPI-Breast-MCF7. The LINCS perturbation data for each cell line were processed using the five PPI networks (Supplementary Note 3). We trained PDGrapher with 1, 2, and 3 GNN layers, selecting the best configuration based on the performance of the validation set. As shown in Figure 5B and Extended Data Figure 3, PDGrapher performs robustly at all levels of confidence in PPIs. It maintains stable performance on both chemical and genetic intervention datasets, even as an increasing number of edges is removed from the PPI networks. The second to fourth analyses are based on the synthetic graphs. We created two sparse PPI networks using different edge removal strategies and one synthetic gene expression dataset with increasing levels of latent confounders. We applied two edge removal strategies to the PPI network: removing increasing numbers of either bridge edges or random edges. Details of the data generation process are provided in the *Methods*. Results from the edge removal experiments indicate that while bridge edges are structurally critical, their limited number in the PPI graph reduces their overall impact on model predictions (Extended Data Figure 5). In contrast, the removal of random edges, which include both high-confidence and redundant connections, has a more pronounced effect on performance, highlighting the model’s sensitivity to network perturbations (Extended Data Figure 6). The fourth dataset introduces latent confounders in the gene expression data. PDGrapher demonstrated stable performance in perturbagen prediction, with only a slight decrease in performance as stronger confounders were introduced (Extended Data Figure 4).

We then evaluate whether PDGrapher can maintain robust performance in the absence of disease intervention data. In our training datasets, some cell lines lack associated healthy control samples from disease-relevant tissues and cell types. These cell lines contain only treatment intervention data (diseased cell state, perturbagen, treated cell state) without disease intervention data (healthy cell state, disease mutations, diseased cell state) for model training and inference. For cell lines with healthy controls, we trained the response prediction module using both intervention datasets. For cell lines without healthy controls, we trained PDGrapher using only treatment intervention data. To evaluate PDGrapher’s dependency on healthy control data, we trained the model on cell lines with available disease intervention data under two conditions: one using the disease intervention data for training and one excluding it. This evaluation was conducted on six chemical perturbation datasets (Chemical-PPI-Lung-A549, -Breast-MCF7, -Breast-MDAMB231, -Breast-BT20, -Prostate-PC3, -Prostate-VCAP) and three genetic perturbation datasets (Genetic-PPI-Lung-A549, -Breast-MCF7, -Prostate-PC3).. The results indicate that the two versions of PDGrapher perform consistently across cell types and data types (chemical and genetic; Figure 5D, Figure S6, and S7). In half of the cell lines (4/9), the model trained without disease intervention data outperformed the model trained with it. This demonstrates that PDGrapher has a weak dependency on healthy control data and can perform well even when such data are unavailable.

## Discussion

We formulate phenotype-driven lead discovery as a combinatorial prediction problem for therapeutic targets. Given a diseased sample, the goal is to identify genes that a genetic or chemical perturbagen should target to reverse disease effects and shift the sample toward a treated state that matches the distribution of a healthy state. This requires predicting a combination of gene targets, framing the task as combinatorial prediction. To address this, we introduce PDGrapher. Using a diseased cell state represented by a gene expression signature and a proxy causal graph of gene-gene interactions, PDGrapher predicts candidate target genes to transition cells to the desired treated state. PDGrapher includes two modules: a perturbagen discovery module that proposes a set of therapeutic targets based on the diseased and treated states, and a response prediction module that evaluates the effect of applying the predicted perturbagen to the diseased state. Both modules are GNN models that operate on gene-gene networks, which serve as approximations of noisy causal graphs. We use PPI networks and GRNs as two representations of these noisy causal graphs. PDGrapher predicts perturbagens that shift gene expression from diseased to treated states across two evaluation settings (random and leave-cell-out) and 19 datasets involving genetic and chemical interventions. Unlike alternative response prediction methods, which rely on indirect prediction to identify perturbagens, PDGrapher selects candidate gene targets to achieve the desired transformation [37, 38, 51–53].

PDGrapher has the potential to improve therapeutic lead design and expand the search space for perturbagens. It leverages large datasets of genetic and chemical interventions to identify sets of candidate targets that can shift cell line gene expression from diseased to treated states. By selecting sets of therapeutic targets for intervention instead of a single perturbagen, PDGrapher enhances phenotype-driven lead discovery. PDGrapher’s approach to identifying therapeutic targets can enable personalized therapies by tailoring treatments to individual gene expression profiles. Its ability to output multiple genes is especially relevant for diseases where dependencies among several genes affect treatment efficacy and safety.

PDGrapher operates under the assumption that there are no unobserved confounders, a stringent condition that is challenging to validate empirically. Future work could focus on reevaluating and relaxing this assumption. Another limitation lies in the reliance on PPIs and GRNs as proxies for causal gene networks, as these networks are inherently noisy and incomplete [54–56] PDGrapher posits that representation learning can overcome incomplete causal graph approximations. A valuable research direction is to theoretically examine the impact of such approximations, focusing on how they influence the accuracy and reliability of predicted likelihoods. Such analyses could uncover high-level causal variables with therapeutic effects from low-level observations and contribute to reconciling structural causality and representation learning approaches, which generally lack any causal understanding [57]. We performed two experiments to evaluate the robustness of PDGrapher. First, we tested PDGrapher on a PPI network with weighted edges, progressively removing low-confidence edges to assess its performance under increasing network sparsity. Second, we applied PDGrapher to synthetic datasets with varying levels of missing graph components and confounding factors in gene expression data. In both experiments, PDGrapher maintained stable performance. PDGrapher also showed robust performance across PPI networks with different numbers of edges.

Phenotype-driven drug discovery using PDGrapher faces certain limitations, one of which is its reliance on transcriptomic data. Although transcriptomics is broadly applicable, including other data modalities, such as cell morphology screens, could produce more comprehensive models. Cell morphology screens, including cell painting, capture cellular responses by staining organelles and cytoskeletal components, generating image profiles that capture the effects of genetic or chemical perturbations [58, 59]. These screens allow identification of phenotypic signatures that correlate with compound activity, mechanisms of action, and potential off-target effects. The recent release of the JUMP Cell Painting dataset [60] exemplifies how high-content morphological profiling can complement databases such as CMap and LINCS, creating integrated datasets for phenotype-driven discovery. By integrating multimodal data, including phenotypic layers from transcriptomic and image data, it becomes possible to uncover more comprehensive patterns of compound effects [61]. Such integration would broaden the scope of PDGrapher, allowing it to capture wider mechanistic insights and support more effective therapeutic discovery [62, 63].

A limitation of our study is the use of NL20 as a control cell line for A549 [64–66]. Although NL20 is a normal human bronchial epithelial cell line and A549 is a human lung carcinoma cell line derived from the alveolar region, the two cell lines differ in anatomical origin and molecular characteristics. This mismatch could introduce biases in comparative analyses due to variations in baseline gene expression profiles and cellular behaviors. To mitigate this concern, we evaluate PDGrapher’s performance across datasets with and without healthy control data. PDGrapher performs consistently regardless of the inclusion of healthy controls, indicating that its predictions are robust to the absence of matched control cells. Ablation analyses showed that incorporating cycle loss improved PDGrapher’s performance in top target predictions for 5 out of 9 cell lines. Based on this improvement, we included the cycle loss in all experiments. Cycle loss helps maintain the robustness and biological relevance of model predictions. PDGrapher learns to predict drug targets that shift cells from a diseased state to a healthy or treated state. It then uses the diseased gene expression profile and the predicted targets to estimate the gene expression after treatment. This bidirectional approach enforces the fidelity of predicted targets as they must contain sufficient information to reconstruct state B from state A. Cycle loss also acts as a regularizer, penalizing discrepancies between the original input and its reconstruction [67].

PDGrapher is a graph neural network approach for combinatorial prediction of perturbations that transition diseased cells to treated states. By leveraging causal reasoning and representation learning on gene networks, PDGrapher identifies perturbagens necessary to achieve specific phenotypic changes. This approach enables the direct prediction of therapeutic targets that can reverse disease phenotypes, bypassing the need for exhaustive response simulations across large perturbation libraries. Its design and evaluation lay the groundwork for future advances in phenotype-based modeling of therapeutic perturbations by improving the precision and scalability of perturbation prediction methods.

## Method

### Preliminaries

A calligraphic letter 𝒳 indicates a set, an italic uppercase letter *X* denotes a graph, uppercase **X** denotes a matrix, lowercase **x** denotes a vector, and a monospaced letter X indicates a tuple. Uppercase letter X indicates a random variable, and lowercase letter x indicates its corresponding value; bold uppercase **X** denotes a set of random variables, and lowercase letter **x** indicates its corresponding values. We denote *P* (**X**) as a probability distribution over a set of random variables **X** and *P* (**X** = **x**) as the probability of **X** that is equal to the value of **x** under the distribution *P* (**X**). For simplicity, *P* (**X** = **x**) is abbreviated as *P* (**x**). The method section is presented with the terminology of casual inference [68].

### Problem formulation: Combinatorial prediction of targets

Intuitively, given a diseased cell line sample, we would like to predict the set of therapeutic genes that need to be targeted to reverse the effects of disease, that is, the genes that need to be perturbed to shift the cell gene expression state as close as possible to the healthy state. Next, we formalize our problem formulation. Let M =*<* **E, V**, ℱ, *P* (**E**) *>* be an SCM (structural causal models, see the description of related works in supplementary note 4) associated with causal graph *G*, where **E** is a set of exogenous variables affecting the system, **V** are the system variables, ℱ are structural equations encoding causal relations between variables and *P* (**E**) is a probability distribution over exogenous variables. Let 𝒯 = {T_1_, …, T_*m*_} be a dataset of paired healthy and diseased samples (namely disease intervention data), where each element is a triplet T =*<* **v**^*h*^, **U, v**^*d*^ *>* with **v**^*h*^ *∈* [0, 1]^*N*^ being normalized gene expression values of healthy cell line (variable states before perturbation), **V**_**U**_ being the disease-causing perturbed variable (gene) set in **V**, and **v**^*d*^ *∈* [0, 1]^*N*^ being gene expression values of diseased cell line (variable states after perturbation). Our goal is to find, for each sample T =*<* **v**^*h*^, **U, v**^*d*^ *>*, the variable set **U**′ with the highest likelihood of shifting variable states from diseased **v**^*d*^ to healthy **v**^*h*^ state. To increase generality, we refer to the desired variable states as *treated* (**v**^*t*^). Our goal can then be expressed as:

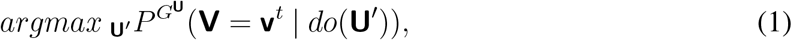

where 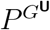 represents the probability on the graph *G* mutilated by perturbations in variables in **U**. Under the assumption of no unobserved confounders, the above interventional probability can be expressed as a conditional probability on the mutilated graph 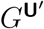 :

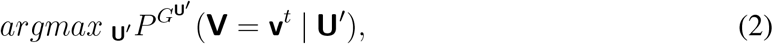

which under the causal Markov condition is:

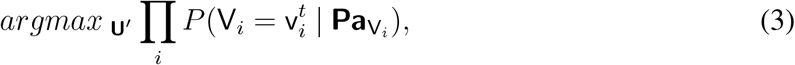

where 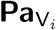 represents parents of variable V_*i*_ according to graph 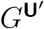 (that is, the mutilated graph upon intervening on variables in **U**′). Here, state of a variable 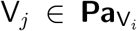 will be equal to an arbitrary value 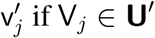. Therefore, intervening on the variable set **U**′ modifies the graph used to obtain conditional probabilities and determine the state of variables in **U**′.

### Problem formulation: a representation learning perspective

In the previous section, we drew on the SCM framework to introduce a generic formulation for the task of combinatorial prediction of therapeutic targets. Instead of approaching the problem from a purely causal inference perspective, we draw upon representation learning to approximate the queries of interest to address the limiting real-world setting of a noisy and incomplete causal graph. Formulating our problem using the SCM framework allows for explicit modeling of interventions and formulation of interventional queries (see the description of related works in supplementary note 4). Inspired by this principled problem formulation, we next introduce the problem formulation using a representation learning paradigm.

We let *G* = (𝒱, ℰ) denote a graph with |𝒱| = *n* nodes and |ℰ| edges, which contains partial information on causal relationships between nodes in 𝒱 and some noisy relationships. We refer to this graph as *proxy causal graph*. Let 𝒯 = {T_1_, …, T_M_} be a dataset with an individual sample being a triplet T =*<* **x**^*h*^, 𝒰, **x**^*d*^ *>* with **x**^*h*^ *∈* [0, 1]^*n*^ being the node states (attributes) of healthy cell sample (before perturbation), 𝒰 being the set of disease-causing perturbed nodes in 𝒱, and **x**^*d*^ *∈* [0, 1]^*n*^ being the node states (attributes) of diseased cell sample (after perturbation). We denote by *G*^𝒰^ = (𝒱, ℰ^𝒰^) the graph resulting from the mutilation of edges in *G* as a result of perturbing nodes in 𝒰 (one graph per perturbagen; we avoid using superindices for simplicity). Here again, we refer to the desired variable states as *treated* (**x**^*t*^). Our goal is then to learn a function:

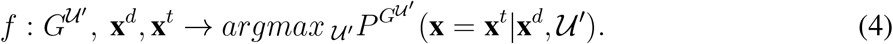

That is, given the graph 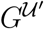, the diseased **x**^*d*^ and treated **x**^*t*^ node states, predicts the combinatorial set of nodes 𝒰′ that if perturbed have the highest chance of shifting the node states to the treated state **x**^*t*^. We note here that 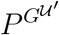 represents probabilities over graph *G*^𝒰^ mutilated upon perturbations in nodes in 𝒰′. Under Causal Markov Condition, we can factorize 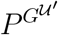 over graph 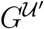 :

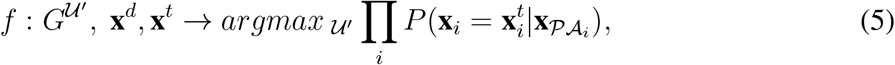

that is, the probability of each node *i* depending only on its parents *PA*_*i*_ in graph 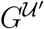.

We assume (i) real-valued node states, (ii) *G* is fixed and given, and (iii) atomic and non-atomic perturbagens (intervening on individual nodes or sets of nodes). Given that the value of each node should depend only on its parents in the graph 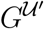, a message-passing framework appears especially suited to compute the factorized probabilities *P*.

In the SCM framework, the conditional probabilities in Equation 3 are computed recursively on the graph, each being an expectation over exogenous variables **E**. Therefore, node states of the previous time point are not necessary. To translate this query into the representation learning realm, we discard the existence of noise variables and directly try to learn a function encoding the transition from an initial state to a desired state. An exhaustive approach to solving Equation 5 would be to search the space of all potential sets of therapeutic targets 𝒰′ and score how effective each is in achieving the desired treated state. This is, indeed, how many cell response prediction approaches can be used for perturbagen discovery [22, 23, 69]. However, with moderately sized graphs, this is highly computationally expensive, if not intractable. Instead, we propose to search for potential perturbagens efficiently with a perturbagen discovery module (*f*_*p*_) and a way to score each potential perturbagen with a response prediction module (*f*_*r*_).

### Relationship to conventional graph prediction tasks

Given that the prediction for each variable is dependent only on its parents in a graph, GNNs appear especially suited for this problem. We can formulate the query of interest under a graph representation learning paradigm as: Given a graph *G* = (𝒱, ℰ), paired sets of node attributes 𝒳 = {**X**_1_, **X**_2_, …, **X**_*m*_} and node labels *Y* = {**Y**_1_, **Y**_2_, …, **Y**_*m*_} where each **Y** = {**y**_1_, …, **y**_*n*_}, with **y**_*i*_ *∈* [0, 1], we aim at training a neural message passing architecture that given node attributes **X**_*i*_ predicts the corresponding node labels **Y**_*i*_. There are, however, differences between our problem formulation and the conventional graph prediction tasks, namely, graph and node classification (summarized in Table S13).

In node classification, a single graph *G* is paired with node attributes **X**, and the task is to predict the node labels **Y**. Our formulation differs in that we have *m* paired sets of node attributes 𝒳 and labels *Y* instead of a single set, yet they are similar in that there is a single graph in which GNNs operate. In graph classification, a set of graphs *G* = {*G*_1_, …, *G*_*m*_} is paired with a set of node attributes 𝒳 = {**X**_1_, **X**_2_, …, **X**_*m*_} and the task is to predict a label for each graph **Y** = {**y**_1_, …, **y**_*m*_}. Here, graphs have a varying structure, and both the topological information and node attributes predict graph labels. In our formulation, a single graph is combined with each node attribute **X**_*i*_, and the goal is to predict a label for each node, not for the whole graph.

### PDGrapher model

PDGrapher is an approach for combinatorial prediction of therapeutic targets composed of two modules. First, a perturbagen discovery module *f*_*p*_ searches the space of potential gene sets to predict a suitable candidate 𝒰′. Next, a response prediction module *f*_*r*_ checks the goodness of the predicted set 𝒰′, that is, how effective intervening on variables in 𝒰′ is to shift node states to the desired treated state **x**^*t*^.

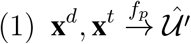

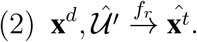

### Model optimization

We optimize our response prediction module *f*_*r*_ using cross-entropy loss on known triplets of disease intervention *<* **x**^*h*^, 𝒰, **x**^*d*^ *>* and treatment intervention *<* **x**^*d*^, 𝒰′, **x**^*t*^ *>*:

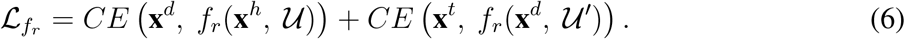

We optimize our intervention discovery module *f*_*p*_ using a cycle loss,ensuring that the response to the predicted intervention set 𝒰′ closely matches the desired treated state (the first part of equation 7). In addition, we provide a supervisory signal for predicting 𝒰′ in the form of cross-entropy loss (the second part of equation 7). So, the total loss is defined as:

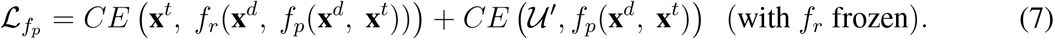

We train *f*_*p*_ and *f*_*r*_ in parallel and implement early stopping separately (see *Experimental setup* for more details). Trained modules *f*_*p*_ are then used to predict, for each diseased cell sample, which nodes should be perturbed (𝒰′) to achieve a desired treated state (Figure 1A).

### Response prediction module

Our response prediction module *f*_*r*_ should learn to map pre-perturbagen node values to post-perturbagen node values through learning relationships between connected nodes (equivalent to learning structural equations in SCMs) and propagating the effects of perturbations downstream in the graph (analogous to the recursive nature of query computations in SCMs).

Given a disease intervention triplet *<* **x**^*h*^, 𝒰, **x**^*d*^ *>*, we propose a neural model operating on a mutilated graph, *G*^𝒰^ where the node attributes are the concatenation of 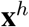and 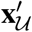, predicting diseased node values **x**^*d*^. The first element is its gene expression value 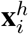, and the second is a perturbation flag: a binary label indicating whether a perturbation occurs at node *i*. So, each node *i* has a two-dimensional attribute vector 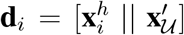. In practice, we embed each node feature into a high-dimensional continuous space by assigning learnable embeddings to each node based on the value of each input feature dimension. Specifically, for each node, we use the binary perturbation flag to assign a d-dimensional learnable embedding, which is different between nodes but shared across samples for each node. To embed the gene expression value 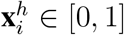, we first calculate thresholds using quantiles to assign the gene expression value into one of the *B* bins. We use the bin index to assign a d-dimensional learnable embedding, which is different between nodes but shared across samples for each node. To increase our model’s representation power, we concatenate a d-dimensional positional embedding (a d-dimensional vector initialized randomly following a normal distribution). Concatenating these three embeddings results in an input node representation of dimensionality 3*d*. For each node *i ∈* 𝒱, an embedding **z**_*i*_ is computed using a graph neural network operating on the node’s neighbors’ attributes. The most general formulation of a GNN layer is:

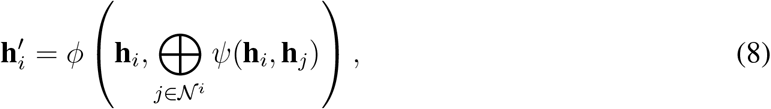

where 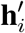 represents the updated information of node *i*, and **h**_*i*_ represents the information of node *i* in the previous layer, with embedded **d**_*i*_ being the input to the first layer. *ψ* is a *MESSAGE* function, EB an *AGGREGATE* function (permutation-invariant), and *ϕ* is an *UPDATE* function. We obtain an embedding **z**_*i*_ for node *i* by stacking *K* GNN layers. The node embedding **z**_*i*_ *∈* ℝ is then passed to a multilayer feedforward neural network to obtain an estimate of the values of the post-perturbation nodes **x**^*d*^.

### Perturbation discovery module

Our perturbagen prediction module *f*_*p*_ should learn the nodes in the graph that should be perturbed to shift the node states (attributes) from diseased **x**^*d*^ to the desired treated state **x**^*t*^. Given a triplet *<* **x**^*d*^, 𝒰′, **x**^*t*^ *>*, we propose a neural model operating on graph 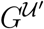 with node features **x**^*d*^ and **x**^*t*^ that predicts a ranking for each node where the top *P* ranked nodes should be predicted as the nodes in 𝒰′. Each node *i* has a two-dimensional attribute vector: 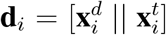. In practice, we represent these binary features in a continuous space using the same approach as described for our response prediction module *f*_*r*_.

For each node *i ∈* 𝒱, an embedding **z**_*i*_ is computed using a graph neural network operating on the node’s neighbors’ attributes. We obtain an embedding **z**_*i*_ for node *i* by stacking *K* GNN layers. The node embedding **z**_*i*_ *∈* ℝ is then passed to a multilayer feedforward neural network to predict a real-valued number for node *i*.

### Model implementation and training

We implement PDGrapher using PyTorch 1.10.1 [70] and the Torch Geometric 2.0.4 Library [71]. The implemented architecture yields a neural network with the following hyperparameters: number of GNN layers and number of prediction layers. We set the number of prediction layers to two and performed a grid search over the number of GNN layers (1-3 layers). We train our model using a 5-fold cross-validation strategy and report PDGrapher’s performance resulting from the best-performing hyperparameter setting.

### Further details on statistical analysis

We next outline the evaluation setup, baseline models, and statistical tests used to evaluate PDGrapher. We evaluate the performance of PDGrapher against a set of baselines:

- **Random baseline:** Given a sample T =*<* **x**^*d*^, 𝒰′, **x**^*t*^ *>*, the random baseline returns *N* random genes as the prediction of target genes in 𝒰′, where *N* is the number of genes in 𝒰′.
- **Cancer genes:** Given a sample T =*<* **x**^*d*^, 𝒰′, **x**^*t*^ *>*, the cancer genes baseline returns the top *N* genes from an ordered list where the first *M* genes are disease-associated (cancer-driver genes). The remaining genes are ranked randomly, and *N* is the number of genes in 𝒰′. The processing of cancer genes is described in the section on *Disease-genes information* in supplementary Note 3.
- **Cancer drug targets:** Given a sample T =*<* **x**^*d*^, 𝒰′, **x**^*t*^ *>*, the cancer targets baseline returns the top *N* genes from an ordered list where the first *M* genes are cancer drug targets and the remaining genes are ranked randomly, and *N* is the number of genes in 𝒰′. The processing of drug target information is described in sections on *Drug-targets information* and *Cancer drug and target information* in supplementary Note 3.
- **Perturbed genes:** Given a sample T =*<* **x**^*d*^, 𝒰′, **x**^*t*^ *>*, the perturbed genes baseline returns the top *N* genes from an ordered list where the first *M* genes are all perturbed genes in the training set and the remaining genes are ranked randomly, and *N* is the number of genes in 𝒰′.
- **scGen [22]:** scGen is a widely-used gold-standard latent variable model for response prediction [72–75]. Given a set of observed cell types in control and perturbed state, scGen predicts the response of a new cell type to the perturbagen seen in training. To utilize scGen as a baseline, we first fit it to our LINCS gene expression data for each dataset type to predict response to perturbagens, training one model per perturbagen (chemical or genetic). Then, given a sample of paired diseased-treated cell line states, T =*<* **x**^*d*^, 𝒰′, **x**^*t*^ *>*, we compute the response of cell line with gene expression 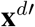 to all perturbagens. The predicted perturbagen is that whose predicted response is closest to **x**^*t*^ in *R*^2^ score. Since scGen trains one model per perturbagen, it needs an exhaustively long training time for datasets with a large number of perturbagens, especially in the leave-cell-out setting. Therefore, we set the maximum training epochs to 100 and only conducted leave-cell-out tests for one split of data for scGEN.
- **Biolord [34]**: Biolord can predict perturbagen response for both chemical and genetic datasets. We followed the official tutorial from the Biolord GitHub repository: https://github.com/nitzanlab/biolord, using the recommended parameters. To prevent memory and quota errors, we implemented two filtering steps: 1) Instead of storing the entire response gene expression (rGEX) matrix of all input (control) cells for each perturbagen, we only store a vector of the averaged rGEX of the input cells per perturbagen, which is necessary for calculating *R*^2^ for evaluation; 2) During prediction, if the number of control cells exceeds 10,000, we randomly down-sample the control cells to 10,000. Similar to scGEN, we predict the responses gene expression **x**^*d′*^ for all perturbagens and use them to calculate *R*^2^ to get the rank of predicted perturbagens.
- **ChemCPA [33]**: ChemCPA is specifically designed for chemical perturbation. We followed the official tutorials on GitHub for running this model (https://github.com/theislab/chemCPA), with all parameters set following the authors’ recommendations. Data processing was also conducted using the provided scripts. We constructed drug embedding using RDKit with canonical SMILES sequences, as this is the default setting in the model and the tutorial. Since the original ChemCPA model lacks functionality to obtain the predicted rGEX for each drug (averaging over the dosages), we developed a custom script to perform this task. These predictions, **x**^*d′*^, were subsequently used for calculating *R*^2^ to get the rank of predicted perturbagens.
- **GEARS [35]**: GEARS is capable of predicting perturbagen responses for genetic perturbation datasets, specifically for predicting the rGEX to unseen perturbagens. However, it is limited to predicting only those genes that are present in the gene network used as prior knowledge for model training. Additionally, GEARS cannot process perturbagens with only one sample, so we filtered the data accordingly. We followed the official tutorial from the GEARS GitHub repository (https://github.com/snap-stanford/GEARS), using the recommended parameters. After confirming with the authors, we established that GEARS is suitable only for within-cell-line prediction. Consequently, our experiments with GEARS were conducted exclusively within this scenario.
- **CellOT [27]**: CellOT is capable of working with both chemical and genetic datasets. We ran this model by following the official tutorial from GitHub (https://github.com/bunnech/cellot), ensuring that all parameters were set according to the provided guidelines. Due to CellOT’s limitation in processing perturbagens with small sample sizes, we filtered the data to retain only those perturbagens with more than five samples or cells. We then used the predicted response gene expression 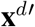 to calculate *R*^2^ and the predicted perturbagen ranks. Similar to scGEN, CellOT trains one model per perturbagen, which results in an exhaustively long training time for datasets with a large number of perturbagens. This issue becomes even more significant when doing leave-cell-out evaluations. Therefore, for this method, we set the maximum training epochs to 100 and only conduct one split in leave-cell-out tests.

### Dataset splits and evaluation settings

We evaluate PDGrapher and competing methods on two different settings:

- **Systematic random dataset splits:** For each cell line, the dataset is split randomly into train and test sets to measure our model performance in an IID setting.
- **Leave-cell-out dataset splits:** To measure model performance on unseen cell lines, we train our model with random splits on one cell line and test on a new cell line. Specifically, for chemical perturbation data, we train a model for each random split per cell line and test it on the entire dataset of the remaining eight cell lines. For genetic data, we train a model for each random split per cell line and test it on the entire dataset of the remaining nine cell lines. For example, with nine cell lines with chemical perturbation (A549, MDAMB231, BT20, VCAP, MCF7, PC3, A375, HT29, and HELA), we conducted an experiment where each split of cell line A549 was used as the training set, and the trained model was tested on the remaining eight cell lines (MDAMB231, BT20, VCAP, MCF7, PC3, A375, HT29, and HELA). Similarly, for cell line MDAMB231, we trained the model on each split of it and tested the model on the other eight cell lines (A549, BT20, VCAP, MCF7, PC3, A375, HT29, and HELA). This process was repeated for all cell lines, providing a comprehensive evaluation of PDGrapher and all competing methods.

### Evaluation setup

For all dataset split settings, our model is trained using 5-fold cross-validation, and metrics are reported as the average on the test set. Within each fold, we further split the training set into training and validation sets (8:2) to perform early stopping: we train the model on the training set until the validation loss has not decreased at least 10^*−*5^ for 15 continuous epochs.

### Evaluation metrics

We report average sample-wise *R*^2^ score, and average perturbagen-wise *R*^2^ score to measure performance in the prediction of **x**^*t*^. The sample-wise *R*^2^ score is computed as the square of Pearson correlation between the predicted sample 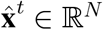 and real sample **x**^*t*^ *∈* ℝ^*N*^. The perturbagen-wise *R*^2^ score is adopted from scGen. It is computed as the square of Pearson correlation of a linear least-squares regression between a set of predicted treated samples 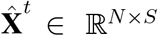 and a set of real treated samples **X**^*t*^ *∈* ℝ^*N ×S*^ for the same perturbagen. Higher values indicate better performance in predicting the treated sample **x**^*t*^ given the diseased sample **x**^*d*^ and predicted perturbagen. This is used for evaluating the performance of response prediction. For evaluating perturbagen discovery, when the competing methods cannot predict perturbagen ranks for chemical perturbation data, we first calculate the rank of drugs based on the *R*^2^ score. We then build a target gene rank from the drug rank by substituting the drugs with their target genes acquired from DrugBank [76] (accessed in November 2022) (see details in supplementary Note 3). A single drug can have multiple target genes, which we place in the rank in random order. Since some methods cannot predict unseen drugs, their predicted target gene lists are often short, introducing bias in evaluation. To address this, we shuffle the missing target genes and attach them to the predicted ranks to create a complete rank. For genetic perturbation data, we directly obtain the target gene rank from the results, then attach the shuffled missing genes to the rank.

To evaluate the performance of our model in ranking predicted therapeutic targets, we use the Normalized Discounted Cumulative Gain (nDCG), a widely-used metric in information retrieval adapted for our setting. The raw DCG score is computed by summing the relevance of each correct target based on its rank in the predicted list, with relevance weighted by a logarithmic discount factor to prioritize higher-ranked interventions. The gain function is defined as 1 *− ranking/N*, ensuring that the score reflects the quality of the ranking relative to the total number of nodes in the system. To ensure comparability across datasets or experiments with different numbers of correct interventions, DCG is normalized by the Ideal DCG (IDCG), which represents the maximum possible score for a perfect ranking. This results in nDCG values in the range [0, 1], where higher values indicate better ranking performance and alignment with the ground truth. This metric is particularly suited for our task as it emphasizes the accuracy of top-ranked interventions while accounting for the diminishing importance of lower-ranked predictions.

In addition, we report the proportion of test samples for which the predicted therapeutic targets set has at least one overlapping gene with the ground-truth therapeutic targets set (denoted as the percentage of accurately predicted samples). We also calculated the ratio of correct therapeutic targets that appeared in the top 1, top 10, and top 100 predicted therapeutic targets in the predicted rank, denoted as recall@1, recall@10, and recall@100, respectively.

To assess the overall performance across all experiments and metrics, we calculated an aggregated metric, averaging all metric values for each method.

### Statistical test

In the benchmarking experiments, we performed a one-tailed pairwise t-test to evaluate whether PDGrapher significantly outperforms the competing methods. For other experiments, such as ablation studies, we employed a two-tailed t-test to determine whether there is a significant difference in performance between the two models. A significance threshold of 0.05 was used for all tests. P-values of perturbagen discovery and response prediction tests are presented in the source data of the figures.

### Ablation studies

In the ablation study, we evaluated PDGrapher by optimizing it with only the supervision loss (PDGrapher-Super) and with only the cycle loss (PDGrapher-Cycle) across all chemical datasets. We then compared the perturbagen prediction performance of these sub-models with that of PDGrapher (PDGrapher-SuperCycle). To train PDGrapher-Super and PDGrapher-Cycle, for each cell line, we set the number of layers to that which was found optimal for the validation set in the random splitting setting for PDGrapher-SuperCycle.

### Sensitivity studies

To test the sensitivity of PDGrapher on protein-protein interactions (PPIs), we utilized data from STRING (string-db.org), which provides a confidence score for each edge. The method for acquiring and preprocessing the PPIs from STRING is detailed in supplementary Note 3. For the sensitivity tests, we selected two cell lines: the chemical dataset MDAMB231 and the genetic dataset MCF7. For each cell line, we processed the data using the five PPI networks described in the supplementary Note 3. We optimized PDGrapher using 5-fold cross-validation as described in the *Evaluation setup* section and optimized the number of GNN layers using the validation set in each split.

### Synthetic datasets

We generated three synthetic datasets: **(1) dataset with missing components removing bridge edges:** this dataset is generated by progressively removing bridge edges from the existing PPI network. Bridge edges are those whose removal disconnects parts of the network. We vary the fraction of bridge edges removed in increments (from [0,0.1,…0.6]), and for each fraction, we create a new edge list representing the modified network (Table S5). This process ensures that different levels of network sparsity are introduced, affecting the overall structure and connectivity. We pair these networks with gene expression data from Chemical-PPI-Breast-MDAMB231. **(2) Dataset with missing components removing random edges:** this dataset is generated by progressively removing random edges from the existing PPI network. We vary the fraction of bridge edges removed in increments (from [0, 0.1, 0.6]), and for each fraction, we create a new edge list representing the modified network. The number of remaining directed edges in the network upon random edge removal are 273,319; 242,912; 212,525; 182,177; 151,811; 121,472. **(3) Dataset with latent confounder noise:** our starting point is the Chemical-PPI-Breast-MDAMB231 dataset. The synthetic datasets were constructed with varying levels of confounding bias introduced into the gene expression data. To simulate latent confounder effects, Gaussian noise with distinct means and variances was progressively added to random subsets of genes. Genes were grouped into 50 predefined subsets, each representing a latent confounder group. For each group, a Gaussian distribution was defined, with the mean drawn randomly from a uniform distribution in the range [0.5, 0.5] and the standard deviation from [0.1, 0.5]. A fraction ([0.2, 0.4, 0.6, 0.8, 1]) of these subsets was randomly selected for perturbation, and for each gene in these subsets, its expression value was incremented by a value sampled from the respective Gaussian distribution. The perturbed gene expression values were then clamped between 0 and 1 to ensure validity. This strategy ensures that different latent biases are introduced globally to gene expression patterns while maintaining controlled variability. We pair the noisy version of the gene expression data with the global unperturbed PPI network.

### Network proximity between predicted and true perturbagens

Let *P* be the set of predicted therapeutic targets, *R* be the set of ground truth therapeutic targets, and *spd*(*p, r*) be the shortestpath distance between nodes in *P* and *R*. We measure the closest distance between *P* and *R* as:

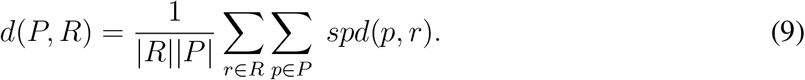

As part of our performance analyses, we measure the network proximity of PDGrapher and competing methods. We compared the distributions of network proximity values using a MannWhitney U test, along with a rank-biserial correlation to measure effect size. To assess the uncertainty of effect sizes, we performed bootstrapping with 1,000 resamples to estimate 95% confidence intervals.

## Data availability

Processed data used in this paper, including the cell line gene expression dataset, protein-protein interaction network, drug targets, and disease-associated genes, are available via the project website at https://zitniklab.hms.harvard.edu/projects/PDGrapher or directly at https://zenodo.org/records/15375990 [77] and https://zenodo.org/records/15390483 [78]. The raw protein-protein interaction network data was obtained from https://downloads.thebiogrid.org/File/BioGRID/Release-Archive/BIOGRID-3.5.186/BIOGRID-MV-Physical-3.5.186.tab3.zip, https://www.science.org/doi/suppl/10.1126/science.1257601/supplfile/datasetss1-s4.zip, and http://www.interactome-atlas.org/data/HuRI.tsv. Raw gene expression datasets were obtained from https://clue.io/releases/data-dashboard. Disease-associated genes were obtained from COSMIC at https://cancer.sanger.ac.uk/celllines/archive-download#:~:text=Complete%20mutation%20data and https://cancer.sanger.ac.uk/cosmic/curation. Drug targets were extracted from DrugBank at https://go.drugbank.com/releases/5-1-9, and a list of cancer drugs was obtained from NCI at https://www.cancer.gov/about-cancer/treatment/types/targeted-therapies/approved-drug-list. STRING data for building PPI graphs were downloaded from https://stringdb-downloads.org/download/protein.physical.links.detailed.v12.0.txt.gz in version 12.0.

## Code availability

Python implementation of PDGrapher is available at https://github.com/mims-harvard/PDGrapher [79].

## Acknowledgements

We would like to thank Domen Mohorcic for his help with the PDGrapher codebase. We gratefully acknowledge the support of NIH R01-HD108794, NSF CAREER 2339524, US DoD FA8702-15-D-0001, ARPA-H BDF program, awards from Chan Zuckerberg Initiative, Bill & Melinda Gates Foundation INV-079038, Amazon Faculty Research, Google Research Scholar Program, AstraZeneca Research, Roche Alliance with Distinguished Scientists, Sanofi iDEA-iTECH, Pfizer Research, John and Virginia Kaneb Fellowship at Harvard Medical School, Biswas Computational Biology Initiative in partnership with the Milken Institute, Harvard Medical School Dean’s Innovation Fund for the Use of Artificial Intelligence, Harvard Data Science Initiative, and Kempner Institute for the Study of Natural and Artificial Intelligence at Harvard University. I.H. was supported, in part, by the Summer Institute in Biomedical Informatics at Harvard Medical School. M.B. and G.G. were supported by the ERC-Consolidator Grant No. 724228 (LEMAN). Any opinions, findings, conclusions or recommendations expressed in this material are those of the authors and do not necessarily reflect the views of the funders.

## Authors contribution

G.G. retrieved, processed, and analyzed gene expression data, protein-protein interaction network data, and disease gene datasets. I.H. retrieved, processed, and analyzed drug target data and processed gene regulatory networks. G.G. and X.L. developed, implemented, and benchmarked PDGrapher and performed detailed analyses of PDGrapher. G.G., X.L., K.V., M.B., and M.Z. designed the study. G.G., X.L., I.H., and M.Z. wrote the manuscript.

## Competing interests

G.G. is currently employed by Genentech, Inc. and I.H. was employed by Merck & Co., Inc. during the study. The remaining authors declare no competing interests.

## Notes

https://zitniklab.hms.harvard.edu/projects/PDGrapher

https://github.com/mims-harvard/PDGrapher

### Summary of Updates

Formatting changes: updated main figures, shortened Method section, moved some supplementary figures to extended data.

## References

1. Vincent, F. et al./person-group>. Phenotypic drug discovery: recent successes, lessons learned and new directions (2022).

2. Moffat, J. G., Vincent, F., Lee, J. A., Eder, J. & Prunotto, M. Opportunities and challenges in phenotypic drug discovery: An industry perspective. Nature Reviews Drug Discovery 16, 531–543 (2017).

3. Minikel, E. V., Painter, J. L., Dong, C. C. & Nelson, M. R. Refining the impact of genetic evidence on clinical success. Nature 1–6 (2024).

4. Druker, B. J. et al./person-group>. Effects of a selective inhibitor of the abl tyrosine kinase on the growth of bcr–abl positive cells. Nature Medicine 2, 561–566 (1996).

5. Bange, J., Zwick, E. & Ullrich, A. Molecular targets for breast cancer therapy and prevention. Nature Medicine 7, 548–552 (2001).

6. Swinney, D. C. & Anthony, J. How were new medicines discovered? Nature Reviews Drug Discovery 10, 507–519 (2011).

7. Musa, A. et al./person-group>. A review of connectivity map and computational approaches in pharmacogenomics. Briefings in Bioinformatics 19, 506–523 (2018).

8. Davies, J. C., Alton, E. W. & Bush, A. Cystic fibrosis. British Medical Journal 335, 1255–1259 (2007).

9. Van Goor, F. et al./person-group>. Rescue of CF airway epithelial cell function in vitro by a CFTR potentiator, VX-770. Proceedings of the National Academy of Sciences of the United States of America 106, 18825–18830 (2009).

10. Van Goor, F. et al./person-group>. Correction of the F508del-CFTR protein processing defect in vitro by the investigational drug VX-809. Proceedings of the National Academy of Sciences of the United States of America 108, 18843–18848 (2011).

11. Keenan, A. B. et al./person-group>. Connectivity Mapping: Methods and Applications (2019).

12. Keenan, A. B. et al./person-group>. The Library of Integrated Network-based Cellular Signatures (LINCS) NIH Program: System-level Cataloging of Human Cells Response to Perturbations. Cell System 6, 13–24 (2018).

13. Lamb, J. et al./person-group>. The Connectivity Map: Using Gene-Expression Signatures to Connect Small Molecules, Genes, and Disease. science 313, 1929–1935 (2006).

14. Samart, K., Tuyishime, P., Krishnan, A. & Ravi, J. Reconciling multiple connectivity scores for drug repurposing. Briefings in Bioinformatics 22 (2021).

15. Guney, E. Reproducible drug repurposing: When similarity does not suffice. Pacific Symposium on Biocomputing 0, 132–143 (2017).

16. Chen, B. et al./person-group>. Reversal of cancer gene expression correlates with drug efficacy and reveals therapeutic targets. Nature Communications 8 (2017).

17. Chen, B. et al./person-group>. Computational Discovery of Niclosamide Ethanolamine, a Repurposed Drug Candidate That Reduces Growth of Hepatocellular Carcinoma Cells In Vitro and in Mice by Inhibiting Cell Division Cycle 37 Signaling. Gastroenterology 152, 2022–2036 (2017).

18. Pessetto, Z. Y. et al./person-group>. In silico and in vitro drug screening identifies new therapeutic approaches for Ewing sarcoma. Oncotarget 8, 4079–4095 (2016).

19. Morselli Gysi, D. et al. Network medicine framework for identifying drug-repurposing opportunities for COVID-19. Proceedings of the National Academy of Sciences of the United States of America 118 (2021).

20. Zhu, J. et al./person-group>. Prediction of drug efficacy from transcriptional profiles with deep learning. Nature Biotechnology 2021 39:11 39, 1444–1452 (2021).

21. Pham, T. H., Qiu, Y., Zeng, J., Xie, L. & Zhang, P. A deep learning framework for high-throughput mechanism-driven phenotype compound screening and its application to COVID-19 drug repurposing. Nature Machine Intelligence 2021 3:3 3, 247–257 (2021).

22. Lotfollahi, M., Wolf, F. A. & Theis, F. J. scGen predicts single-cell perturbation responses. Nature Methods 2019 16:8 16, 715–721 (2019).

23. Hetzel, L. et al./person-group>. Predicting Cellular Responses to Novel Drug Perturbations at a Single-Cell Resolution.

24. Hauser, A. & Bühlmann, P. Two optimal strategies for active learning of causal models from interventional data. International Journal of Approximate Reasoning 55, 926–939 (2014).

25. Ghassami, A. E., Salehkaleybar, S., Kiyavash, N. & Bareinboim, E. Budgeted experiment design for causal structure learning. 35th International Conference on Machine Learning, ICML 2018 4, 2788–2801 (2018).

26. Agrawal, R., Squires, C., Yang, K., Shanmugam, K. & Uhler, C. ABCD-strategy: Budgeted experimental design for targeted causal structure discovery. AISTATS 2019 - 22nd International Conference on Artificial Intelligence and Statistics 89 (2020).

27. Bunne, C. et al./person-group>. Learning single-cell perturbation responses using neural optimal transport. Nature Methods 20, 1759–1768 (2023).

28. Barr, M. P. et al./person-group>. Vascular endothelial growth factor is an autocrine growth factor, signaling through neuropilin-1 in non-small cell lung cancer. Molecular cancer 14, 1–16 (2015).

29. Riquelme, E. et al./person-group>. Vegf/vegfr-2 upregulates ezh2 expression in lung adenocarcinoma cells and ezh2 depletion enhances the response to platinum-based and vegfr-2–targeted therapy. Clinical Cancer Research 20, 3849–3861 (2014).

30. Huynh-Thu, V. A., Irrthum, A., Wehenkel, L. & Geurts, P. Inferring regulatory networks from expression data using tree-based methods. PloS one 5, e12776 (2010).

31. Yuan, B. et al./person-group>. CellBox: interpretable machine learning for perturbation biology with application to the design of cancer combination therapy. Cell Systems 12, 128–140 (2021).

32. Kamimoto, K. et al./person-group>. Dissecting cell identity via network inference and in silico gene perturbation. Nature 614, 742–751 (2023).

33. Hetzel, L. et al./person-group>. Predicting cellular responses to novel drug perturbations at a single-cell resolution. Advances in Neural Information Processing Systems 35, 26711–26722 (2022).

34. Piran, Z., Cohen, N., Hoshen, Y. & Nitzan, M. Disentanglement of single-cell data with biolord. Nature Biotechnology 1–6 (2024).

35. Roohani, Y., Huang, K. & Leskovec, J. Predicting transcriptional outcomes of novel multigene perturbations with gears. Nature Biotechnology 42, 927–935 (2024).

36. Barabási, A. L., Gulbahce, N. & Loscalzo, J. Network Medicine: A Network-based Approach to Human Disease. Nature reviews. Genetics 12, 56 (2011).

37. Ruiz, C., Zitnik, M. & Leskovec, J. Identification of disease treatment mechanisms through the multiscale interactome. Nature Communications 12, 1796 (2021).

38. Eyuboglu, S., Zitnik, M. & Leskovec, J. Mutual interactors as a principle for phenotype discovery in molecular interaction networks. In Pacific Symposium on Biocomputing, 61–72 (2023).

39. Gu, Z. et al./person-group>. Role of duplicate genes in genetic robustness against null mutations. Nature 421, 63–66 (2003).

40. Deutscher, D., Meilijson, I., Kupiec, M. & Ruppin, E. Multiple knockout analysis of genetic robustness in the yeast metabolic network. Nature Genetics 38, 993–998 (2006).

41. Deutscher, D., Meilijson, I., Schuster, S. & Ruppin, E. Can single knockouts accurately single out gene functions? BMC Systems Biology 2 (2008).

42. Ochoa, D. et al./person-group>. The next-generation open targets platform: reimagined, redesigned, rebuilt. Nucleic acids research 51, D1353–D1359 (2023).

43. Cai, Q. et al./person-group>. The role of dexmedetomidine in tumor-progressive factors in the perioperative period and cancer recurrence: a narrative review. Drug Design, Development and Therapy 2161–2175 (2023).

44. Gainor, J. F. et al./person-group>. Pralsetinib for ret fusion-positive non-small-cell lung cancer (arrow): a multi-cohort, open-label, phase 1/2 study. The lancet oncology 22, 959–969 (2021).

45. Faraat Ali, G. C., Kumari Neha. Pralsetinib: chemical and therapeutic development with fda authorization for the management of ret fusion-positive non-small-cell lung cancers. Arch Pharm Res. 45, 309–327 (2022).

46. Wu, J. et al./person-group>. Expression and potential molecular mechanism of top2a in metastasis of non-small cell lung cancer. Scientific Reports 14, 12228 (2024).

47. Pautier, P. et al./person-group>. Doxorubicin–trabectedin with trabectedin maintenance in leiomyosarcoma. New England Journal of Medicine 391, 789–799 (2024).

48. Hartwell, L. H., Hopfield, J. J., Leibler, S. & Murray, A. W. Hartwell Et Al 199. Nature 402, C47–C52 (1999).

49. Menche, J. et al./person-group>. Uncovering disease-disease relationships through the incomplete interactome. Science 347, 841 (2015).

50. Szklarczyk, D. et al./person-group>. The string database in 2023: protein–protein association networks and functional enrichment analyses for any sequenced genome of interest. Nucleic Acids Research 51, D638–D646 (2023).

51. Bagherian, M. et al./person-group>. Machine learning approaches and databases for prediction of drug-target interaction: A survey paper. Briefings in Bioinformatics 22, 247–269 (2021).

52. Pan, J. et al./person-group>. Sparse dictionary learning recovers pleiotropy from human cell fitness screens. Cell Systems 13, 286–303 (2022).

53. Huang, K. et al./person-group>. Deeppurpose: a deep learning library for drug–target interaction prediction. Bioinformatics 36, 5545–5547 (2020).

54. Hart, G. T., Ramani, A. K. & Marcotte, E. M. How complete are current yeast and human protein-interaction networks? Genome Biology 7 (2006).

55. Stumpf, M. P. et al./person-group>. Estimating the size of the human interactome. Proceedings of the National Academy of Sciences of the United States of America 105, 6959–6964 (2008).

56. Zitnik, M., Sosic?, R., Feldman, M. W. & Leskovec, J. Evolution of resilience in protein interactomes across the tree of life. Proceedings of the National Academy of Sciences 116, 4426–4433 (2019).

57. Schölkopf, B. et al./person-group>. Toward causal representation learning. Proceedings of the IEEE 109, 612–634 (2021).

58. Gustafsdottir, S. M. et al./person-group>. Multiplex cytological profiling assay to measure diverse. PLoS ONE 8, 1–7 (2013).

59. Bray, M.-A. et al./person-group>. Cell Painting, a high-content image-based assay for morphological profiling using multiplexed fluorescent dyes. Physiology & behavior 176, 139–148 (2017).

60. Chandrasekaran, S. N. et al./person-group>. JUMP Cell Painting dataset: morphological impact of 136,000 chemical and genetic perturbations. bioRxiv (2023).

61. Akbarzadeh, M. et al./person-group>. Morphological profiling by means of the Cell Painting assay enables identification of tubulin-targeting compounds. Cell Chemical Biology 29, 1053–1064 (2022).

62. Pruteanu, L. L. & Bender, A. Using Transcriptomics and Cell Morphology Data in Drug Discovery: The Long Road to Practice. ACS Medicinal Chemistry Letters 14, 386–395 (2023).

63. Huang, K. et al./person-group>. Artificial intelligence foundation for therapeutic science. Nature Chemical Biology 18, 1033–1036 (2022).

64. Thomas, K. J. & Jacobson, M. R. Defects in mitochondrial fission protein dynamin-related protein 1 are linked to apoptotic resistance and autophagy in a lung cancer model (2012).

65. Le, C. F., Yusof, M. Y. Y., Hassan, H. & Sekaran, S. D. In vitro properties of designed antimicrobial peptides that exhibit potent antipneumococcal activity and produce synergism in combination with penicillin. Scientific Reports 5, 9761 (2015).

66. Zhao, S., Song, P., Zhou, G., Zhang, D. & Hu, Y. Mettl3 promotes the malignancy of non-small cell lung cancer by n6-methyladenosine modifying sfrp2. Cancer Gene Therapy 30, 1094–1104 (2023).

67. Zhu, J.-Y., Park, T., Isola, P. & Efros, A. A. Unpaired image-to-image translation using cycle-consistent adversarial networks. In Proceedings of the IEEE International Conference on Computer Vision (ICCV) (2017).

68. Neal, B. Introduction to causal inference. Course Lecture Notes (draft) 132 (2020).

69. Lotfollahi, M. et al./person-group>. Predicting cellular responses to complex perturbations in high-throughput screens. Molecular systems biology e11517 (2023).

70. Paszke, A. et al./person-group>. Automatic differentiation in PyTorch. Tech. Rep.

71. Fey, M. & Lenssen, J. E. Fast Graph Representation Learning with PyTorch Geometric. ICLR Workshop on Representation Learning on Graphs and Manifolds (2019).

72. Heumos, L. et al./person-group>. Best practices for single-cell analysis across modalities. Nature Reviews Genetics 2023 1–23 (2023).

73. Gayoso, A. et al./person-group>. A Python library for probabilistic analysis of single-cell omics data. Nature Biotechnology 2022 40:2 40, 163–166 (2022).

74. Jovic, D. et al./person-group>. Single-cell RNA sequencing technologies and applications: A brief overview. Clinical and Translational Medicine 12, e694 (2022).

75. Luecken, M. D. et al./person-group>. Benchmarking atlas-level data integration in single-cell genomics. Nature Methods 2021 19:1 19, 41–50 (2021).

76. Wishart, D. S. et al./person-group>. DrugBank 5.0: a major update to the DrugBank database for 2018. Nucleic Acids Research 46, D1074–D1082 (2018).

77. Gonzalez, G. et al./person-group>. Combinatorial prediction of therapeutic perturbations using causally-inspired neural networks (genetic data). Datasets. Zenodo 10.5281/zenodo.15375990 (2025).

78. Gonzalez, G. et al./person-group>. Combinatorial prediction of therapeutic perturbations using causally-inspired neural networks (chemical data). Datasets. Zenodo 10.5281/zenodo.15390483 (2025).

79. Gonzalez, G. et al./person-group>. PDGrapher. GitHub https://github.com/mims-harvard/PDGrapher/tree/main (2025).

