## Supplementary data for "Combinatorial prediction of therapeutic perturbations using causally-inspired neural networks"

### Contents

#### 1. Supplementary Notes

**Supplementary Note 1.** PDGrapher illuminates mode of action of chemical perturbagens.
**Supplementary Note 2.** Scoring system for target prediction.
**Supplementary Note 3.** Data generation and preprocessing.
**Supplementary Note 4.** Related works.

#### 2. Supplementary Tables

**Supplementary Table 1.** Number of seen and unseen perturbagens per cell line and across dataset splits, considering chemical perturbations data.
**Supplementary Table 2.** Number of unseen perturbagens in each dataset split across cell lines and considering chemical perturbation data.
**Supplementary Table 3.** Number of seen and unseen perturbagens per cell line per dataset split with genetic perturbation data.
**Supplementary Table 4.** Number of unseen perturbagens in each dataset split across cell lines and considering genetic perturbation data.
**Supplementary Table 5.** Details of synthetic experiments.
**Supplementary Table 6.** PDGrapher predicts targets that are closer in the network to ground-truth targets than what would be expected by change in Chemical PPI datasets in splits with new samples.
**Supplementary Table 7.** PDGrapher predicts targets that are closer in the network to ground-truth targets than what would be expected by change in Chemical PPI datasets in splits with new samples and cell lines.
**Supplementary Table 8.** Scoring system of evidence for targets (Due to extended size of this table, we provide it as a separate XLSX file.)
**Supplementary Table 9.** Cell line selection in chemical LINCS dataset.
**Supplementary Table 10.** Cell line selection in genetic LINCS dataset.
**Supplementary Table 11.** Sample size and number of perturbation of the genetic perturbation datasets from CMAP.
**Supplementary Table 12.** Sample size and number of perturbation of the chemical perturbation datasets from CMAP.
**Supplementary Table 13**. Problem formulation of PDGrapher.

#### 3. Supplementary Figures

**Supplementary Figure 1.** The performance of response prediction across nine cell lines under chemical perturbation.

**Supplementary Figure 2.** PDGrapher predicts genetic perturbagens to shift cells from diseased to treated states in unseen cell lines.

**Supplementary Figure 3.** Network visualization of predicted and ground-truth targets can aid in ellucidation of mechanism of action of drugs.

**Supplementary Figure 4.** PDGrapher predicts genetic perturbagens and response gene expression to genetic perturbagens using cell type specific GRNs.

**Supplementary Figure 5.** PDGrapher predicts chemical perturbagens and response gene expression to chemical perturbagens using cell type specific GRNs.

**Supplementary Figure 6.** Figure S6: PDGrapher predicts chemical perturbagens to shift cells from diseased to treated states without using disease intervention data.

**Supplementary Figure 7.** PDGrapher predicts genetic perturbagens to shift cells from diseased to treated states without using disease intervention data.

**Supplementary Figure 8.** Ablation studies for loss functions of PDGrapher.

#### Supplementary Notes

##### Note 1. PDGrapher illuminates mode of action of chemical perturbagens: a case study for Vorinostat and Sorafenib predicted by PDGrapher for lung cancer

To demonstrate PDGrapher's ability to illuminate the mechanism of action of therapeutic perturbagens, we analyze PDGrapher's predictions for Vorinostat and Sorafenib in Chemical-PPI-Lung-A549 in the random splitting setting. We visualize ground truth and predicted combinations of therapeutic targets together with their one-hop neighbors in the protein interaction network using Gephi v0.10[1] and the ForceAtlas graphical layout (Figure S3). The network structure reveals distinct community clusters, identified by Gephi's modularity algorithm, with clear visual delineation based on the predicted and known targets. This analysis demonstrates the model's ability to correctly identify several key therapeutic targets while also proposing novel candidates, potentially elucidating alternative pathways for drug action.

**Note 2. Scoring system for target prediction**

**A. Scoring system of different evidence sources.** Open Targets utilizes 23 sources of evidence across six categories: Drug (1), Genetic Association (9), Pathology and Systems Biology (7), Text Mining (1), Somatic Mutation (3), Gene Expression (1), and Animal Model (1). However, the Animal Model database was excluded due to a lack of relevant evidence. Most evidence sources employ different scoring systems based on distinct criteria and statistical methods. We have summarized the database names, evidence types, weights, scoring systems, and background information for each evidence source in Table S8.

**B. Association score for one or multiple types of evidence.** Open Targets calculates a association score for each target (such as a gene or protein) by integrating the scores of evidence in multiple types (see the seven evidence types mentioned above). Each type of evidence contributes to the global score based on a weighted system. For example, evidence from drug and genetic association databases is assigned higher weights (1) due to its experimental nature, while evidence from text mining and computational models receives lower weights (0.2) as it is considered indirect evidence. Here we listed the weights ($W$) for each database: Europe PubMed Central (0.2), Expression Atlas (0.2), IMPC (0.2); PROGENy (0.5), SLAPenrich (0.5), Cancer Biomarkers (0.5), SysBio/gene signature (0.5) and all other database (1). To calculate the Association score ($\hat{S}_i$ for target $i$, the scores from all evidence are first sorted as

$$R_{ij} = Rank(S_{ij}) \tag{1}$$

Where $S_{ij}$ and $R_{ij}$ indicate the score and rank of the $j$ th evidence for target $i$, respectively. Function $Rank()$ returns the rank of each data point in the input vector (greater value gets lower rank). Then a harmonic sum is calculated as

$$\bar{S}_i = \sum_j W_j / R_{ij} \tag{2}$$

Where $\bar{S}_i$ is the raw global score of target $i$ and $W_j$ is the predefined weight for evidence $j$. The final global score ($\hat{S}_i$) is calculated by scaling $\bar{S}$ into the range of 0-1 by dividing $\bar{S}_i$ by the maximum theoretical harmonic sum. By using this approach, stronger and more reliable evidence types have a greater impact on the final score: 0 indicates no evidence and 1 indicates very strong evidence. We also use this approach to calculate a score for each type of evidence (Drug, genetic association, pathology and systemic biology, text mining, somatic mutation and gene expression) which is not provided by OpenTarget.

**Note 3. Data generation and preprossessing**

We compiled and processed six data sources and two additional biological information repositories. The data sources include protein-protein interactions (PPI), gene expression in healthy and diseased cell lines, gene expression in diseased cell lines upon chemical or genetic interventions, phenotype-associated genes, drug targets, and drug indications. The following describes the data sources and preprocessing steps.

**Human protein-protein interaction network.** We built a PPI network by aggregating proteins and protein-protein interactions from BIOGRID[2] (accessed in March 2022), HuRI[3], and Menche et al.[4] In this graph, nodes represent human proteins, and edges exist between nodes if there is physical interaction between proteins. We downloaded a gene ID mapping file from the HUGO Gene Nomenclature Committee. Using this file, we mapped proteins in BIOGRID and Menche et al.[4] from Entrez Gene ID[5] to HUGO Gene Nomenclature Committee ID[6], and proteins in HuRI from Ensembl Gene ID[7] to HUGO Gene Nomenclature Committee ID[6]. Our final PPI comprises the union of nodes and edges, resulting in a graph with 15,742 nodes and 222,498 undirected edges.

**Human protein-protein interaction network for sensitivity analyses.** To test the sensitivity of PDGrapher on protein-protein interactions, we utilized data from STRING (string-db.org), which provides a confidence score for each edge. The specific dataset used was "9606.protein.physical.links .detailed.v12.0.txt.gz" for Homo sapiens. We filtered the edges by using the 0.1, 0.2, 0.3, 0.4, and 0.5 quantiles of the confidence scores as cutoffs, resulting in five PPI networks with 625,818, 582,305, 516,683, 443,051, and 296,451 edges, respectively. Subsequently, we filtered these PPIs to retain only the nodes (proteins) that were present in the gene expression data.

**Gene expression data.** We downloaded the Library of Integrated Network-Based Cellular Signatures (LINCS[8]) level 3 gene expression data from https://clue.io/releases/data-dashboard (accessed in February 2022). Level 3 data consists of quantile-normalized samples across each plate and is appropriate for cross-plate analyses. LINCS contains measurements of gene expression for 12,327 genes after genetic and chemical interventions. There are 387,317 samples upon CRISPR genetic interventions (treated samples), with 5,156 unique knocked-out genes in 27 cell lines. There are an average of 17.18 replicates per cell line-knocked-out gene pair. The number of unique genes knocked out in each cell line varies from 1 to 5,114, with an average of 2,042.14 unique genes knocked out per cell line.

Control data for CRISPR interventions, that is, diseased samples, are genetic interventions that do not contain a gene-specific sequence or whose gene-specific sequence targets a gene not expressed in the human genome. There are 47,781 diseased samples in 50 cell lines. The number of diseased samples for each cell line varies from 1 to 6,890, with an average of 955.62 diseased samples per cell line.

There are 1,313,292 samples after chemical interventions (treated samples), with 31,234 unique compounds in 229 cell lines. There is an average of 7.96 replicates per cell line-compound pair. The number of compounds tested in each cell line varies from 1 to 19,509, with an average of 719.69 unique compounds tested per cell line. Drugs are administered at different doses and measured at varying time points after treatment. On average, there are 2.73 different doses per compound-cell line pair, with a minimum of 1 and a maximum of 26 different doses. On average, gene expression is measured at 1.25 time points per compound-cell line pair, with a minimum of 1 and a maximum of 13 different time points.

Control data for chemical interventions, that is, diseased samples, are treated with a vehicle (dimethyl sulfoxide). There are 76,795 diseased samples across 226 cell lines. The number of diseased samples for each cell line varies from 1 to 7,336, with an average of 339.80 diseased samples per cell line. On average, gene expression of diseased samples is measured at 1.4 time points, with a minimum of 1 and a maximum of 5 different time points.

We restricted our analysis to a subset of LINCS cell lines to ensure sufficient perturbational coverage and the inclusion of healthy cell line counterparts. For chemical perturbagens, we first extracted cell lines with "trt_cp" perturbation types and no recorded "failure_mode", and then filtered out any cell lines with fewer than 1,000 treated samples. Next, we applied the following selection criteria: (1) we selected those that had healthy counterparts available, yielding six cell lines, (2) from the remaining cell lines lacking healthy counterparts, we selected those with more than 15,000 treated samples, (3) of all remaining cell lines, we excluded those which were not found in COSMIC Cell Line Project. Table S9 contains all cell lines that have more than or equal than 1,000 treated samples and their corresponding criteria for exclusion or inclusion. For genetic perturbagens, we first extracted cell lines with "trt_xpr" perturbation types and no recorded "failure_mode", and then filtered out any cell lines with fewer than 5,000 treated samples. Next, we applied the following selection criteria: (1) we selected those with healthy counterparts available, resulting in three cell lines; (2) from the remaining cell lines lacking healthy counterparts, we selected those with more than 15,000 treated samples. Table S10 contains all cell lines that have more or equal than 5,000 treated samples and their corresponding reason for exclusion or inclusion.

To find healthy cell line counterparts, we extracted all cell lines with the "Unknown" tumor phase in the downloaded LINCS dataset (N=145). Then, we filtered the cell lines by tissue type. To find the exact match to diseased cell lines, we performed a manual literature search to confirm their experimental use as healthy counterparts. We extracted healthy counterparts for three of the ten diseased cell lines: cell line NL20 as the healthy counterpart for A549, cell line MCF10A as the healthy counterpart for MCF7, and cell line RWPE1 as the healthy counterpart for PC3.

We acknowledge that NL20 is a normal human bronchial epithelial cell line, while A549 is a human lung carcinoma cell line derived from a tumor in the alveolar region. These cell lines are not perfectly matched. However, without a more closely matched control cell line for lung cancer research, we have opted to use NL20 as a control for A549.

Genetic interventions correspond to gene experiment knockouts in which the gene expression of the knocked-out gene after the intervention is zero. Chemical interventions correspond to small molecule drugs, each targeting one or more proteins. Chemical interventions were performed at different dose levels and measured at different time points. We included replicates measured at all time points and doses. For each cell line and condition (healthy, diseased, and treated), we log-normalized the level 3 gene expression data. We applied a min-max normalization to transform gene expression values into the range $[0, 1]$ following established practices in the field.

We match genes in LINCS to proteins in our PPI using the HUGO Gene Nomenclature Committee ID[6], resulting in 10,716 overlapping genes and 151,839 undirected edges. Furthermore, we excluded treated samples from our datasets whose targeted genes were not included in the PPI.

We have healthy, diseased, and treated gene expression samples for each cell line treated with several genetic or chemical perturbagens (Table S11 and S12). For healthy counterparts, samples with the corresponding treatment ("vector" for genetic perturbagens, and "vehicle" for chemical perturbagens) are not available; therefore, we use the closest possible (see "Sample category" in Table

S11 and S12.

**Gene regulatory networks.** We computed a gene regulatory network (GRN) for each diseased cell line in each condition (genetic and chemical datasets), using the GENIE3[9] algorithm on the gene expression values of each diseased cell line. We filtered genes in our gene expression dataset (LINCS) to contain only those in the PPI before running the GRN algorithm for consistency between the PPI and GRNs. GENIE3, introduced in 2010, won the Dialogue for Reverse Engineering Assessments and Methods 4 (DREAM4) challenge[10], which evaluates the success of GRN inference algorithms on benchmarks of simulated data. GENIE3 was introduced in open source software for bioinformatics bioconductor[11] and is often used for GRN generation[12–15]. It is a model based on an ensemble of regression trees and requires as input a matrix of gene expression levels under various conditions. This expression data are multifactorial. This means that they represent expression levels resulting from a perturbation over a set of genes rather than from a targeted experiment. Multifactorial expression can be obtained as samples from different patients or other biological systems. Therefore, cell line diseased samples are closest to the ideal input data for GENIE3. GENIE3 produces a directed graph that represents regulatory interactions between genes and genes. This is achieved by assigning weights to regulatory links and maximizing weights for more significant links. Then a significance threshold is used to determine which links are substantial enough to be predicted as a regulatory link. We adopted the threshold to generate GRNs with close network density as the PPI from STRING, which was achieved by keeping about 500,000 directed edges.

**Disease-gene information.** We extracted disease-associated genes from COSMIC[16] (Accessed in September 2022) in addition to expert-curated genes available at https://cancer.sanger.ac.uk/cosmic/curation (Accessed in March 2024). Genes were represented using the HUGO Gene Nomenclature Committee ID. For each cell line in our dataset that has disease intervention data (see *Disease intervention data section)*, we extracted cancer-causing mutations as the list of genes with "Verified" *Mutation verification status* in COSMIC that are also included in the list of genes curated by experts. The mapping of the resulting genes to our list of genes in the PPI produced eight disease-associated genes for the lung cancer cell line A549, nine disease-associated genes for breast cancer cell line MCF7, one disease-associated gene for prostate cancer cell line PC3, two disease-associated genes for prostate cancer cell line VCAP, six disease-associated genes for breast cancer cell line MDAMB231, and eight disease-associated genes for breast cancer cell line BT20.

**Drug-target information.** We retrieved drug-target information from DrugBank[17] (accessed in November 2022). We extracted drug names and synonyms, chemical identifiers, drug-gene targets, and all available synonyms for each drug target. Only the nominal targets, genes that produce proteins to which the drug physically interacts, were considered when processing the DrugBank data. Genes or gene products involved in the mechanism of action (MoA) of a drug were excluded and not retained for further analysis. We mapped drugs in DrugBank with chemical perturbagens in LINCS using InChI Key[18], resulting in 1,522 out of 31,234 unique LINCS compounds mapped to DrugBank with information about at least one target. We mapped drug targets to our PPI network using the HUGO Gene Nomenclature Committee ID, excluding any drug target that was not mapped. Chemical interventions target multiple genes, with a minimum of 1, a maximum of 300, and an average of 2.44 targets per compound.

**Cancer drug and target information.** We extracted the list of cancer drugs by cancer type from NCI (https://www.cancer.gov/about-cancer/treatment/types/targeted-therapies/approved-drug-list; Accessed in July 2024). We mapped drug names to DrugBank to obtain cancer drug-gene targets. In total, there are 24 drugs associated with breast cancer (cell lines MCF7 , MDAMB231, and BT20), 30 drugs associated with lung cancer (cell line A549), 11 drugs associated with prostate cancer (cell lines PC3 and VCAP), 13 drugs associated with colon cancer (cell line HT29), 18 drugs associated with skin cancer (cell line A375), three drugs associated with cervical cancer (cell line HELA), five drugs associated with ovarian cancer (cell line ES2), four drugs associated with head and neck cancer (cell line BICR6), five drugs associated with pancreatic cancer (cell line YAPC), five drugs associated with stomach cancer (cell line AGS), and six drugs associated with brain cancer (cell line U251MG).

**Disease intervention data.** Disease intervention datasets consist of gene expression measurements of healthy cell lines, disease-associated genes, and gene expression measurements of diseased cell lines. Gene expression samples from healthy and diseased cell lines were retrieved from LINCS[8], and disease-associated genes were retrieved from COSMIC[16], as detailed previously. Each dataset $T = \{T_1, \ldots, T_M\}$ is a collection of paired healthy-diseased cell lines where in each sample $T =< x^h, U, x^d >$, $x^h$ corresponds to gene expression values of the healthy cell line, set $U$ is comprised by a randomized subset of disease-associated genes, and $x^d$ corresponds to gene expression values of diseased cell lines (that is, upon mutations on genes in $U$). To select the randomized set of disease-associated genes, we first choose a proportion $p \in \{0.25, 0.50, 0.75, 1\}$, and then select $N$ disease-associated genes at random where $N$ is the proportion multiplied by the total number of disease-associated genes. Given that more diseased samples are available than healthy samples (see Table S11 and S12) when building the triplets, we select a random sample from the set of healthy samples and, therefore, have non-unique healthy samples during training. In total, we built three genetic disease intervention datasets and six chemical disease intervention datasets. The first genetic disease intervention dataset is comprised of gene expression of healthy cell line MCF10A, breast cancer mutations, and gene expression of breast cancer cell line MCF7; the second is comprised of gene expression of healthy cell line NL20, lung cancer mutations, and gene expression of lung cancer cell line A549; and the third is comprised of gene expression of healthy cell line RWPE1, prostate cancer mutations, and gene expression of prostate cancer cell line PC3. The first three chemical disease intervention datasets are comprised of gene expression of healthy cell line MCF10A, breast cancer mutations, and gene expression of breast cancer cell line MCF7, MDAMB231, and BT20; the fourth comprised of gene expression of healthy cell line NL20, lung cancer mutations, and gene expression of lung cancer cell line A549; the fifth and sixth comprised of gene expression of healthy cell line RWPE1, prostate cancer mutations, and gene expression of prostate cancer cell line PC3 and VCAP.

**Treatment intervention data - genetic.** Genetic treatment intervention datasets consist of single-gene knockout experiments using CRISPR / Cas9-mediated gene knockout. The genetic treatment intervention data include measurements of gene expression of diseased cell lines, single knocked-out genes, and measurements of gene expression of treated cell lines. Gene expression samples from diseased and treated cell lines and knocked-out genes were retrieved from LINCS[8]. Each dataset $T = \{T_1, \ldots, T_M\}$ is a collection of paired diseased-treated cell lines where in each sample $T =< x^d, U', x^t >$, $x^d$ corresponds to gene expression values of the diseased cell line, set $U'$ is comprised by the knocked-out gene, and $x^t$ corresponds to gene expression values of treated cell lines (that is, upon knocking-out the gene in $U'$). Given that more treated samples are available than diseased samples (see Table S11) when building the triplets, we select a random sample from the set of diseased samples and, therefore, have non-unique diseased samples during training. In total, we built ten datasets of treatment interventions: A549, MCF7, PC3, A375, HT29, ES2, BICR6, YAPC, AGS, and U251MG.

They are comprised of gene expression of diseased cells, knocked-out genes, and gene expression of treated cells. Find more details on data compilation and processing in previous subsections.

**Treatment intervention data - chemical.** Chemical treatment intervention datasets consist of chemical compound treatment experiments. The chemical treatment intervention data include measurements of gene expression of diseased cell lines, drug targets of chemical compounds, and measurements of gene expression of treated cell lines. Gene expression samples of diseased and treated cell lines were retrieved from LINCS, and chemical compound targets were retrieved from DrugBank, as detailed previously. Each dataset $T = \{T_1, \ldots, T_M\}$ is a collection of paired diseased-treated cell lines where in each sample $T =< x^d, U', x^t >$, $x^d$ corresponds to gene expression values of the diseased cell line, set $U'$ is comprised by the chemical compound targets, and $x^t$ correspond to gene expression values of treated cell lines (that is, upon treated with the chemical perturbagen targeting genes in $U'$). Given that more treated samples are available than diseased samples (see Table S12) when building the triplets, we select a random sample from the set of diseased samples and, therefore, have non-unique diseased samples during training. In total, we built nine datasets of treatment interventions: A549, MCF7, PC3, VCAP, MDAMB231, BT20, HT29, A375, and HELA. They are comprised by gene expression of diseased cells, drug targets, and gene expression of treated cells.

#### Note 4. Related works

**Learning optimal interventions.** The problem of learning interventions to achieve a desired state has gained interest in recent years. Recent research formulates this problem as the finding of optimal interventions to optimize an associated outcome [19–22]. These works offer varied approaches. For example, Mueller et al. [19] aim to learn an intervention policy defined by a covariate transformation that produces the largest post-intervention improvement with high uncertainty. Pacchiano et al. [20] formalize the task as a bandit optimization problem in which each bandit's arm corresponds to a covariate to intervene, and the goal is to recover an almost optimal arm in the least number of arm pulls possible. Mueller et al. [21] and Hie et al. [22] approach the problem of sequence-based data where each sequence is associated with an outcome, and the goal is to find mutations in the input sequence that increase a desired outcome. Other recent works formulate this problem as finding optimal interventions to shift the system to a desired state. Zhang et al. [23,24] aimed to find an intervention that applied to a distribution helps match a desired distribution. Specifically, given a distribution $P$ over $X$ and a desired distribution $Q$ over $X$, the goal is to find an optimal matching intervention $I$ such that $P^I$ best matches $Q$ under some metric. They address the special case of soft interventions (shift interventions) and use the expectation of distributions as the distance metric.

**Neural networks and structural causal models (SCMs).** Causal representation learning has been a growing trend in recent years[25]. It aims to combine the strength of traditional causal learning methods with the robust capabilities of deep learning in the face of large and noisy data. Bottlenecks of traditional causal learning methods include unstructured high-dimensional variables, combinatorial optimization problems, unknown intervention, unobserved confounders, selection bias, and estimation bias[25]. There are three areas in which deep learning helps to overcome these bottlenecks [25]. First, in learning causal variables from high-dimensional unstructured data. Second, in learning the causal structure between causal variables, called *causal discovery* within the causal inference literature. Third, it facilitates the inference of interventional and counterfactual queries. Within the last branch, a promising approach aims to join SCMs and neural models to facilitate interventional and counterfactual querying. Parafita et al. put forward the requirements that any DL model should fulfill to approximate causal queries and introduced normalizing causal flows as a specific instantiation[26]. Pawlowski et al. followed a similar approach to introduce a model capable of computing counterfactual queries[27]. Xia et al. approached the problem differently, introducing a Neural Causal Model (NCM), a type of SCM with neural networks as structural equations[28]. Together with the NCM, they introduced an algorithm that provably performs the identification and inference of interventional queries[28]. A follow-up work extended the NCM framework for the identification and inference of counterfactual queries[29]. The concept of NCMs inspires our work by considering the graph in which we operate as a noisy version of a causal graph and our model operating on the graph as a proxy for the structural equations.

**Interventions in graph neural networks (GNNs).** GNNs are a type of neural model that falls under the umbrella term of geometric deep learning[30–32]. These models use graph-structured data to compute transformed representations useful for downstream predictive tasks. Their ability to operate over graphs makes them especially relevant to NCMs. A recent work by Zecevic et al. [33] explored this connection. It introduced interventional GNNs, a GNN in which interventions are represented through mutilations in the input graph, and interventional inference as GNN computations on the mutilated graph[34]. We borrow this concept in our work and extend the representational capabilities of GNNs by assigning learnable embeddings to input nodes.

#### Supplementary Tables

**Table S1:** Number of seen and unseen perturbagens per cell line and across dataset splits, considering chemical perturbations data.

| Cell line | Split | Total | Seen | Unseen |
|---|---|---|---|---|
| **Chemical-PPI/GRN-Lung-A549** | 1 | 858 | 841 | 17 |
| | 2 | 850 | 828 | 22 |
| | 3 | 834 | 821 | 13 |
| | 4 | 853 | 837 | 16 |
| | 5 | 839 | 825 | 14 |
| **Chemical-PPI/GRN-Breast-MCF7** | 1 | 1043 | 1039 | 4 |
| | 2 | 1041 | 1037 | 4 |
| | 3 | 1038 | 1032 | 6 |
| | 4 | 1048 | 1044 | 4 |
| | 5 | 1048 | 1046 | 2 |
| **Chemical-PPI/GRN-Prostate-PC3** | 1 | 1088 | 1086 | 2 |
| | 2 | 1075 | 1073 | 2 |
| | 3 | 1090 | 1086 | 4 |
| | 4 | 1088 | 1085 | 3 |
| | 5 | 1075 | 1071 | 4 |
| **Chemical-PPI/GRN-Prostate-VCAP** | 1 | 574 | 568 | 6 |
| | 2 | 575 | 565 | 10 |
| | 3 | 585 | 579 | 6 |
| | 4 | 567 | 555 | 12 |
| | 5 | 557 | 547 | 10 |
| **Chemical-PPI/GRN-Breast-MDAMB231** | 1 | 458 | 454 | 4 |
| | 2 | 469 | 458 | 11 |
| | 3 | 464 | 459 | 5 |
| | 4 | 459 | 459 | 0 |
| | 5 | 453 | 448 | 5 |
| **Chemical-PPI/GRN-Breast-BT20** | 1 | 37 | 37 | 0 |
| | 2 | 38 | 38 | 0 |
| | 3 | 39 | 37 | 2 |
| | 4 | 37 | 37 | 0 |
| | 5 | 38 | 38 | 0 |
| **Chemical-PPI/GRN-Colon-HT29** | 1 | 884 | 879 | 5 |
| | 2 | 916 | 901 | 15 |
| | 3 | 877 | 860 | 17 |
| | 4 | 887 | 883 | 4 |
| | 5 | 877 | 867 | 10 |
| **Chemical-PPI/GRN-Skin-A375** | 1 | 924 | 910 | 14 |
| | 2 | 933 | 917 | 16 |
| | 3 | 911 | 901 | 10 |
| | 4 | 948 | 927 | 21 |
| | 5 | 939 | 919 | 20 |
| **Chemical-PPI/GRN-Cervix-HELA** | 1 | 659 | 658 | 1 |
| | 2 | 657 | 657 | 0 |
| | 3 | 669 | 668 | 1 |
| | 4 | 649 | 649 | 0 |
| | 5 | 664 | 664 | 0 |

**Table S2:** Number of unseen perturbagens in each dataset split across cell lines for chemical perturbation data.

| Cell line | Split | A375 | A549 | BT20 | HA1E | HELA | HT29 | MCF7 | MDAMB231 | PC3 | VCAP |
|---|---|---|---|---|---|---|---|---|---|---|---|
| Chemical-PPI/GRN-Skin-A375 | 1 | 14 | 44 | 0 | 32 | 3 | 26 | 118 | 3 | 113 | 22 |
| | 2 | 16 | 45 | 0 | 33 | 4 | 26 | 118 | 4 | 114 | 21 |
| | 3 | 10 | 41 | 0 | 28 | 2 | 22 | 114 | 2 | 109 | 20 |
| | 4 | 21 | 51 | 0 | 39 | 6 | 32 | 124 | 6 | 119 | 27 |
| | 5 | 20 | 48 | 0 | 36 | 5 | 29 | 121 | 5 | 116 | 24 |
| Chemical-PPI/GRN-Lung-A549 | 1 | 95 | 17 | 0 | 76 | 37 | 64 | 146 | 1 | 173 | 36 |
| | 2 | 100 | 22 | 0 | 81 | 36 | 69 | 151 | 0 | 178 | 41 |
| | 3 | 94 | 13 | 0 | 74 | 35 | 63 | 145 | 0 | 172 | 36 |
| | 4 | 93 | 16 | 0 | 74 | 38 | 62 | 144 | 0 | 171 | 34 |
| | 5 | 92 | 14 | 0 | 73 | 34 | 61 | 143 | 0 | 170 | 35 |
| Chemical-PPI/GRN-Breast-BT20 | 1 | 1055 | 1003 | 0 | 1030 | 653 | 1015 | 1116 | 488 | 1144 | 712 |
| | 2 | 1054 | 1002 | 0 | 1029 | 652 | 1014 | 1115 | 487 | 1143 | 711 |
| | 3 | 1056 | 1004 | 2 | 1031 | 654 | 1016 | 1117 | 489 | 1145 | 713 |
| | 4 | 1055 | 1003 | 0 | 1030 | 653 | 1015 | 1116 | 488 | 1144 | 712 |
| | 5 | 1054 | 1002 | 0 | 1029 | 652 | 1014 | 1115 | 487 | 1143 | 711 |
| Chemical-PPI/GRN-Cervix-HELA | 1 | 413 | 392 | 8 | 386 | 1 | 385 | 486 | 10 | 500 | 357 |
| | 2 | 412 | 391 | 8 | 385 | 0 | 384 | 485 | 9 | 499 | 357 |
| | 3 | 413 | 392 | 8 | 386 | 1 | 385 | 486 | 10 | 500 | 358 |
| | 4 | 413 | 392 | 8 | 386 | 0 | 385 | 486 | 10 | 500 | 358 |
| | 5 | 412 | 391 | 8 | 385 | 0 | 384 | 485 | 9 | 499 | 357 |
| Chemical-PPI/GRN-Colon-HT29 | 1 | 53 | 41 | 0 | 32 | 14 | 5 | 117 | 0 | 139 | 9 |
| | 2 | 59 | 46 | 0 | 38 | 15 | 15 | 123 | 1 | 145 | 15 |
| | 3 | 64 | 52 | 0 | 43 | 16 | 17 | 128 | 0 | 150 | 21 |
| | 4 | 52 | 40 | 0 | 30 | 14 | 4 | 116 | 0 | 138 | 9 |
| | 5 | 58 | 46 | 0 | 37 | 14 | 10 | 122 | 0 | 144 | 15 |
| Chemical-PPI/GRN-Breast-MCF7 | 1 | 36 | 14 | 0 | 15 | 14 | 8 | 4 | 0 | 40 | 2 |
| | 2 | 38 | 16 | 0 | 17 | 14 | 10 | 4 | 0 | 40 | 4 |
| | 3 | 39 | 17 | 0 | 18 | 15 | 11 | 6 | 0 | 42 | 4 |
| | 4 | 38 | 16 | 0 | 17 | 14 | 10 | 4 | 0 | 40 | 4 |
| | 5 | 37 | 14 | 0 | 16 | 15 | 8 | 2 | 0 | 40 | 2 |
| Chemical-PPI/GRN-Breast-MDAMB231 | 1 | 575 | 521 | 0 | 548 | 172 | 533 | 634 | 4 | 662 | 459 |
| | 2 | 583 | 529 | 0 | 556 | 180 | 541 | 642 | 11 | 670 | 463 |
| | 3 | 574 | 520 | 0 | 547 | 171 | 532 | 633 | 5 | 661 | 456 |
| | 4 | 571 | 517 | 0 | 544 | 168 | 529 | 630 | 0 | 658 | 456 |
| | 5 | 577 | 523 | 0 | 550 | 174 | 535 | 636 | 5 | 664 | 460 |
| Chemical-PPI/GRN-Prostate-PC3 | 1 | 3 | 13 | 0 | 3 | 0 | 2 | 11 | 0 | 2 | 2 |
| | 2 | 3 | 13 | 0 | 3 | 1 | 2 | 11 | 1 | 2 | 1 |
| | 3 | 3 | 13 | 0 | 3 | 1 | 2 | 14 | 1 | 4 | 1 |
| | 4 | 3 | 13 | 0 | 3 | 1 | 2 | 14 | 1 | 3 | 1 |
| | 5 | 3 | 13 | 0 | 3 | 1 | 2 | 13 | 1 | 4 | 1 |
| Chemical-PPI/GRN-Prostate-VCAP | 1 | 363 | 327 | 13 | 341 | 305 | 322 | 423 | 245 | 451 | 6 |
| | 2 | 369 | 333 | 13 | 347 | 306 | 328 | 429 | 246 | 457 | 10 |
| | 3 | 365 | 329 | 12 | 343 | 305 | 324 | 425 | 244 | 453 | 6 |
| | 4 | 370 | 334 | 15 | 348 | 307 | 329 | 430 | 247 | 458 | 12 |
| | 5 | 371 | 335 | 13 | 349 | 307 | 330 | 431 | 246 | 459 | 10 |

**Table S3:** Number of seen and unseen perturbagens per cell line per dataset split with genetic perturbation data.

| Cell line | Split | Total | Seen | Unseen |
|---|---|---|---|---|
| **Genetic-PPI-Lung-A549** | 1 | 2697 | 2666 | 31 |
| | 2 | 2672 | 2648 | 24 |
| | 3 | 2681 | 2656 | 25 |
| | 4 | 2654 | 2626 | 28 |
| | 5 | 2704 | 2676 | 28 |
| **Genetic-PPI-Breast-MCF7** | 1 | 2230 | 2216 | 14 |
| | 2 | 2266 | 2252 | 14 |
| | 3 | 2255 | 2247 | 8 |
| | 4 | 2253 | 2246 | 7 |
| | 5 | 2261 | 2250 | 11 |
| **Genetic-PPI-Prostate-PC3** | 1 | 2588 | 2559 | 29 |
| | 2 | 2597 | 2570 | 27 |
| | 3 | 2635 | 2609 | 26 |
| | 4 | 2603 | 2579 | 24 |
| | 5 | 2605 | 2584 | 21 |
| **Genetic-PPI-Skin-A375** | 1 | 2552 | 2507 | 45 |
| | 2 | 2524 | 2489 | 35 |
| | 3 | 2518 | 2472 | 46 |
| | 4 | 2565 | 2504 | 61 |
| | 5 | 2538 | 2497 | 41 |
| **Genetic-PPI-Colon-HT29** | 1 | 2539 | 2495 | 44 |
| | 2 | 2524 | 2477 | 47 |
| | 3 | 2544 | 2506 | 38 |
| | 4 | 2564 | 2529 | 35 |
| | 5 | 2563 | 2523 | 40 |
| **Genetic-PPI-Ovary-ES2** | 1 | 2579 | 2522 | 57 |
| | 2 | 2557 | 2505 | 52 |
| | 3 | 2591 | 2526 | 65 |
| | 4 | 2617 | 2553 | 64 |
| | 5 | 2573 | 2520 | 53 |
| **Genetic-PPI-Head-BICR6** | 1 | 2597 | 2568 | 29 |
| | 2 | 2601 | 2573 | 28 |
| | 3 | 2592 | 2565 | 27 |
| | 4 | 2602 | 2564 | 38 |
| | 5 | 2573 | 2549 | 24 |
| **Genetic-PPI-Pancreas-YAPC** | 1 | 2502 | 2458 | 44 |
| | 2 | 2524 | 2480 | 44 |
| | 3 | 2504 | 2461 | 43 |
| | 4 | 2491 | 2453 | 38 |
| | 5 | 2539 | 2504 | 35 |
| **Genetic-PPI-Stomach-AGS** | 1 | 2574 | 2544 | 30 |
| | 2 | 2637 | 2606 | 31 |
| | 3 | 2615 | 2586 | 29 |
| | 4 | 2573 | 2542 | 31 |
| | 5 | 2590 | 2565 | 25 |
| **Genetic-PPI-Brain-U251MG** | 1 | 2784 | 2772 | 12 |
| | 2 | 2782 | 2774 | 8 |
| | 3 | 2834 | 2827 | 7 |
| | 4 | 2810 | 2796 | 14 |
| | 5 | 2792 | 2778 | 14 |

**Table S4:** Number of unseen perturbagens in each dataset split across cell lines for genetic perturbation data.

| Cell Line | Split | A375 | A549 | AGS | BICR6 | ES2 | HT29 | MCF7 | PC3 | U251MG | YAPC |
|---|---|---|---|---|---|---|---|---|---|---|---|
| **Genetic-PPI-Skin-A375** | 1 | 45 | 59 | 59 | 59 | 64 | 58 | 53 | 59 | 59 | 59 |
| | 2 | 35 | 54 | 54 | 54 | 59 | 54 | 53 | 54 | 54 | 54 |
| | 3 | 46 | 66 | 66 | 65 | 71 | 66 | 59 | 66 | 66 | 66 |
| | 4 | 61 | 80 | 81 | 81 | 85 | 81 | 76 | 81 | 81 | 81 |
| | 5 | 41 | 62 | 62 | 62 | 66 | 62 | 56 | 62 | 62 | 62 |
| **Genetic-PPI-Lung-A549** | 1 | 40 | 31 | 35 | 35 | 43 | 35 | 31 | 35 | 35 | 35 |
| | 2 | 33 | 24 | 29 | 29 | 37 | 29 | 23 | 29 | 29 | 29 |
| | 3 | 35 | 25 | 30 | 30 | 40 | 30 | 26 | 30 | 30 | 30 |
| | 4 | 39 | 28 | 34 | 34 | 43 | 34 | 31 | 34 | 34 | 34 |
| | 5 | 42 | 28 | 37 | 37 | 45 | 37 | 33 | 36 | 37 | 37 |
| **Genetic-PPI-Stomach-AGS** | 1 | 37 | 32 | 30 | 32 | 40 | 32 | 21 | 32 | 32 | 32 |
| | 2 | 38 | 32 | 31 | 33 | 42 | 33 | 21 | 33 | 33 | 33 |
| | 3 | 36 | 31 | 29 | 31 | 41 | 31 | 25 | 31 | 31 | 31 |
| | 4 | 38 | 33 | 31 | 33 | 41 | 33 | 26 | 33 | 33 | 33 |
| | 5 | 30 | 26 | 25 | 25 | 34 | 26 | 20 | 26 | 26 | 26 |
| **Genetic-PPI-Head-BICR6** | 1 | 41 | 36 | 36 | 29 | 43 | 36 | 27 | 36 | 36 | 36 |
| | 2 | 39 | 34 | 34 | 28 | 43 | 33 | 31 | 34 | 34 | 34 |
| | 3 | 36 | 31 | 31 | 27 | 40 | 31 | 28 | 31 | 31 | 31 |
| | 4 | 48 | 44 | 44 | 38 | 52 | 44 | 39 | 44 | 44 | 44 |
| | 5 | 39 | 35 | 35 | 24 | 43 | 35 | 30 | 34 | 35 | 34 |
| **Genetic-PPI-Ovary-ES2** | 1 | 153 | 151 | 151 | 150 | 57 | 149 | 149 | 150 | 151 | 151 |
| | 2 | 152 | 146 | 147 | 147 | 52 | 147 | 143 | 146 | 147 | 147 |
| | 3 | 151 | 151 | 151 | 151 | 65 | 151 | 150 | 150 | 151 | 151 |
| | 4 | 144 | 141 | 141 | 141 | 64 | 141 | 140 | 140 | 141 | 141 |
| | 5 | 148 | 146 | 146 | 146 | 53 | 146 | 144 | 145 | 146 | 146 |
| **Genetic-PPI-Colon-HT29** | 1 | 60 | 55 | 55 | 55 | 62 | 44 | 46 | 55 | 55 | 55 |
| | 2 | 66 | 61 | 61 | 61 | 64 | 47 | 53 | 61 | 61 | 61 |
| | 3 | 58 | 53 | 53 | 53 | 62 | 38 | 48 | 53 | 53 | 53 |
| | 4 | 52 | 48 | 48 | 48 | 53 | 35 | 43 | 48 | 48 | 48 |
| | 5 | 64 | 60 | 60 | 60 | 65 | 40 | 49 | 60 | 60 | 60 |
| **Genetic-PPI-Breast-MCF7** | 1 | 642 | 637 | 637 | 637 | 647 | 637 | 14 | 637 | 637 | 637 |
| | 2 | 641 | 636 | 636 | 636 | 646 | 636 | 14 | 636 | 636 | 636 |
| | 3 | 634 | 630 | 630 | 630 | 640 | 630 | 8 | 630 | 630 | 630 |
| | 4 | 634 | 629 | 629 | 629 | 639 | 629 | 7 | 629 | 629 | 629 |
| | 5 | 638 | 633 | 633 | 633 | 643 | 633 | 11 | 633 | 633 | 633 |
| **Genetic-PPI-Prostate-PC3** | 1 | 37 | 32 | 32 | 32 | 37 | 32 | 30 | 29 | 32 | 32 |
| | 2 | 36 | 30 | 31 | 31 | 37 | 31 | 26 | 27 | 31 | 31 |
| | 3 | 37 | 32 | 32 | 32 | 37 | 32 | 30 | 26 | 32 | 32 |
| | 4 | 32 | 27 | 27 | 27 | 35 | 26 | 22 | 24 | 27 | 27 |
| | 5 | 31 | 26 | 26 | 26 | 34 | 26 | 25 | 21 | 26 | 26 |
| **Genetic-PPI-Brain-U251MG** | 1 | 17 | 12 | 12 | 12 | 22 | 12 | 11 | 12 | 12 | 12 |
| | 2 | 13 | 8 | 8 | 8 | 18 | 8 | 8 | 8 | 8 | 7 |
| | 3 | 12 | 7 | 7 | 7 | 17 | 7 | 7 | 7 | 7 | 7 |
| | 4 | 19 | 14 | 14 | 14 | 24 | 14 | 13 | 14 | 14 | 14 |
| | 5 | 19 | 14 | 14 | 14 | 24 | 14 | 13 | 14 | 14 | 14 |
| **Genetic-PPI-Pancreas-YAPC** | 1 | 56 | 51 | 51 | 51 | 58 | 51 | 45 | 51 | 51 | 44 |
| | 2 | 53 | 48 | 48 | 48 | 56 | 48 | 38 | 48 | 48 | 44 |
| | 3 | 56 | 51 | 51 | 51 | 58 | 51 | 44 | 51 | 51 | 43 |
| | 4 | 50 | 45 | 45 | 45 | 54 | 45 | 37 | 45 | 45 | 38 |
| | 5 | 52 | 47 | 47 | 47 | 54 | 47 | 38 | 47 | 47 | 35 |

**Table S5:** Table containing number of edges remaining, number of edges removed, fraction of edges removed (fraction of the total number of edges in the PPI), and number of connected components ordered by increasing fraction of bridge edges removed in synthetic experiments (Extended Data Figure 5).

| Fraction | Remaining edges | Removed edges | Removed edges (fraction) | Connected components |
|---|---|---|---|---|
| 0.1 | 303,500 | 178 | 0.0005 | 90 |
| 0.2 | 303,322 | 356 | 0.0011 | 179 |
| 0.3 | 303,144 | 534 | 0.0017 | 268 |
| 0.4 | 302,964 | 714 | 0.0023 | 358 |
| 0.5 | 302,786 | 892 | 0.0029 | 447 |
| 0.6 | 302,608 | 1,070 | 0.0035 | 536 |
| 0.7 | 302,428 | 1,250 | 0.0041 | 626 |
| 0.8 | 302,250 | 1,428 | 0.0047 | 715 |
| 0.9 | 302,072 | 1,606 | 0.0052 | 804 |
| 1.0 | 301,892 | 1,786 | 0.0058 | 894 |

**Table S6: PDGrapher predicts targets that are closer in the network to ground-truth targets than what would be expected by change in Chemical PPI datasets in splits with new samples.** Table with median distance between ground-truth targets and targets predicted by PDGrapher and a Random baseline, together with U-statistic and p-value of one-sided Mann-Whitney U test, and effect size (rank-biserial correlation) with 95% confidence interval.

| **Dataset** | **PDGrapher** | **Random** | **Effect size $\pm$ CI** | **U** | **p-value** |
|---|---|---|---|---|---|
| Colon-HT29 | 3.0 | 3.0 | $0.1645 \pm [0.1626, 0.1664]$ | 1.3032e11 | $< 0.001$ |
| Prostate-PC3 | 3.0 | 3.0 | $0.1142 \pm [0.1127, 0.1156]$ | 4.1938e11 | $< 0.001$ |
| Cervix-HELA | 3.0 | 3.0 | $0.2113 \pm [0.2094, 0.2130]$ | 1.5735e11 | $< 0.001$ |
| Breast-MCF7 | 3.0 | 3.0 | $0.2160 \pm [0.2146, 0.2174]$ | 3.9197e11 | $< 0.001$ |
| Lung-A549 | 3.0 | 3.0 | $0.3531 \pm [0.3515, 0.3549]$ | 1.2900e11 | $< 0.001$ |
| Skin-A375 | 3.0 | 3.0 | $0.2648 \pm [0.2633, 0.2662]$ | 2.2462e11 | $< 0.001$ |
| Prostate-VCAP | 3.0 | 3.0 | $0.2083 \pm [0.2053, 0.2111]$ | 2.1493e10 | $< 0.001$ |
| Breast-MDAMB231 | 3.0 | 3.0 | $0.3038 \pm [0.3012, 0.3062]$ | 2.6933e10 | $< 0.001$ |
| Breast-BT20 | 2.0 | 3.0 | $0.5748 \pm [0.5699, 0.5799]$ | 5.7782e8 | $< 0.001$ |

**Table S7: PDGrapher predicts targets that are closer in the network to ground-truth targets than what would be expected by change in Chemical PPI datasets in splits with new samples and cell lines.** Table with median distance between ground-truth targets and targets predicted by PDGrapher and a Random baseline, together with U-statistic and p-value of one-sided Mann-Whitney U test, and effect size (rank-biserial correlation) with 95% confidence interval.

| **Dataset** | **PDGrapher** | **Random** | **Effect size ± CI** | **U** | **p-value** |
|---|---|---|---|---|---|
| Breast-BT20 | 2.0 | 3.0 | 0.2296 $\pm[0.2264, 0.2326]$ | 1.5779e10 | $< 0.001$ |
| Breast-MDAMB231 | 3.0 | 3.0 | 0.2423 $\pm[0.2409, 0.2436]$ | 4.6308e11 | $< 0.001$ |
| Lung-A549 | 3.0 | 3.0 | 0.2191 $\pm[0.2182, 0.2200]$ | 2.4676e12 | $< 0.001$ |
| Cervix-HELA | 3.0 | 3.0 | 0.2490 $\pm[0.2481, 0.2500]$ | 2.3807e12 | $< 0.001$ |
| Breast-MCF7 | 3.0 | 3.0 | 0.2457 $\pm[0.2451, 0.2464]$ | 6.0754e12 | $< 0.001$ |
| Skin-A375 | 3.0 | 3.0 | 0.2384 $\pm[0.2377, 0.2392]$ | 3.6503e12 | $< 0.001$ |
| Colon-HT29 | 3.0 | 3.0 | 0.2571 $\pm[0.2562, 0.2581]$ | 1.8346e12 | $< 0.001$ |
| Prostate-PC3 | 3.0 | 3.0 | 0.2569 $\pm[0.2562, 0.2576]$ | 5.6374e12 | $< 0.001$ |
| Prostate-VCAP | 3.0 | 3.0 | 0.2486 $\pm[0.2472, 0.2501]$ | 3.1982e11 | $< 0.001$ |

**Table S8: Scoring system of evidence for targets (Due to extended size of this table, we provide it as a separate XLSX file.)**

**Table S9: Cell line selection in chemical LINCS dataset.** For chemical perturbagens, we first extracted cell lines with `trt_cp` perturbation types and no recorded `failure_mode`, and then filtered out any cell lines with fewer than 1,000 treated samples. Next, we applied the following selection criteria: (1) we selected those that had healthy counterparts available, yielding six cell lines, (2) from the remaining cell lines lacking healthy counterparts, we selected those with more than 15,000 treated samples, (3) of all remaining cell lines, we excluded those which were not found in COSMIC Cell Line Project. This table contains all cell lines that have more or equal than 1,000 treated samples and their corresponding reason for exclusion or inclusion.

| Cell line | Number of treated samples | Healthy counterpart in data | Inclusion label | Reason for inclusion/exclusion |
|---|---|---|---|---|
| MCF7 | 35935 | MCF10A | Included | Meets selection criteria (1) |
| PC3 | 33173 | RWPE1 | Included | Meets selection criteria (1) |
| A375 | 25946 | - | Included | Meets selection criteria (2) |
| A549 | 23423 | NL20 | Included | Meets selection criteria (1) |
| HELA | 20894 | - | Included | Meets selection criteria (2) |
| HT29 | 19724 | - | Included | Meets selection criteria (2) |
| YAPC | 12598 | - | Excluded | Does not meet selection criteria (2) |
| MDAMB231 | 10167 | MCF10A | Included | Meets selection criteria (1) |
| THP1 | 8642 | - | Excluded | Does not meet selection criteria (2) |
| U2OS | 7688 | - | Excluded | Does not meet selection criteria (2) |
| VCAP | 7469 | RWPE1 | Included | Meets selection criteria (1) |
| NPC | 7337 | - | Excluded | Does not meet selection criteria (2) |
| HCC515 | 7176 | - | Excluded | Does not meet selection criteria (2) |
| JURKAT | 6334 | - | Excluded | Does not meet selection criteria (2) |
| HEPG2 | 6138 | - | Excluded | Does not meet selection criteria (2) |
| XC.L10 | 3993 | - | Excluded | Does not meet selection criteria (2) |
| HAP1 | 1791 | - | Excluded | Does not meet selection criteria (2) |
| SKBR3 | 1714 | MCF10A | Excluded | Does not meet selection criteria (3) |
| BT20 | 1403 | MCF10A | Included | Meets selection criteria (1) |
| SKL | 1351 | - | Excluded | Does not meet selection criteria (2) |
| XC.L100 | 1218 | - | Excluded | Does not meet selection criteria (2) |
| TMDB | 1184 | - | Excluded | Does not meet selection criteria (2) |
| XC.R10 | 1166 | - | Excluded | Does not meet selection criteria (2) |
| HCT116 | 1143 | - | Excluded | Does not meet selection criteria (2) |
| HS578T | 1049 | - | Excluded | Does not meet selection criteria (2) |
| OCILY19 | 1047 | - | Excluded | Does not meet selection criteria (2) |

**Table S10: Cell line selection in genetic LINCS dataset.** For genetic perturbagens, we first extracted cell lines with `trt_xpr` perturbation types and no recorded `failure_mode`, and then filtered out any cell lines with fewer than 5,000 treated samples. Next, we applied the following selection criteria: (1) we selected those that had healthy counterparts available, yielding three cell lines, (2) from the remaining cell lines lacking healthy counterparts, we selected those with more than 15,000 treated samples. This table contains all cell lines that have more or equal than 5,000 treated samples and their corresponding reason for exclusion or inclusion.

| **Cell line** | **Number of treated samples** | **Healthy counterpart in data** | **Inclusion label** | **Reason for inclusion/exclusion** |
|---|---|---|---|---|
| U251MG | 28337 | - | Included | Meets selection criteria (2) |
| A549 | 26094 | NL20 | Included | Meets selection criteria (1) |
| ES2 | 25540 | - | Included | Meets selection criteria (2) |
| A375 | 23523 | - | Included | Meets selection criteria (2) |
| AGS | 23071 | - | Included | Meets selection criteria (2) |
| PC3 | 23040 | RWPE1 | Included | Meets selection criteria (1) |
| BICR6 | 22981 | - | Included | Meets selection criteria (2) |
| HT29 | 22220 | - | Included | Meets selection criteria (2) |
| YAPC | 21809 | - | Included | Meets selection criteria (2) |
| MCF7 | 20328 | MCF10A | Included | Meets selection criteria (1) |

**Table S11:** Table shows several healthy, diseased, and treated samples for lung cancer (A549), breast cancer (MCF7), and prostate cancer (PC3), and only diseased and treated samples for skin cancer (A375), colon cancer (HT29), ovarian cancer (ES2), head and neck cancer (BICR6), pancreatic cancer (YAPC), stomach cancer (AGS), and brain cancer (U251MG) with genetic perturbations.

| Cancer type | Cell line | Sample type | N samples | Category | N perturbagens |
|---|---|---|---|---|---|
| Lung cancer | A549 | healthy | 50 | vehicle | - |
| | | diseased | 4,327 | vector | - |
| | | treated | 24,255 | CRISPR | 3,711 |
| Breast cancer | MCF7 | healthy | 113 | untreated | - |
| | | diseased | 4,852 | vector | - |
| | | treated | 18,774 | CRISPR | 3,090 |
| Prostate cancer | PC3 | healthy | 185 | vector | - |
| | | diseased | 6,890 | vector | - |
| | | treated | 21,229 | CRISPR | 3,710 |
| Skin cancer | A375 | diseased | 4,777 | vector | - |
| | | treated | 21,794 | CRISPR | 3,709 |
| Colon cancer | HT29 | diseased | 4,235 | vector | - |
| | | treated | 20,525 | CRISPR | 3,706 |
| Ovary cancer | ES2 | diseased | 1,277 | vector | - |
| | | treated | 23,708 | CRISPR | 3,654 |
| Head and Neck | BICR6 | diseased | 1,362 | vector | - |
| | | treated | 21,183 | CRISPR | 3,711 |
| Pancreas cancer | YAPC | diseased | 1,275 | vector | - |
| | | treated | 20,135 | CRISPR | 3,711 |
| Stomach cancer | AGS | diseased | 1,352 | vector | - |
| | | treated | 21,284 | CRISPR | 3,712 |
| Brain cancer | U251MG | diseased | 1,449 | vector | - |
| | | treated | 26,323 | CRISPR | 3,712 |

**Table S12:** Table shows several healthy, diseased, and treated samples for lung cancer (A549), breast cancer (MCF7, MDAMB231, and BT20), and prostate cancer (PC3 and VCAP), and only diseased and treated samples for cervical cancer (HELA), colon cancer (HT29), and skin cancer (A375) with chemical perturbations.

| Cancer type | Cell line | Sample type | N samples | Category | N perturbagens |
|---|---|---|---|---|---|
| Lung cancer | A549 | healthy | 50 | vehicle | - |
| | | diseased | 5,261 | vehicle | - |
| | | treated | 23,100 | compound | 1,041 |
| Breast cancer | MCF7 | healthy | 2,675 | untreated | - |
| | | diseased | 7,336 | vehicle | - |
| | | treated | 35,421 | compound | 1,154 |
| Breast cancer | MDAMB231 | healthy | 2,675 | untreated | - |
| | | diseased | 1,591 | vehicle | - |
| | | treated | 10,004 | compound | 526 |
| Breast cancer | BT20 | healthy | 2,675 | untreated | - |
| | | diseased | 409 | vehicle | - |
| | | treated | 1,403 | compound | 39 |
| Prostate cancer | PC3 | healthy | 185 | vector | - |
| | | diseased | 7,202 | vehicle | - |
| | | treated | 32,555 | compound | 1,182 |
| Prostate cancer | VCAP | healthy | 185 | untreated | - |
| | | diseased | 3,904 | vehicle | - |
| | | treated | 7,364 | compound | 738 |
| Colon cancer | HT29 | diseased | 4,317 | vehicle | - |
| | | treated | 19,386 | compound | 1,053 |
| Skin cancer | A375 | diseased | 5,165 | vehicle | - |
| | | treated | 25347 | compound | 1,093 |
| Cervix cancer | HELA | diseased | 2,905 | vehicle | - |
| | | treated | 20,308 | compound | 683 |

**Table S13:** Problem formulation of PDGrapher

| Task | Number of graphs | Number of node attribute sets | Label dimensions |
|---|---|---|---|
| Graph Classification | m | m | m x 1 (one for each graph) |
| Node Classification | 1 | 1 | 1 x n (one for each node) |
| Ours | 1 | m | m x n (one for each node of each graph) |

### Supplementary Figures

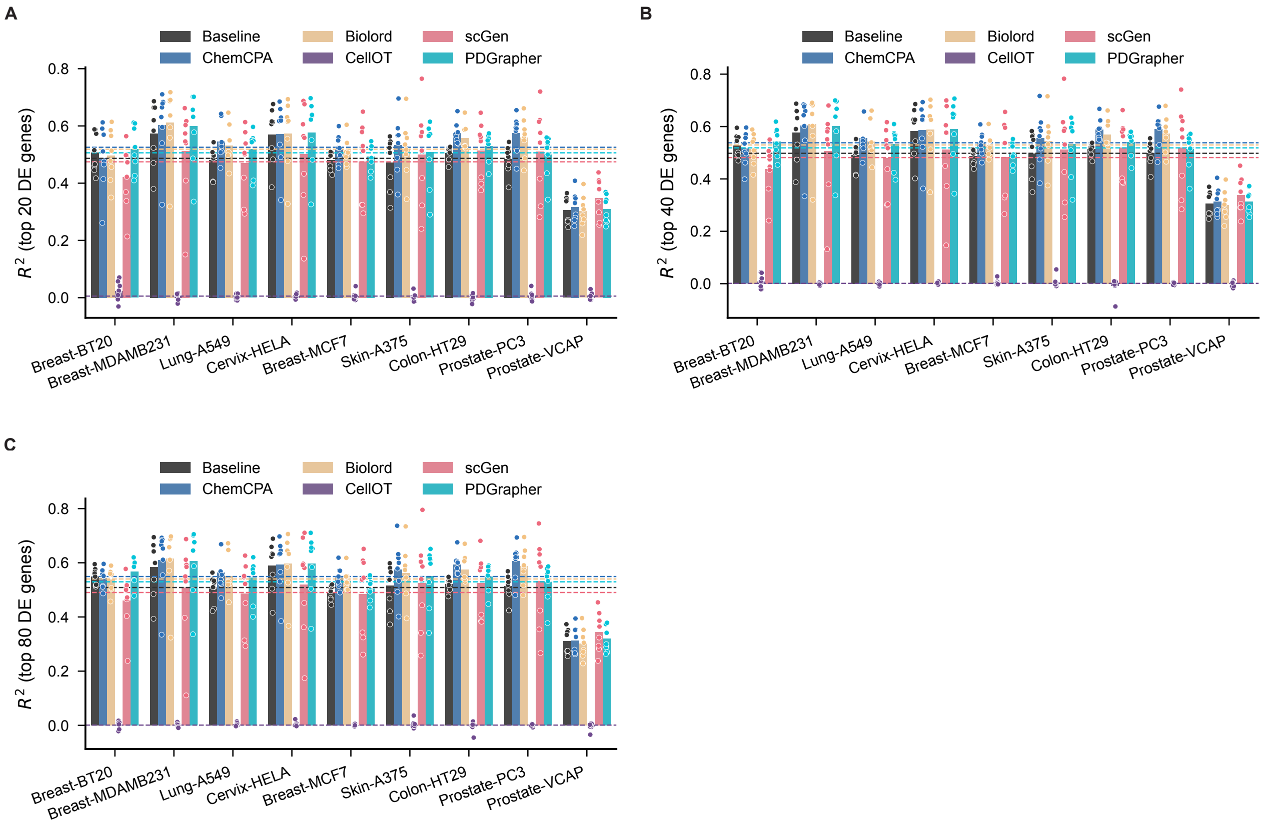


**Figure S1: The performance of response prediction across nine cell lines under chemical perturbation**. We trained each model on a single cell line and made predictions on the remaining eight cell lines. The $R^2$ values are calculated between the predicted and actual gene expression for the top 20 (**A**), 40 (**B**), and 80 (**C**) differentially expressed genes per cell line. Dotted lines represent the average performance across cell lines, dots indicate individual data points, and bars represent the average $R^2$ across eight cell lines. Statistical tests were conducted between PDGrapher and each competing method for each cell line (n = 8 for scGen and CellOT and n = 40 for other methods), and the corresponding P-values are provided in the Source Data.

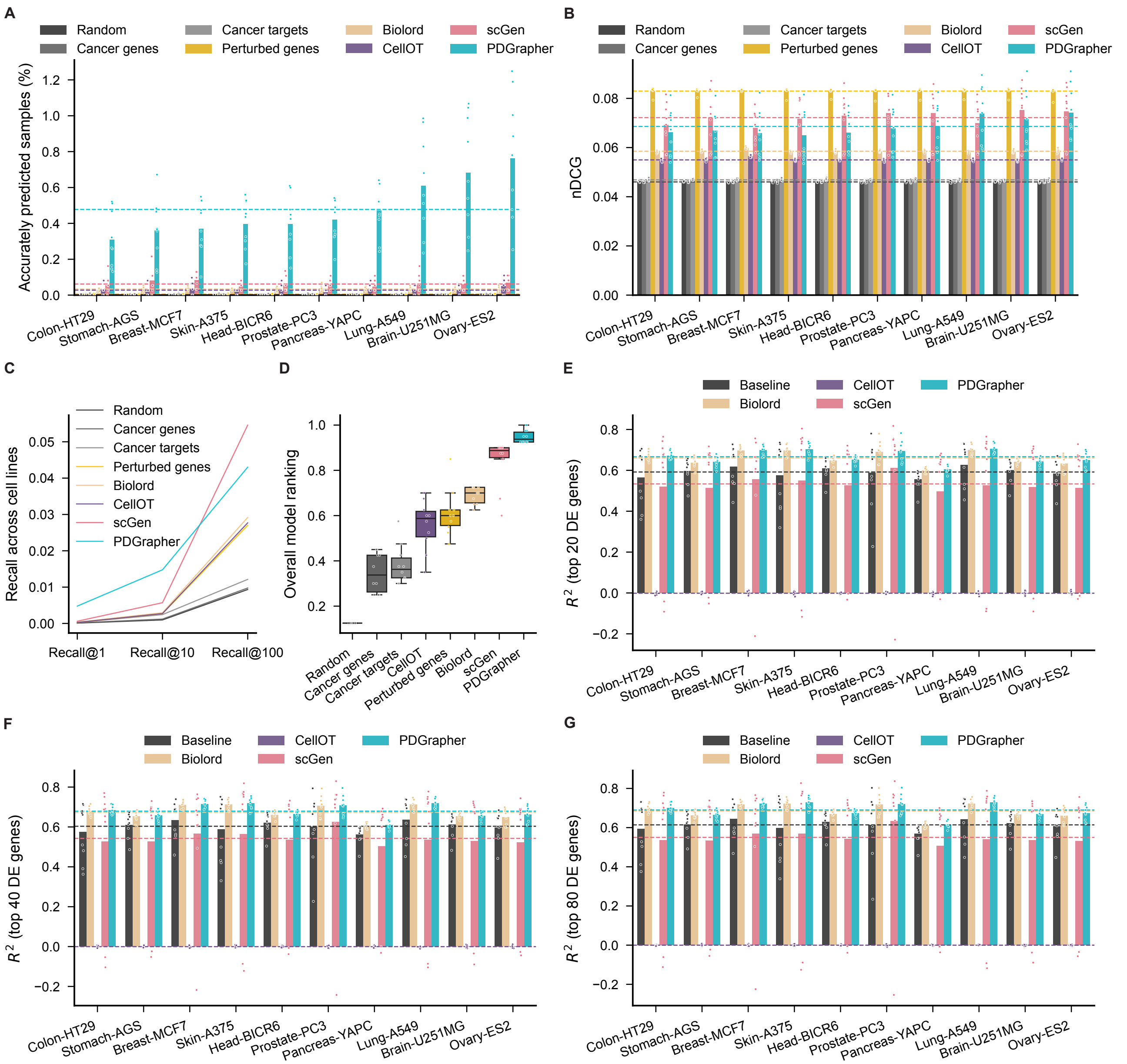


**Figure S2: PDGrapher predicts genetic perturbagens to shift cells from diseased to treated states in unseen cell lines**. **(A)** PDGrapher provides accurate predictions for up to 0.68% (Genetic-PPI-Ovary-ES2: 0.75% vs 0.07%) more samples in the test set compared to the second-best baseline across Genetic-PPI datasets **(B)** The Perturbed genes baseline takes the leading position in nDCG across genetic Genetic-PPI datasets. **(C)** PDGrapher recovers ground-truth therapeutic targets at comparable rates compared to competing methods for Genetic-PPI datasets. **(D)** PDGrapher has the second-best overall performance in perturbagen prediction in unseen cell lines evaluated by the averaged rank over multiple cell lines and metrics. **(E-G)** Shown is the $R^2$ of the response prediction module of PDGrapher compared to competing baselines for the top 20 (E), 40 (F), and 80 (G) differentially expressed (DE) genes. The central line inside the box represents the median, while the top and bottom edges correspond to the first (Q1) and third (Q3) quartiles. The whiskers extend to the smallest and largest values within 1.5 times the interquartile range (IQR) from the quartiles. Statistical tests were conducted between PDGrapher and each competing method for each cell line (n = 9 for scGen and CellOT and n = 45 for other methods), and the corresponding P-values are provided in the Source Data.

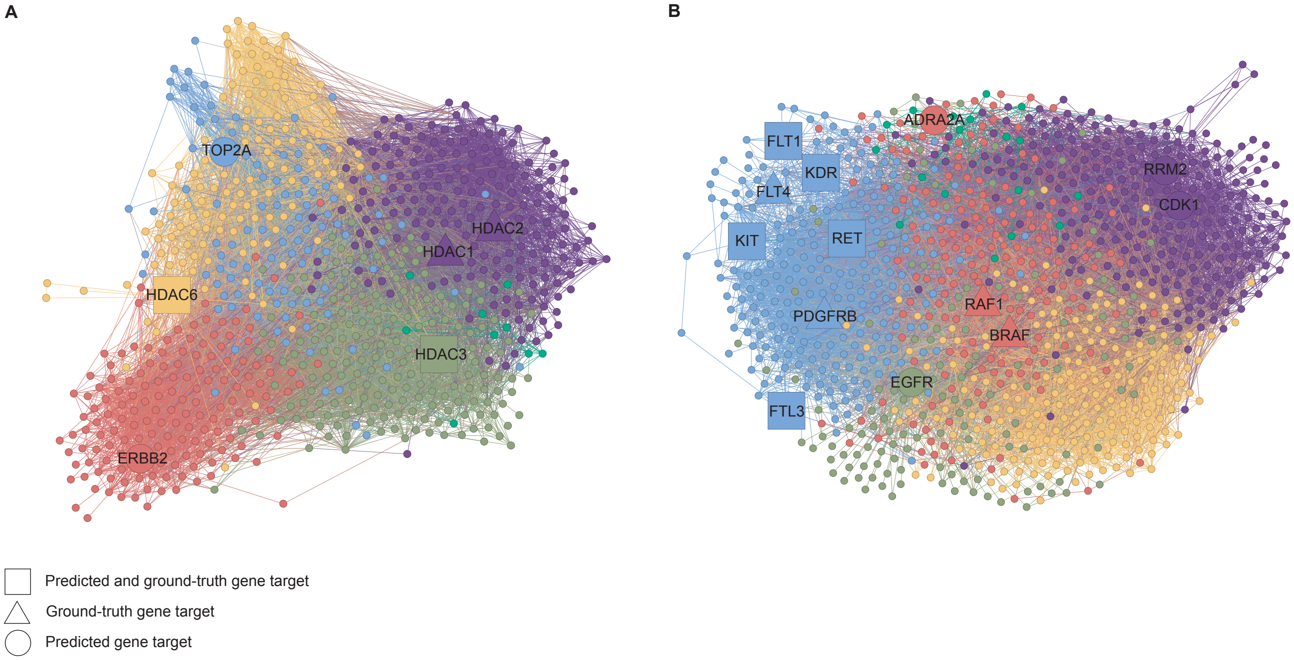


**Figure S3: Network visualization of predicted and ground-truth targets can aid in ellucidation of mechanism of action of drugs. (A,B)** We visualize ground-truth, and predicted therapeutic targets for Vorinostat (A) and Sorafenib (B) in Chemical-PPI-Lung-A549 using Gephi with ForceAtlas embedding. We highlight in different colors distinct communities identified by Gephi's modularity algorithm.

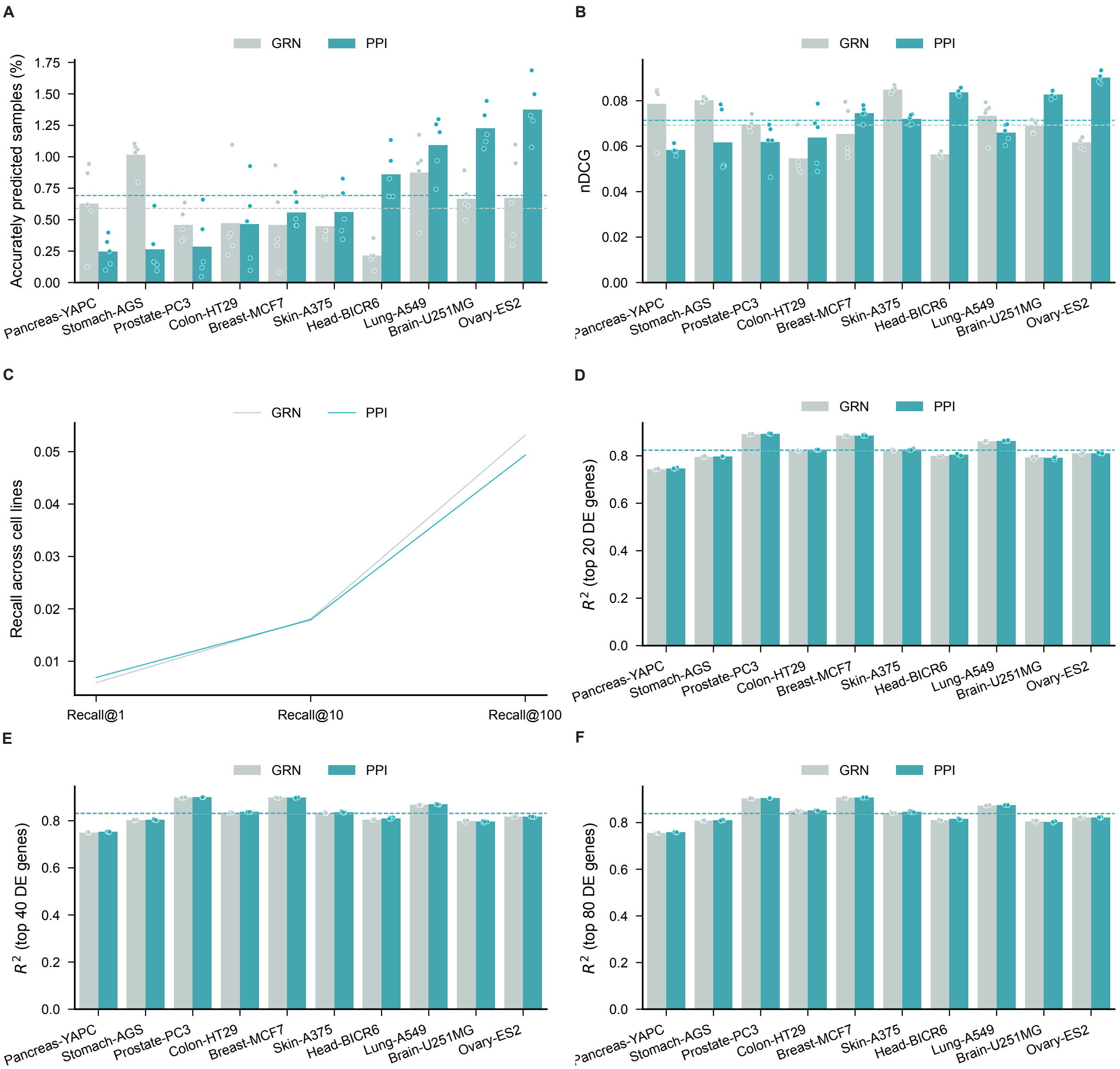


**Figure S4: PDGrapher predicts genetic perturbagens to shift cells from diseased to treated states and predicts response gene expression to genetic perturbagens using cell type specific GRNs. (A)** PDGrapher using cell type specific GRN (PDGrapher-GRN) provides accurate predictions comparable to PDGrapher using cell type agnostic PPI (PDGrapher-PPI) in cell lines with genetic perturbation. **(B)** PDGrapher-GRN ranks shows nDCG comparable to PDGrapher-PPI. **(C)** Both PDGrapher-PPI and PDGrapher-GRN recover ground-truth therapeutic targets at comparable rates compared to competing methods for genetic datasets. **(D-F)** Shown is the $R^2$ of the response prediction module of PDGrapher-GRN and PDGrapher-PPI for the top 20 (D), 40 (E), and 80 (F) differentially expressed (DE) genes. Statistical tests were conducted between PDGrapher and each competing method for each cell line (n = 5), and the corresponding P-values are provided in the Source Data.

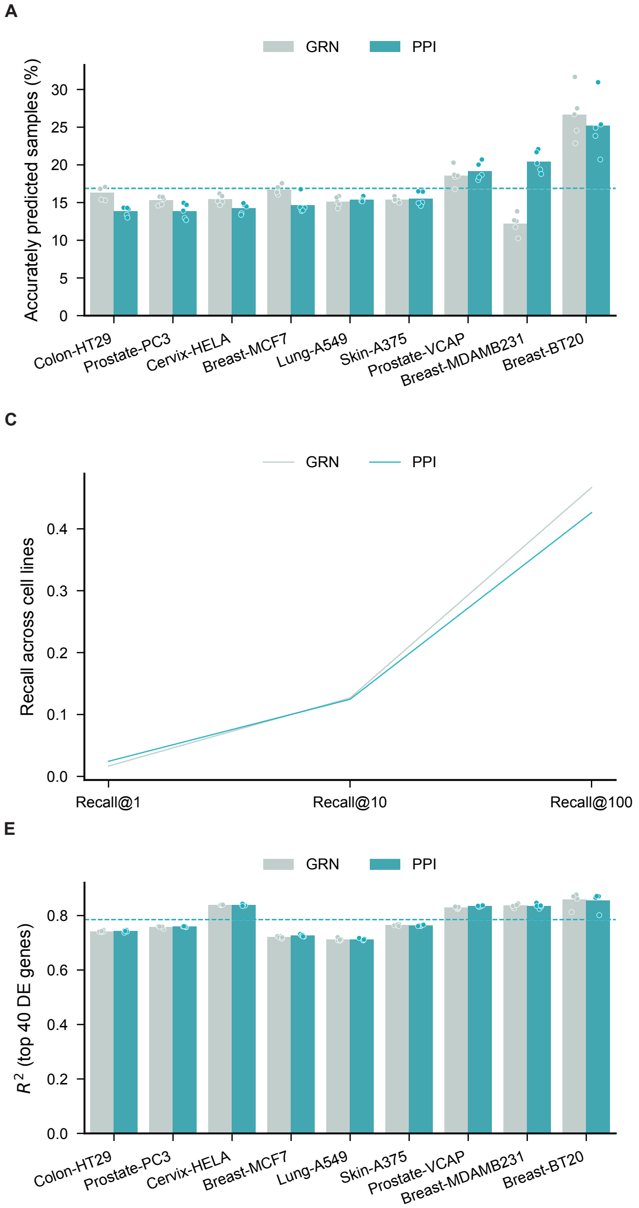


**Figure S5: PDGrapher predicts <u>chemical</u> perturbagens to shift cells from diseased to treated states and predicts response gene expression to <u>chemical</u> perturbagens using cell type specific GRNs.** **(A)** PDGrapher using cell type specific GRN (PDGrapher-GRN) provides accurate predictions comparable to PDGrapher using cell type agnostic PPI (PDGrapher-PPI) in cell lines with chemical perturbation. **(B)** PDGrapher-GRN shows nDCG comparable to PDGrapher-PPI. **(C)** Both PDGrapher-PPI and PDGrapher-GRN recover ground-truth therapeutic targets at comparable rates for chemical datasets. **(D-F)** Shown is the $R^2$ of the response prediction module of PDGrapher-GRN and PDGrapher-PPI for the top 20 (D), 40 (E), and 80 (F) differentially expressed (DE) genes. Statistical tests were conducted between PDGrapher and each competing method for each cell line (n = 5), and the corresponding P-values are provided in the Source Data.

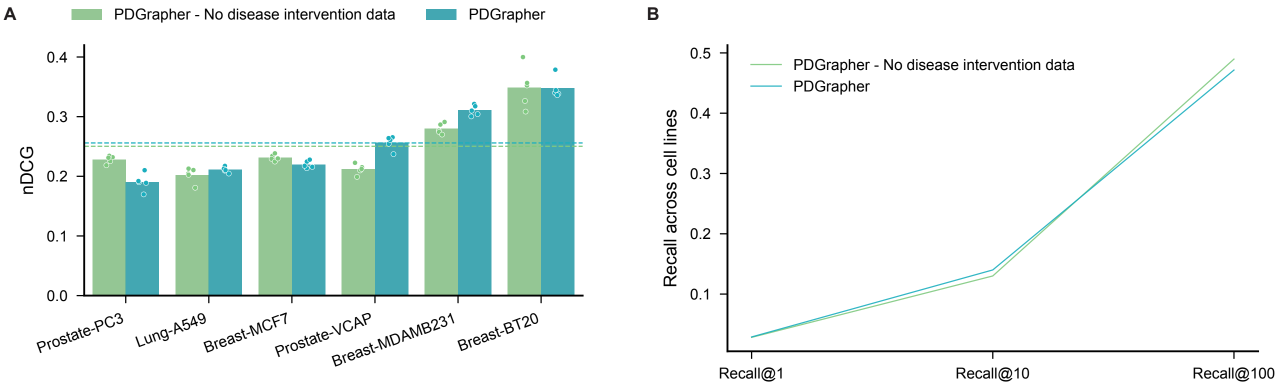


**Figure S6: PDGrapher predicts chemical perturbagens to shift cells from diseased to treated states without using disease intervention data. (A)** PDGrapher (no disease intervention data) shows nDCG comparable to PDGrapher. **(B)** Both PDGrapher and PDGrapher (no disease intervention data) recover ground-truth therapeutic targets at comparable rates for chemical datasets. The results of percentage of accurately predicted samples are shown in Figure 5D. Statistical tests were conducted between PDGrapher and each competing method for each cell line (n = 5), and the corresponding P-values are provided in the Source Data.

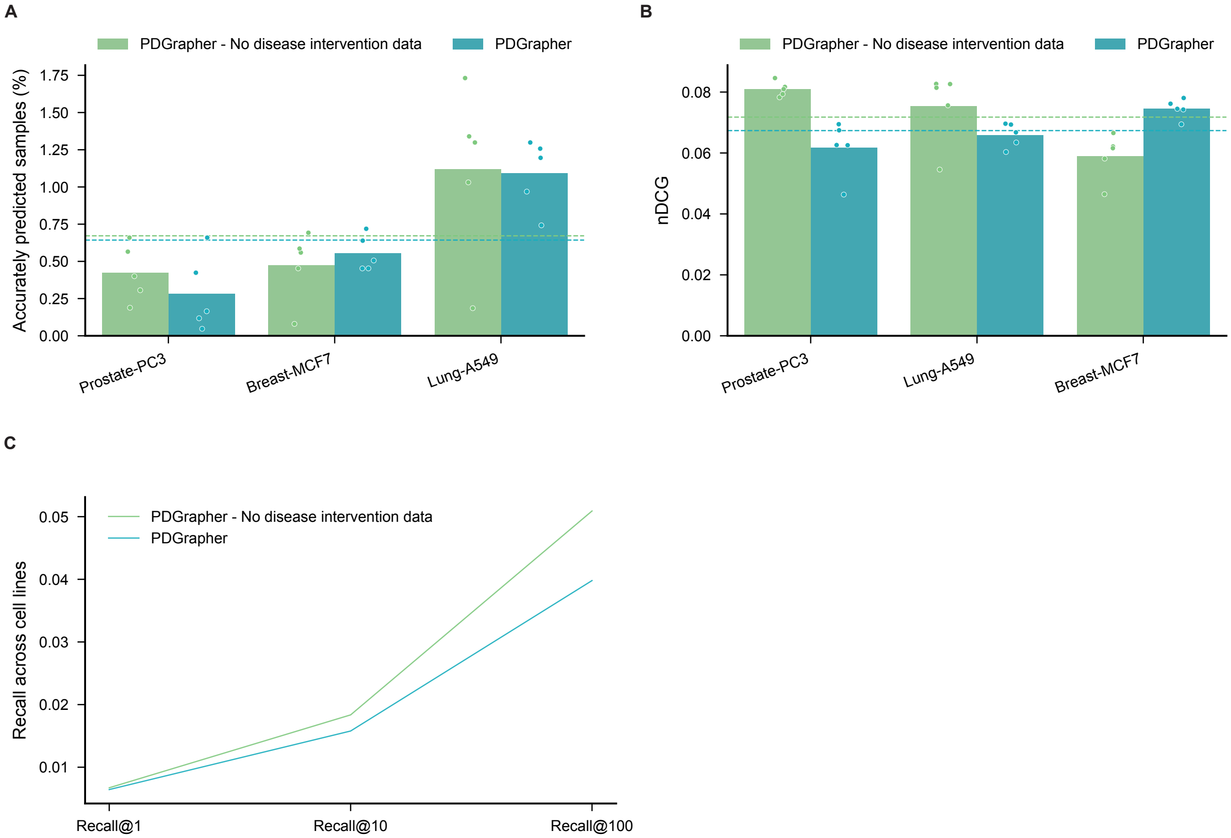


**Figure S7: PDGrapher predicts genetic perturbagens to shift cells from diseased to treated states without using disease intervention data.** **(A)** Performance metrics of the second ablation study on PDGrapher's input data: PDGrapher-No forward data using only treatment intervention data and PDGrapher using both disease and treatment intervention data. The disease and treatment intervention data are organized as (healthy, mutation, diseased) and (diseased, perturbagen, treated), respectively. **(B)** PDGrapher (no disease intervention data) shows nDCG comparable to PDGrapher. **(C)** Both PDGrapher and PDGrapher (no disease intervention data) recover ground-truth therapeutic targets at comparable rates for genetic datasets. Statistical tests were conducted between PDGrapher and each competing method for each cell line (n = 5), and the corresponding P-values are provided in the Source Data.

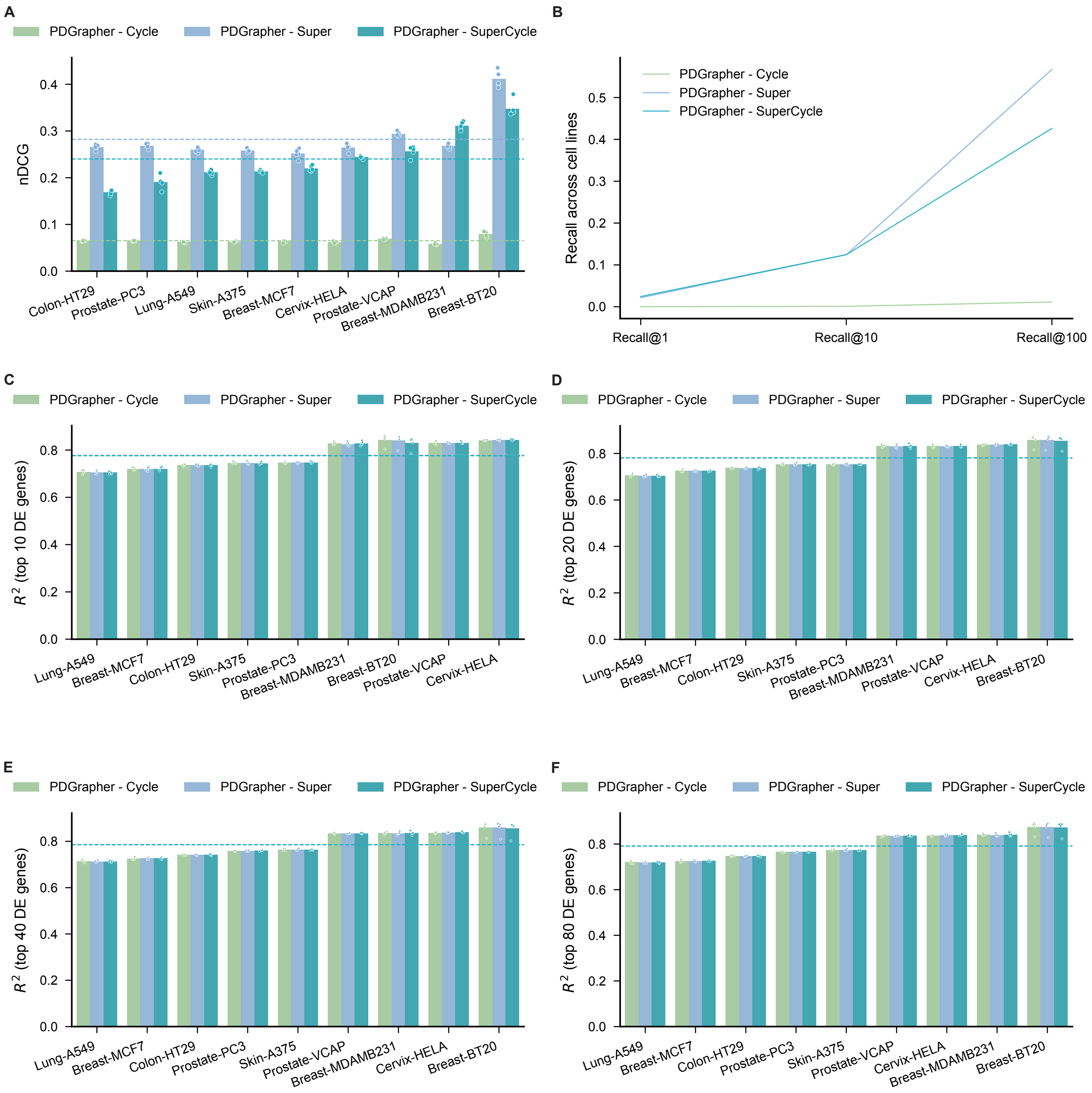


**Figure S8: Ablation studies for loss functions of PDGrapher.** PDGrapher-Cycle trained using only the cycle loss, PDGrapher-Super trained using only the supervision loss, and PDGrapher-SuperCycle trained using both supervision and cycle loss are compared in this ablation study evaluated by nDCG (**A**), recalls (**B**), and $R^2$ of response prediction (C-F). The results of percentage of accurately predicted samples are shown in Figure 5C. Statistical tests were conducted between PDGrapher and each competing method for each cell line (n = 5), and the corresponding P-values are provided in the Source Data.

#### Supplementary references


1. Bastian, M., Heymann, S. & Jacomy, M. Gephi: An open source software for exploring and manipulating networks. BT - International AAAI Conference on Weblogs and Social. *International AAAI Conference on Weblogs and Social Media* 361–362 (2009).

2. Oughtred, R. *et al.* The BioGRID interaction database: 2019 update. *Nucleic Acids Research* **47**, D529–D541 (2019).

3. Luck, K. *et al.* A reference map of the human binary protein interactome. *Nature 2020 580:7803* **580**, 402–408 (2020).

4. Menche, J. *et al.* Uncovering disease-disease relationships through the incomplete interactome. *Science* **347**, 841 (2015).

5. Maglott, D., Ostell, J., Pruitt, K. D. & Tatusova, T. Entrez Gene: gene-centered information at NCBI. *Nucleic Acids Research* **35**, D26 (2007).

6. Tweedie, S. *et al.* Genenames.org: the HGNC and VGNC resources in 2021. *Nucleic Acids Research* **49**, D939–D946 (2021).

7. Cunningham, F. *et al.* Ensembl 2022. *Database issue Nucleic Acids Research* **50**, 989 (2022).

8. Stathias, V. *et al.* LINCS Data Portal 2.0: next generation access point for perturbation-response signatures. *Nucleic Acids Research* **48**, D431–D439 (2020).

9. Huynh-Thu, V. A., Irrthum, A., Wehenkel, L. & Geurts, P. Inferring Regulatory Networks from Expression Data Using Tree-Based Methods. *PLOS ONE* **5**, e12776 (2010).

10. Greenfield, A., Madar, A., Ostrer, H. & Bonneau, R. DREAM4: Combining Genetic and Dynamic Information to Identify Biological Networks and Dynamical Models. *PLoS ONE* **5** (2010).

11. Gentleman, R. C. *et al.* Bioconductor: open software development for computational biology and bioinformatics. *Genome Biology 2004 5:10* **5**, 1–16 (2004).

12. Aibar, S. *et al.* SCENIC: single-cell regulatory network inference and clustering. *Nature Methods 2017 14:11* **14**, 1083–1086 (2017).

13. Seçilmiş, D. *et al.* Knowledge of the perturbation design is essential for accurate gene regulatory network inference. *Scientific Reports* **12**, 1–12 (2022).

14. Qin, R. *et al.* Analysis of oxidase activity and transcriptomic changes related to cutting propagation of hybrid larch. *Scientific Reports* **13**, 1–12 (2023).

15. Song, Q., Ruffalo, M. & Bar-Joseph, Z. Using single cell atlas data to reconstruct regulatory networks. *Nucleic Acids Research* 1–13 (2023).

16. Tate, J. G. *et al.* COSMIC: the Catalogue Of Somatic Mutations In Cancer. *Nucleic Acids Research* **47**, D941–D947 (2019).

17. Wishart, D. S. *et al.* DrugBank 5.0: a major update to the DrugBank database for 2018. *Nucleic Acids Research* **46**, D1074–D1082 (2018).

18. Heller, S. R., McNaught, A., Pletnev, I., Stein, S. & Tchekhovskoi, D. InChI, the IUPAC International Chemical Identifier. *Journal of Cheminformatics* **7**, 1–34 (2015).

19. Mueller, J., Reshef, D. N., Du, G. & Jaakkola, T. Learning Optimal Interventions (2016).

20. Pacchiano, A. & Barton, R. A. Neural Design for Genetic Perturbation Experiments 1–37.

21. Mueller, J., Gifford, D. & Jaakkola, T. Sequence to Better Sequence : Continuous Revision of Combinatorial Structures (2017).

22. Hie, B., Bryson, B. D., Zhong, E. D. & Berger, B. Learning Mutational Semantics 1–13 (2020).

23. Zhang, J., Squires, C. & Uhler, C. Matching a Desired Causal State via Shift Interventions (2021).

24. Zhang, J., Cammarata, L., Squires, C., Sapsis, T. P. & Uhler, C. Active Learning for Optimal Intervention Design in Causal Models (2022).

25. Deng, Z., Zheng, X., Tian, H. U. & Zeng, D. D. Deep Causal Learning: Representation, Discovery and Inference .

26. Parafita, & Vitrià, J. Causal Inference with Deep Causal Graphs (2020).

27. Pawlowski, N., Castro, D. C. & Glocker, B. Deep structural causal models for tractable counterfactual inference. *Advances in Neural Information Processing Systems* **2020-Decem** (2020).

28. Xia, K., Lee, K.-Z., Bengio, Y. & Bareinboim, E. The Causal-Neural Connection: Expressiveness, Learnability, and Inference .

29. Xia, K., Pan, Y. & Bareinboim, E. Neural Causal Models for Counterfactual Identification and Estimation **2**, 1–57 (2022).

30. Bronstein, M. M., Bruna, J., LeCun, Y., Szlam, A. & Vandergheynst, P. Geometric deep learning: Going beyond Euclidean data. *IEEE Signal Processing Magazine* **34**, 18–42 (2017).

31. Bronstein, M. M., Bruna, J., Cohen, T. & Veličković, P. Geometric Deep Learning: Grids, Groups, Graphs, Geodesics, and Gauges (2021).

32. Li, M. M., Huang, K. & Zitnik, M. Graph representation learning in biomedicine and healthcare. *Nature Biomedical Engineering* **6**, 1353–1369 (2022).

33. Zečević, M., Dhami, D. S., Veličković, P. & Kersting, K. Relating graph neural networks to structural causal models. *arXiv:2109.04173* (2021).

34. Zečević, M., Dhami, D. S., Veličković, P. & Kersting, K. Relating Graph Neural Networks to Structural Causal Models (2021).
