## Extended data figures for "Combinatorial prediction of therapeutic perturbations using causally-inspired neural networks"

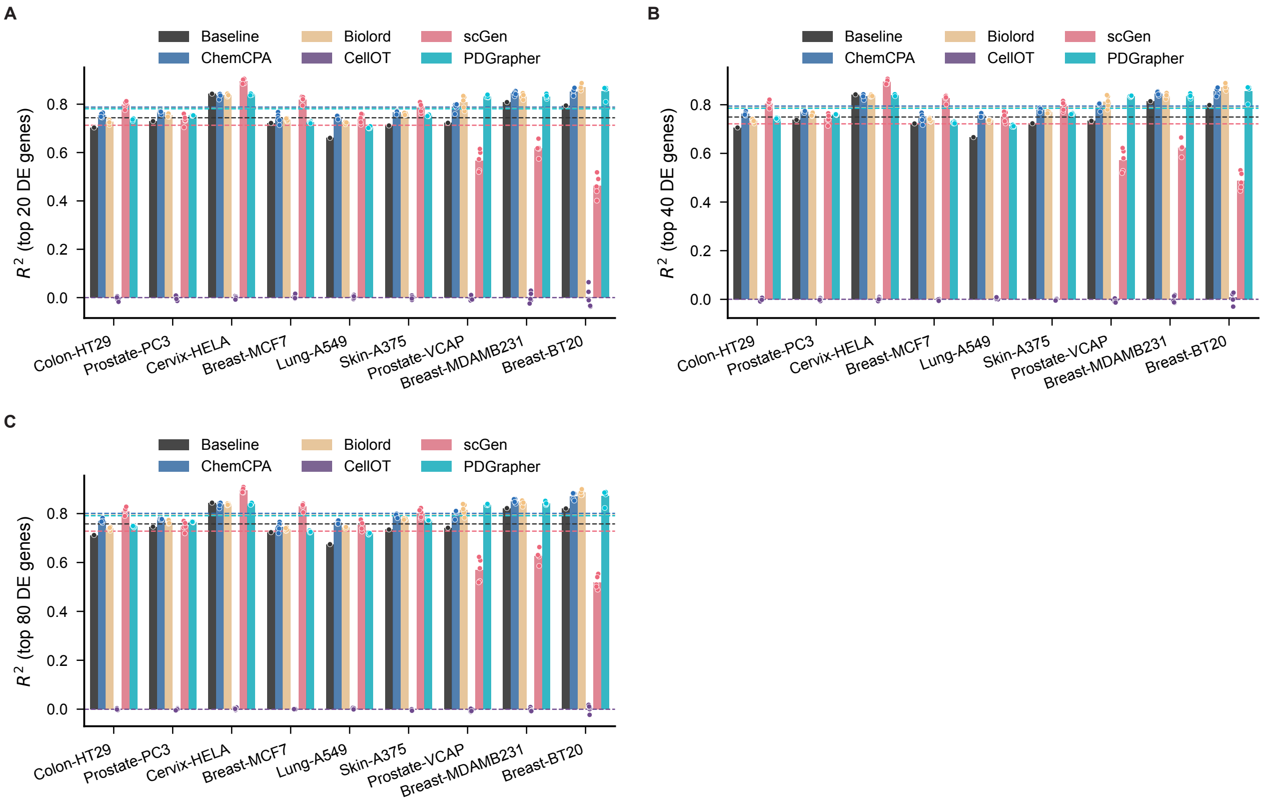


**Extended Data Fig. 1: The performance of response prediction within nine cell lines under chemical perturbation.** The $R^2$ values are calculated between the predicted and actual gene expression for the top 20 (**A**), 40 (**B**), and 80 (**C**) differentially expressed genes per cell line. Dotted lines represent the average performance across cell lines, dots indicate individual data points, and bars represent the average $R^2$ across five data splits. P-values from the statistical tests are provided in the Source Data.

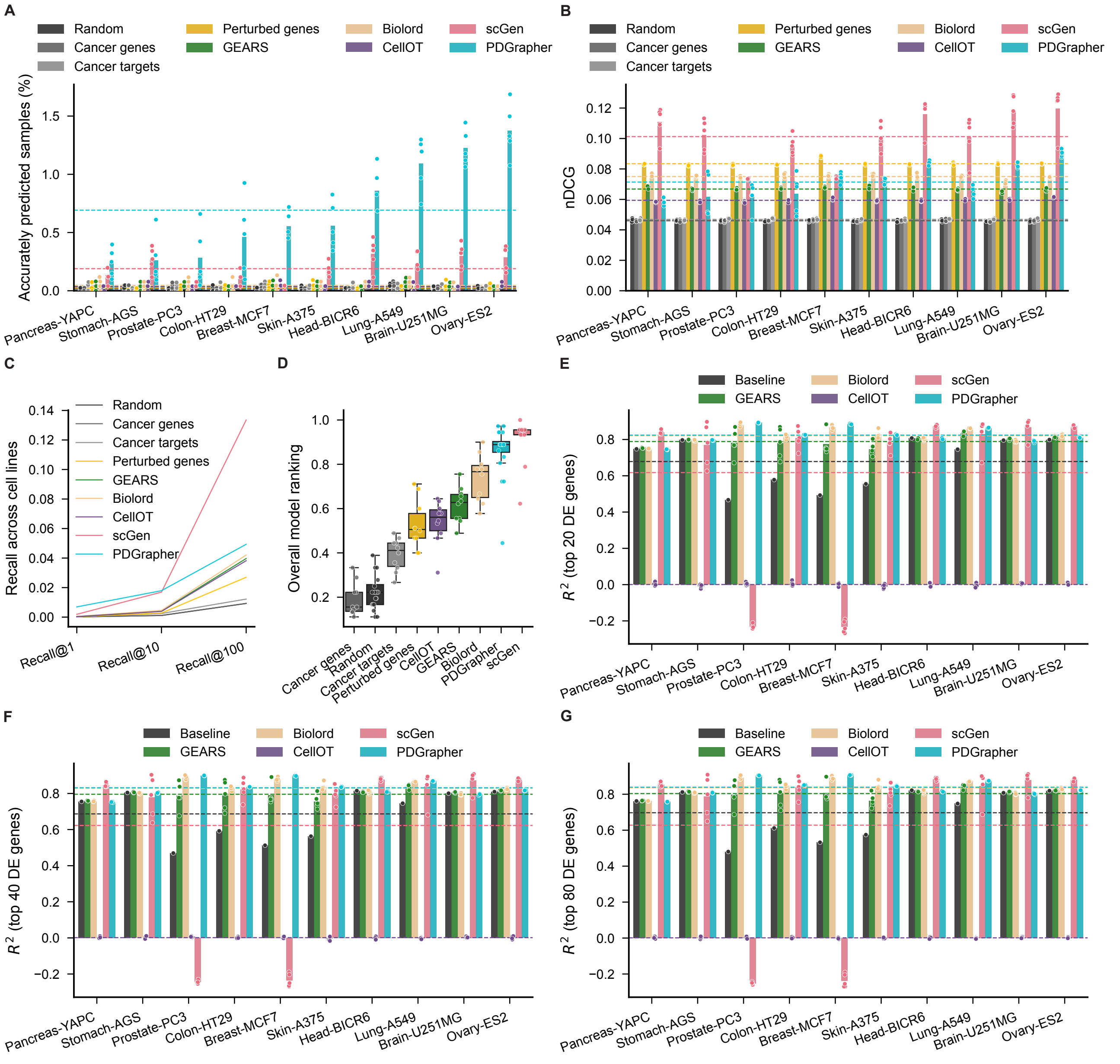


**Extended Data Fig. 2: PDGrapher efficiently predicts genetic perturbagens to shift cells from diseased to treated states in a random splitting setting within ten cell lines.** **(A)** PDGrapher provides accurate predictions for up to 1.09% (Genetic-PPI-Ovary-ES2: 1.37% vs 0.28%) more samples in the test set compared to the second-best baseline across Genetic-PPI datasets **(B)** scGen takes the leading position in nDCG across genetic Genetic-PPI datasets. **(C)** PDGrapher recovers ground-truth therapeutic targets at comparable rates compared to competing methods for Genetic-PPI datasets. **(D)** PDGrapher has the best overall performance in perturbagen prediction within each cell line evaluated by the averaged rank over multiple cell lines and metrics. **(E-G)** Shown is the $R^2$ of the response prediction module of PDGrapher compared to competing baselines for the top 20 (E), 40 (F), and 80 (G) differentially expressed (DE) genes. The central line inside the box represents the median, while the top and bottom edges correspond to the first (Q1) and third (Q3) quartiles. The whiskers extend to the smallest and largest values within 1.5 times the interquartile range (IQR) from the quartiles. P-values from the statistical tests are provided in the Source Data.

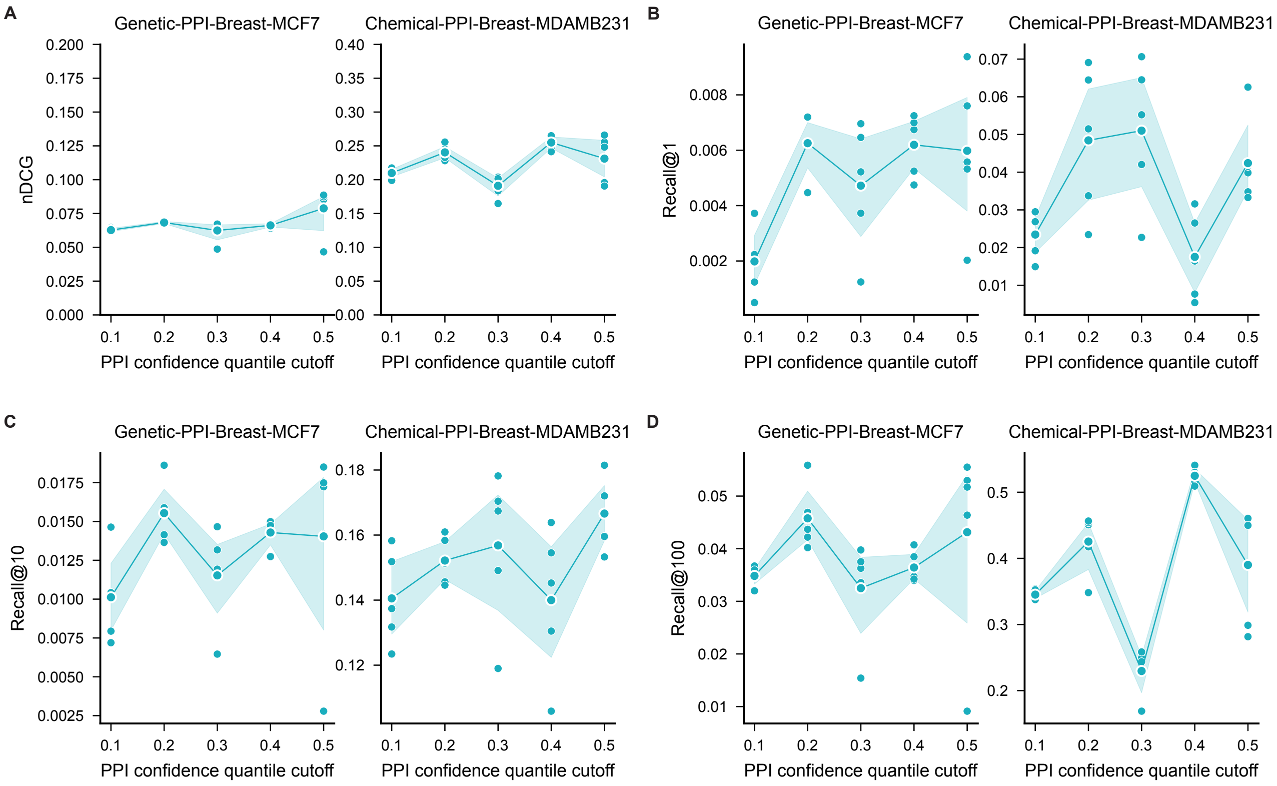


**Extended Data Fig. 3: Sensitivity analysis of the PPI used for training PDGrapher.** Performance of sensitivity analyses evaluated by nDCG (**A**) and recalls (**B-D**) for datasets Genetic-PPI-Breast-MCF7 (left) and Chemical-PPI-Breast-MDAMB231 (right). The PPI used here is from STRING (string-db.org), which includes a confidence score for each edge. The edges are filtered by the 0.1, 0.2, 0.3, 0.4, and 0.5 quantiles of the confidence scores as cutoffs, resulting in five PPI networks with 625,818, 582,305, 516,683, 443,051, and 296,451 edges, respectively. The results of percentage of accurately predicted samples are shown in Figure 5B.

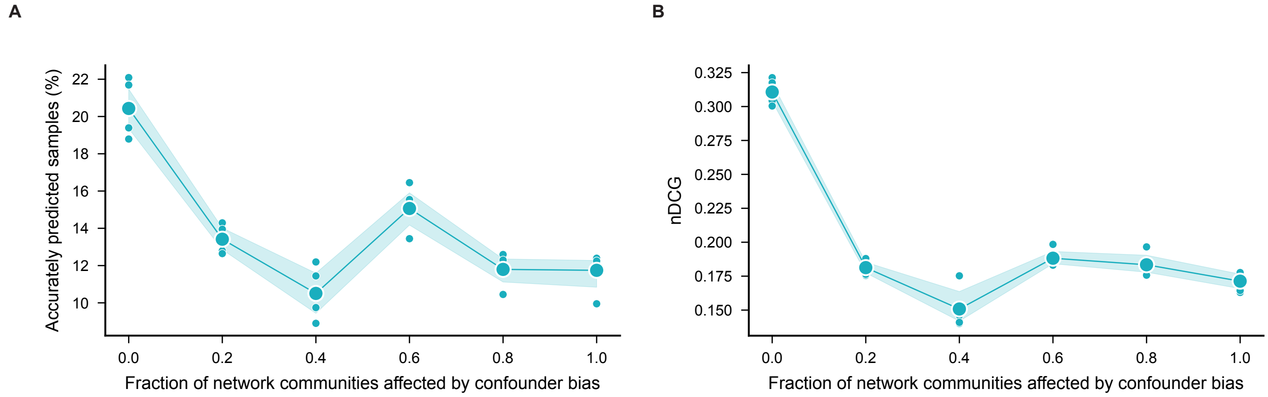


**Extended Data Fig. 4: PDGrapher has stable performance on the synthetic datasets with various intensities of confounders added on the gene expression.** Performance of simulation analyses evaluated by percentage of accurately predicted samples **(A)** and nDCG **(B)** for the synthetic datasets with with varying levels (0 to 1) of confounding bias introduced into the gene expression data. Gaussian noise, with distinct means and variances, was added progressively to random subsets of genes, simulating latent confounder effects in the treated gene expression data. The intensity of the confounding bias increases as more gene groups (representing network communities) are affected. This approach creates global, controlled variability in the gene expression data, paired with an unperturbed PPI network, allowing for the evaluation of algorithmic performance across different degrees of confounder noise. See Online Methods for more details on data generation.

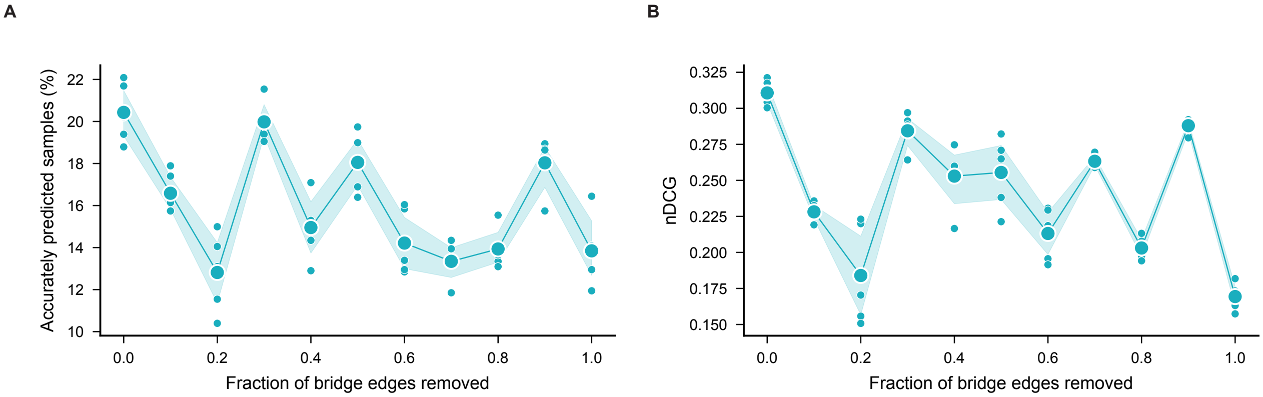


**Extended Data Fig. 5: PDGrapher has stable performance on the synthetic datasets with various fractions of bridge edges removed.** Performance of simulation analyses evaluated by percentage of accurately predicted samples **(A)** and nDCG **(B)** for the synthetic datasets with a [0, 0.1, ..., 1] fraction of bridge edges removed in the simulated PPI. Bridge edges are those with high connectivity in the network which, if removed, increase the number of disconnected communities. The number of connected components in the network upon bridge edge removal are [90, 179, 268, 358, 447, 536, 626, 715, 804, 894]. See more information in Table S5.

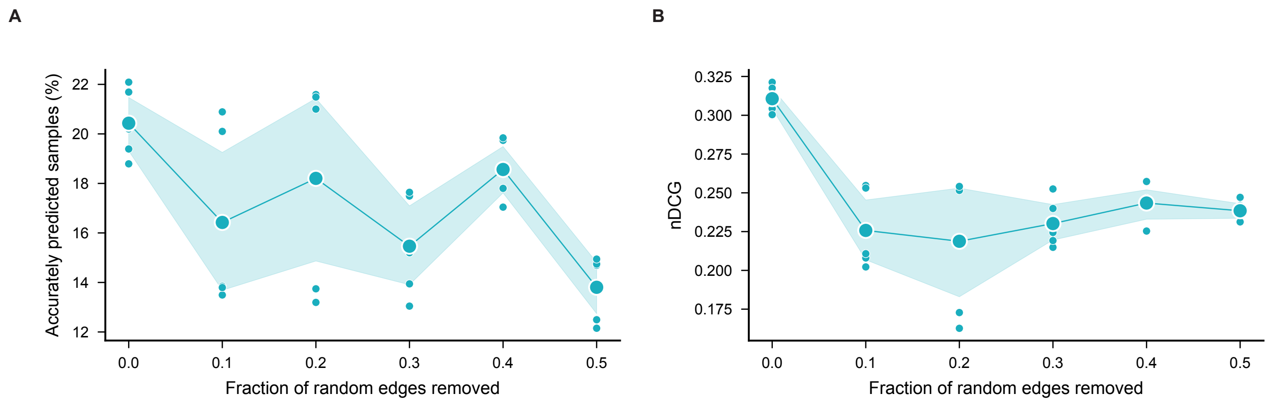


**Extended Data Fig. 6: PDGrapher's performance is influenced by network incompleteness.** Performance in ablation studies evaluated by percentage of accurately predicted samples **(A)** and nDCG **(B)** for the synthetic datasets with a [0, 0.1, … 0.6] fraction of random edges removed in the PPI. The number of remaining edges in the network upon random edge removal are [273,319; 242,912; 212,525; 182,177; 151,811; 121,472]. See the Methods section for more details.

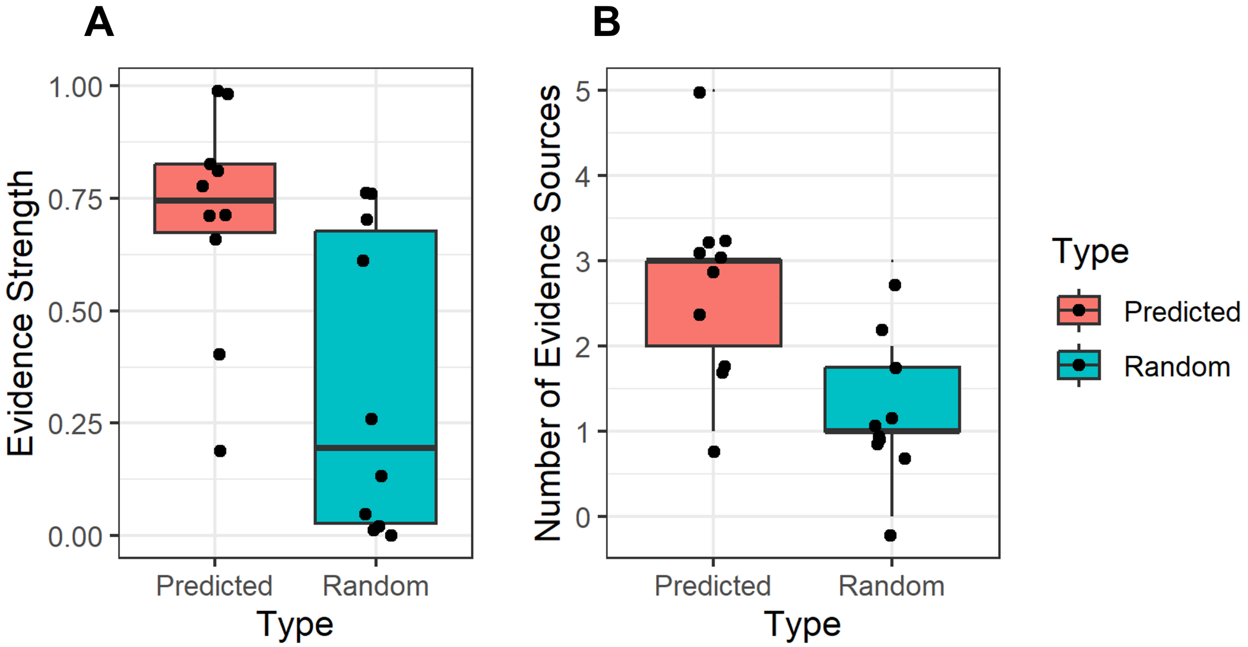


**Extended Data Fig. 7: Comparison of predicted targets from PDGrapher and a random model for lung cancer.** Boxplots display the evidence strength (A) and the number of evidence sources (B) for the top 10 predicted targets from PDGrapher versus 10 randomly selected genes. The central line inside the box represents the median, while the top and bottom edges correspond to the first (Q1) and third (Q3) quartiles. The whiskers extend to the smallest and largest values within 1.5 times the interquartile range (IQR) from the quartiles. P-values from the statistical tests are provided in the Source Data.

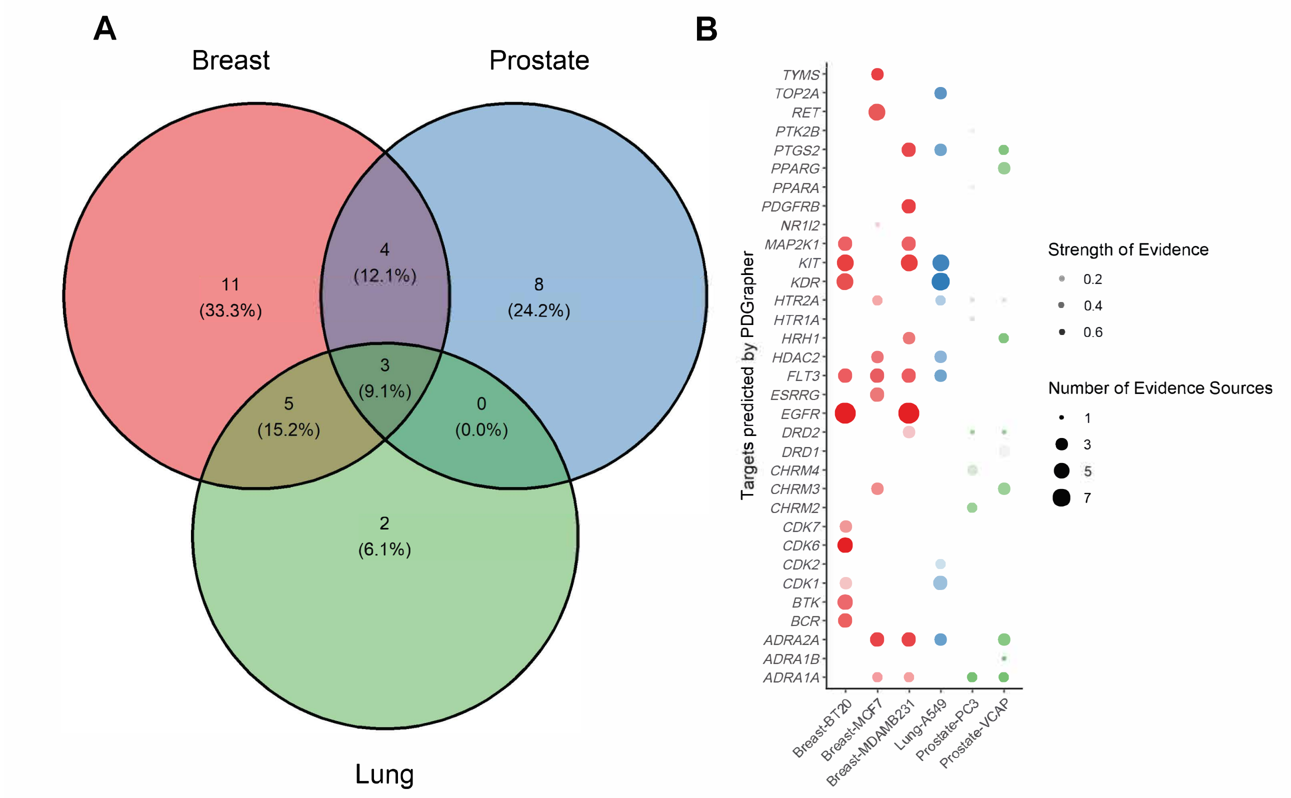


**Extended Data Fig. 8: Unique and common targets predicted by PDGrapher among three cancer types.** The Venn diagram (A) shows the number and ratio of unique and common predicted targets, while the bubble plot (B) shows the strength of evidence and the number of evidence sources for each cell line. The evidence strengths are the global association scores and overall evidence sources provided by OpenTargets. Details of the scoring system are in Supplementary Note 2.

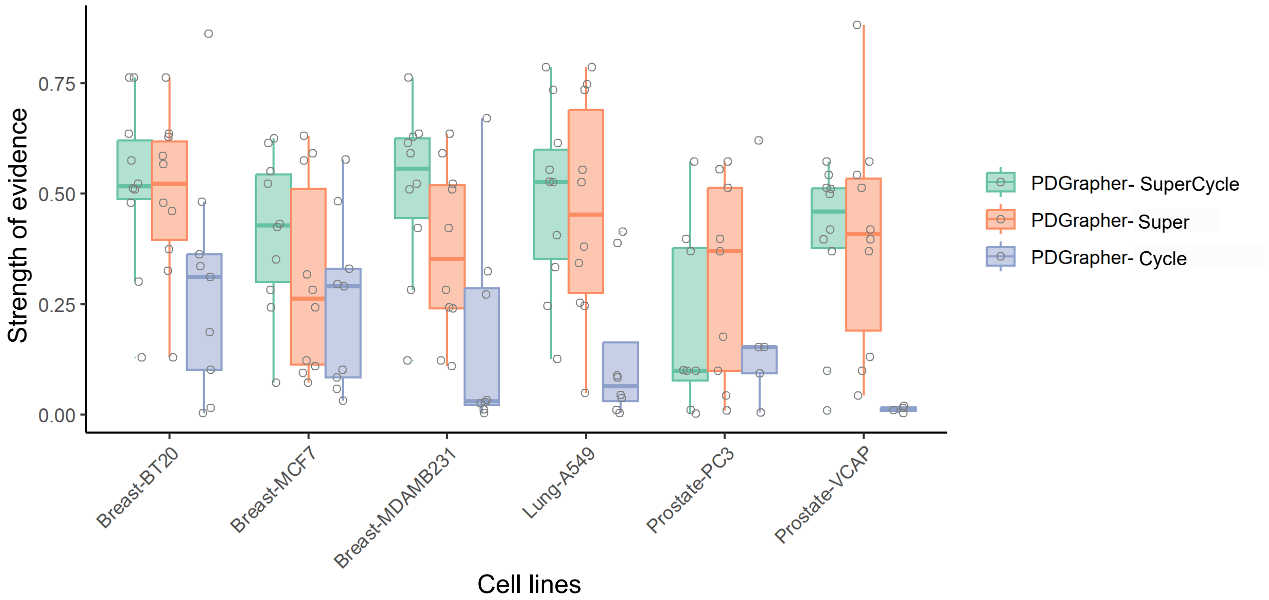


**Extended Data Fig. 9: Ablation studies for loss functions of PDGrapher evaluated by the evidence from OpenTargets.** The strength of evidence in the cell lines with healthy control data is shown in the box plots. The strength of the evidence is the global association scores, which are based on all evidence sources provided by OpenTargets. See details of the scoring system in Supplementary Note 2. The central line inside the box represents the median, while the top and bottom edges correspond to the first (Q1) and third (Q3) quartiles. The whiskers extend to the smallest and largest values within 1.5 times the interquartile range (IQR) from the quartiles. P-values from the statistical tests are provided in the Source Data.
